# Warmer water temperature and epizootic shell disease reduces diversity but increases cultivability of bacteria on the shells of American Lobster (*Homarus americanus*)

**DOI:** 10.1101/2022.11.11.512360

**Authors:** Suzanne L. Ishaq, Sarah M. Turner, Grace Lee, M. Scarlett Tudor, Jean D. MacRae, Heather Hamlin, Deborah Bouchard

## Abstract

The American lobster, *Homarus americanus*, is an economically valuable and ecologically important crustacean along the North Atlantic coast of North America. Populations in southern locations have declined in recent decades due to increasing ocean temperatures and disease, and these circumstances are progressing northward. We monitored 57 adult female lobsters, healthy and shell-diseased, under three seasonal temperature cycles for a year, to track shell bacterial communities using culturing and 16S rRNA gene sequencing, progression of ESD using visual assessment, and antimicrobial activity of hemolymph. The richness of bacterial taxa present, evenness of abundance, and community similarity between lobsters was affected by water temperature at the time of sampling, water temperature over time based on seasonal temperature regimes, shell disease severity, and molt stage. Several bacteria were prevalent on healthy lobster shells but missing or less abundant on diseased shells, although putative pathogens were found on all shells regardless of health status.

**Graphical abstract:** 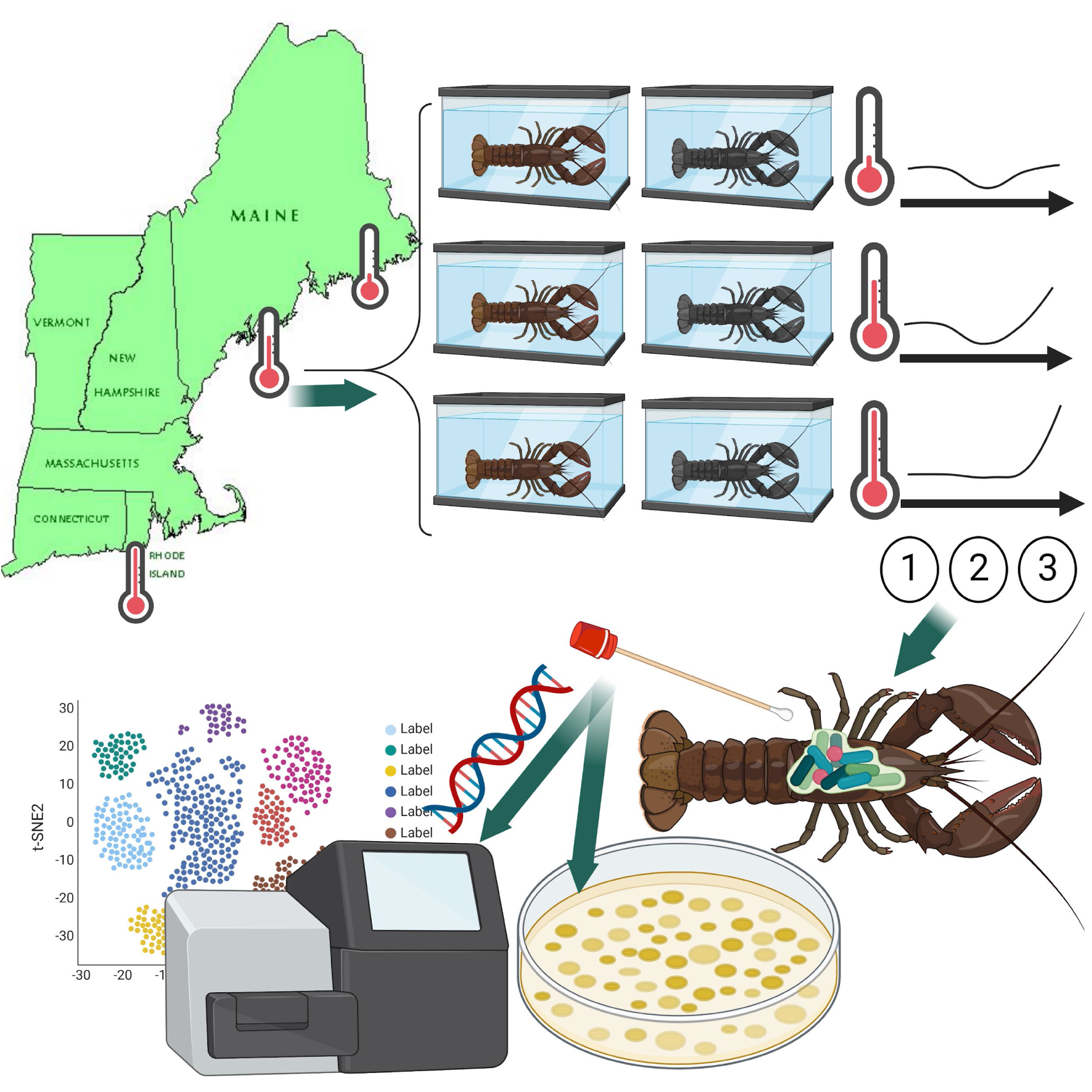

## Introduction

American lobsters, *Homarus americanus* (family Nephropidae), are bottom-feeding crustaceans found along the cold coastal waters in the Northwestern Atlantic Ocean, where they scavenge detritus or prey on other crustaceans, shellfish, or small fish. Lobsters have a hard carapace, which they molt regularly to grow during their entire life, and can live upwards of 100 years before their size, disease, or shell deformities makes molting too physically and energetically difficult to survive (Bowser and Rosemark, 1981; Groner et al., 2018). They travel over areas of several miles, and migrate to find cooler waters or a more hospitable location (Campbell, 1986; Watson, 2005). Their muscular tail and large, unequally-sized front claws (Aiken, 1980) historically and currently makes them a food staple for local peoples, with lobstering and lobster consumption contributing to local cultural identities. Lobster meat is a popular and highly valued delicacy for locals and tourists (Daniel et al., 2008; Steneck et al., 2011), and landings represent the most economically important fishery in the northwestern Atlantic Ocean (Maine Department of Marine Resources, 2021).

While currently the fishery remains viable and healthy in northern New England, lobster populations in southern New England (SNE) collapsed more than a decade ago, with no sign of recovery (Wahle et al., 2015). Temperatures in SNE appear to have surpassed the lobsters’ thermal tolerance, 22℃, not only in highest temperature but also the number of days over 20°C (Glenn and Pugh, 2006; Pinsky and Fogarty, 2012; Wahle et al., 2015). High temperatures and other stressors can result in physiological signs (Lorenzon et al., 2007), which can impact lobsters’ homeostasis and ability to molt or fight infections (Dove et al., 2005). Lobster population declines in SNE regions have also been attributed to ocean acidification through carbon dioxide deposition (Niemisto et al., 2021), pollution and anthropogenic disturbance (Goodman et al., 2021), and disease emergence (Harrington et al., 2020; Shields et al., 2012). Since then, population declines have been noted in progressively-more-northern fisheries, and there is concern that warmer ocean temperatures which are already pushing lobsters further north to cooler waters (Greenhalgh, 2016) may also push infectious disease to new locations (Ishaq et al., 2022; Reardon et al., 2018; Shields, 2019; Watson, 2005). In particular, the Gulf of Maine is warming faster than predicted (Poppick, 2018), and there is a concern that lobster fisheries could collapse in the way that other species, such as Atlantic cod (*Gadus morhua*), already have (Pershing et al., 2015).

Epizootic shell disease (ESD) emerged in the later part of 1996 in SNE and by 2000 was spreading along the Northeastern coast. This new, aggressive form of shell disease was described as a ‘severe erosive shell disease affecting the dorsal carapace of *H. americanus*’ (Smolowitz et al., 2005). ESD is recognized as a significant disease syndrome and is described as having a multifactorial and complex etiology with increased temperatures implicated as a contributing factor (Castro et al., 2012; Shields et al., 2012). The histological characteristics of the disease suggested a bacterial etiology but a causative agent has yet to be determined (Ishaq et al., 2022). Culture-dependent and culture-independent bacterial investigations have been performed on shell-diseased lobsters and shown that the etiology of the disease is complex, likely involving multiple bacterial species in the Alteromonadaceae, Flavobacteriaceae, Pseudomonadaceae, Rhodobacteraceae and Vibrionaceae families (Bell et al., 2012; Chistoserdov et al., 2012, 2005; Meres et al., 2012). Particular focus has been placed on two microbial genera consistently associated with ESD, *Aquimarina* and *Thalassobius* species (Chistoserdov et al., 2012; Quinn et al., 2017, 2013), but a true etiology has yet to be defined. In fact, the role of these bacteria in disease lesions remain unclear and a dysbiotic shift in the shell microbial community resulting from environmental stressors has been suggested (Feinman et al., 2017; Ishaq et al., 2022; Meres et al., 2012).

While past research focused on identification of microbial communities on wild caught or laboratory housed lobsters, these studies did not follow changes in microbial communities and profiles over time, nor did they examine the effect of colder temperatures as well as warmer ones. Previous work (Bouchard, 2018) and this related study seek to understand how seasonal temperature regimes influence plasma antimicrobial activity and the progression of ESD in adult female American lobsters, in a controlled aquaculture environment. Current literature indicates that ESD is not transferable in laboratory aquaria but that it will continue to progress in the individuals with shell disease.

We hypothesized that more extreme temperature regimes would alter shell bacterial communities, exacerbate ESD progression, and reduce metrics of immune health. Previous research has investigated heat stress in lobsters, but none have included groups treated with colder water than that which they had been acclimated. We monitored 57 adult female lobsters, healthy versus shell-diseased, under three regimes mimicking the seasonal temperatures cycles by geographic location: colder water in Northern Maine (NME), warm water in Southern Maine (SME) where lobsters originated from, and hotter waters in Southern New England (SNE). Samples were collected at three timepoints over a year, to track shell bacterial communities using culturing and 16S rRNA gene sequencing, progression of ESD using visual assessment, and antimicrobial activity of hemolymph.

## Results

### Warmer tank temperature reduced bacterial community richness but increased cultured isolate counts

Bacterial sequence variant (SV) richness on lobster shells (Figure 1A) was highest at the summer baseline timepoint, immediately after lobsters were collected from ocean waters and prior to the implementation of temperature treatments. The high bacterial diversity on shells found at baseline did not persist in tanks over time, reflecting the low biological and chemical diversity in tanks which cannot support complex microbial communities. Across all tanks, richness was lower by an estimated 78 SVs in the winter samples, 4 months after baseline (linear regression model, t = -7.14, *p* < 0.001), as well as 57 SVs lower in the summer samples, 10 months after baseline (linear regression model, t = -5.13, *p* < 0.001).

**Figure 1.**
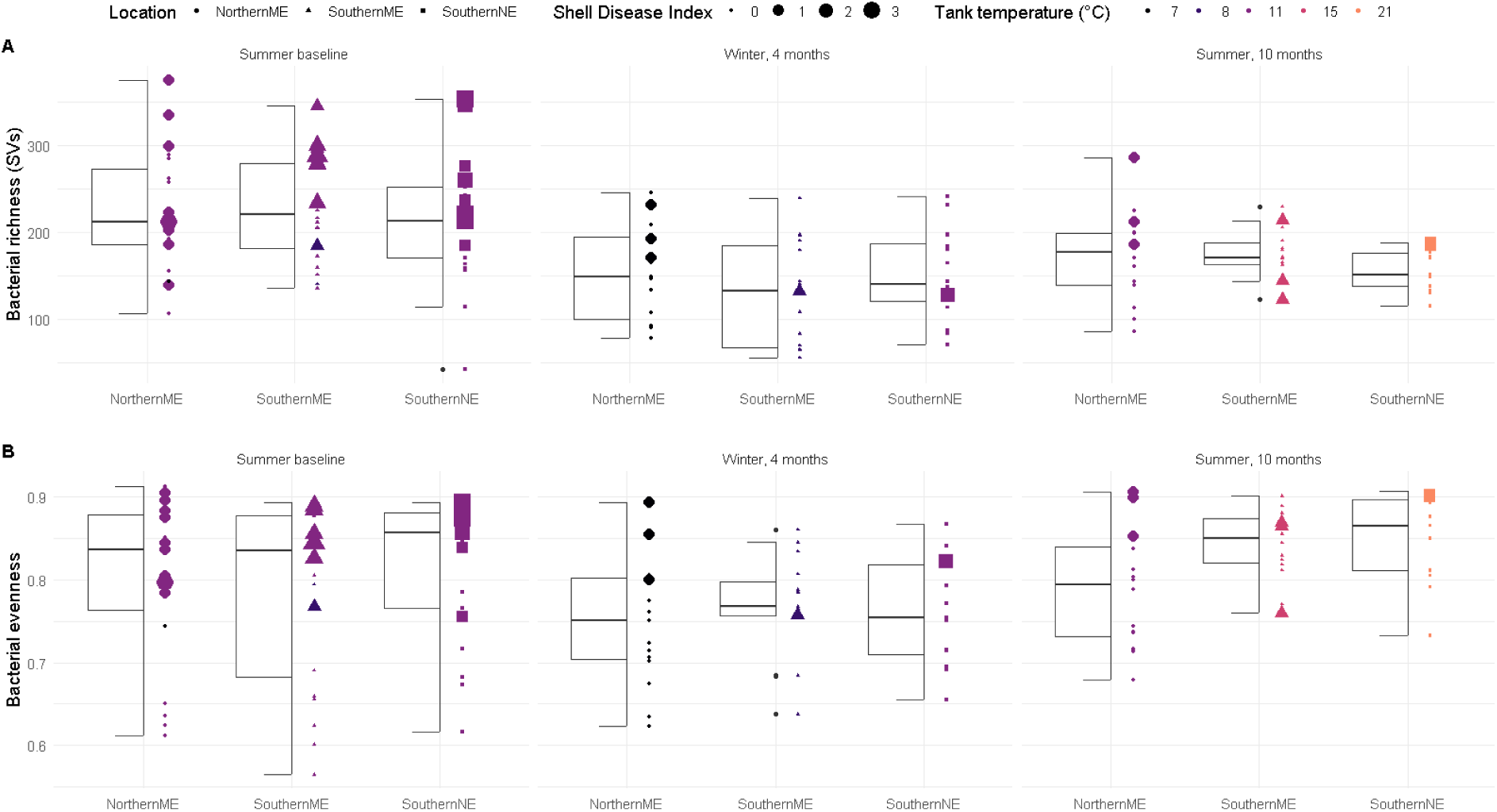
Sequence variant richness (A) and Shannon’s evenness (B) for lobster shell bacterial communities under three simulated seasonal ocean temperatures over time. Average seasonal ocean temperatures were simulated (i.e., tank temperature denoted by point color) for Northern Maine (ME), Southern Maine, and Southern New England (NE). Point size indicates epizootic shell disease stage.

For lobsters in the Northern Maine (NME) tanks, maintained at cooler temperatures, shells had 73 fewer bacterial SVs after 4 months (lm, t = -3.75, *p* < 0.001), and 54 fewer SVs after 10 months (lm, t = -2.86, *p* < 0.005). Lobsters in the Southern Maine (SME) tanks, which simulated the temperature where all lobsters originated but is somewhat warmer than their ideal seasonal temperatures, had 100 fewer bacterial SVs on shells after 4 months as compared to baseline (lm, t = -5.22, *p* < 0.001), and 53 fewer SVs at 10 months as compared to baseline (lm, t = -2.72, *p* < 0.007). For lobsters in the Southern New England (SNE) tanks maintained at higher temperatures, shells had 63 fewer bacterial SVs at 4 months (lm, t = -3.36, *p* < 0.001), and 65 fewer SVs at 10 months compared to baseline (lm, t = -3.33, *p* < 0.002).

The evenness of bacterial SV abundance was likewise highest at the summer baseline timepoint, lower in the winter, and increased again at the next summer timepoint (Figure 1B). Low evenness at cold temperatures and high evenness at high temperature may indicate that only certain bacterial species are affected by cold water, but all/most bacterial taxa are affected by warmer water. Evenness of bacterial SV abundance was increased by tank temperature (lm, increase = 0.01, t = 4.16, *p* < 0.001), and in particular, higher in the SME samples in summer, 10 months post (lm, increase = 0.06, t = 2.03, *p* = 0.04), and lower in the SNE samples in winter, 4 months post (lm, decrease = -0.06, t = -2.21, *p* = 0.029).

Over the three sampling timepoints, a total of 145 carapace samples were processed for microbial enumeration and culture. Bacterial richness in sequencing data was not correlated with bacterial colony counts (Figure 8). At the baseline sampling, in summer, the bacterial counts for lobsters from all three systems were combined as the temperatures were all the same (temp = 12.5℃). Lobsters showing signs of shell disease had significantly more bacteria per cm^2^ than non-shell diseased lobsters (*t* test: *t*_48_ = -1.989; *p* = 0.052; Figure 2A). At the first sampling period 4 months later, winter, for healthy and diseased lobsters, there were significantly more bacteria in the high temperature regimes (SNE) than the low (NME) and mid (SME) temperature regimes (Kruskal-Wallis: non-shell diseased: c^2^ =14.385, df = 2, *p* = 0.001; shell diseased: c^2^ = 7.232, df = 2, *p* = 0.027; Figure 2B). In the low temperature regime, the shell diseased lobsters had significantly more bacteria than the healthy lobsters (Mann-Whitney: low/NME: Z = -2.100, *p* = 0.036, Figure 2B). There was no difference in bacterial counts between healthy and diseased lobsters in the mid and high temperature regime (Mann-Whitney: mid/SME: Z = -0.476, *p* = 0.634; high/SNE: Z = -0.880, *p* = 0.379; Figure 2B).

**Figure 2.**
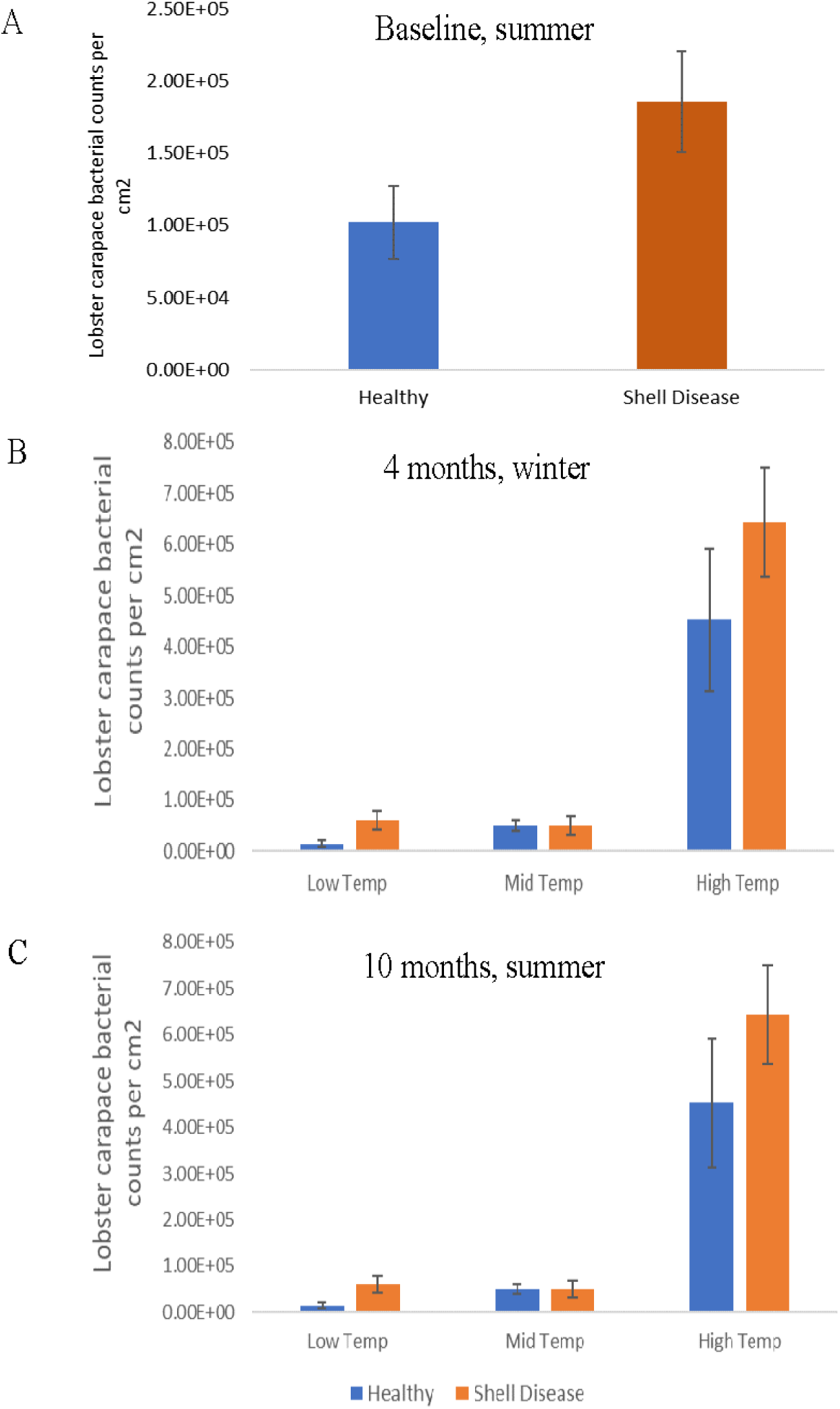
Bacterial counts from lobster shell isolates plated on marine agar plates. A; Baseline, with all three temperature treatment systems at an average temperature of 12.5**°**C. B; Time 1, average system temperatures: Low (NME) = 7.5**°**C, Mid (SME) = 8.5**°**C, and High (SNE) = 11.0**°**C. C; Time 2, average system temperatures: Low = 10**°**C, Mid = 15**°**C, and High = 21.0**°**C. Error bars indicate standard error.

At the second sampling period 10 months after baseline, again in summer, there were significantly more bacteria for both healthy and diseased lobsters. There were significantly more bacteria in the mid and high temperature regimes than the low temperature regimes (Kruskal-Wallis: Healthy: c^2^ =13.412, df = 2, *p* = 0.001; Diseased: c^2^ = 11.498, df = 2, *p* = 0.003; Figure 2C). In the low temperature regime, there was a trend for the diseased lobsters to have more bacteria than the healthy lobsters (Mann-Whitney: low/NME: Z = -1.890, *p* = 0.059, Figure 2C). There was no difference in bacterial counts between healthy and diseased lobsters in the mid and high temperature regime (Mann-Whitney: mid/SME: Z = -0.867, *p* = 0.386; high/SNE: Z = -1.757, *p* = 0.079; Figure 2C).

### Warmer and cooler tank temperatures drove bacterial community clustering

Bacterial community similarity was calculated using both unweighted community distance based on presence/absence of bacterial taxa (Figure 3A), and weighted community distance, which reflects the presence/absence and relative abundances of bacterial SVs (Figure 3B). All lobster shell bacterial communities were distinct from tank water control samples. When considering experimental factors individually, timepoint of sampling was the most important factor driving bacterial community similarity and sample clustering, followed by geographic location/seasonal temperature simulations for the winter and summer timepoints (not baseline), shell disease index, and shell molting stage (Table 1). All samples taken from baseline clustered together, indicating similar bacterial communities on shells, and then clusters separated by geographic locations and the temperature regimes respective to those treatments at subsequent timepoints (Figure 3A, B). Within clusters by timepoint and geographic location/seasonal temperature, shell disease index was a strong driver of bacterial community clustering (Table 1). For geographic location, shell disease index, and shell molting stage, weighted metrics of bacterial community similarity resulted in higher F-values (Table 1), indicating a stronger effect of bacterial SV abundance in driving clustering than simply which SVs were present. This likely means that bacterial SVs which were unique to particular treatments may have been rare or made up a small proportion of the total community.

**Figure 3.**
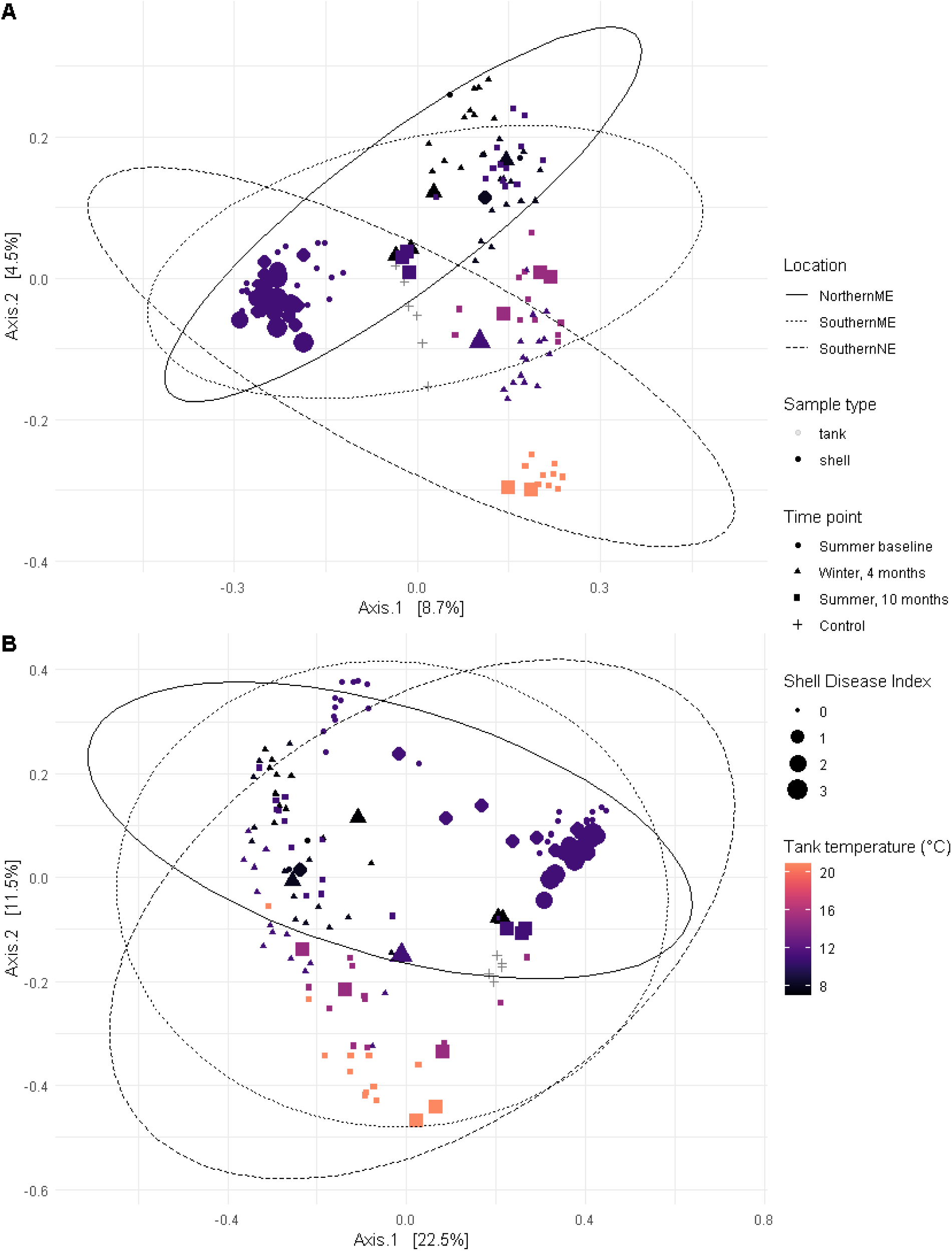
Principal coordinate analysis (PCoA) of lobster shell bacterial communities under three simulated seasonal ocean temperatures over time, using A) unweighted Jaccard or B) weighted Bray-Curtis distances to calculate community similarity. Tank temperature is indicated by color, epizootic shell disease stage by point size, timepoint by point shape, and simulated region by ellipses line type. Control samples, consisting of a 15 cm^2^ area swabbed from tank surfaces at the second timepoint, are shown in grey. Unweighted Jaccard uses presence/absence of bacterial SV and can be interpreted as a measure of whether taxa membership of the original community has changed. Weighted Bray-Curtis uses presence/absence of bacterial SVs and their relative abundance, and can be interpreted as a measure of whether different taxa are present under new conditions or if the original taxa remain but at different levels of abundance.

**Table 1.**
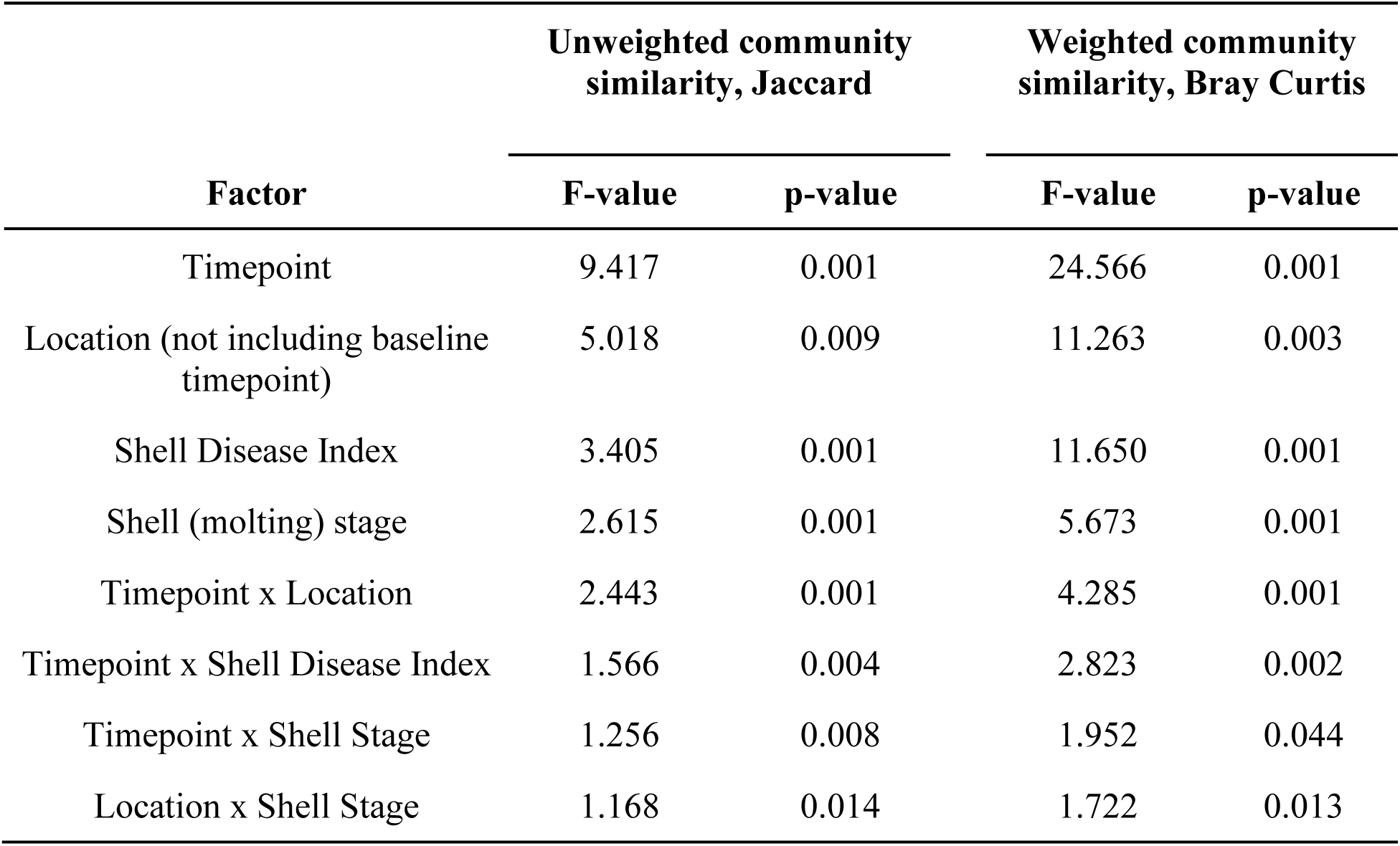
Permutational analysis of variance (permANOVA) comparisons for individual and interacting factors on the similarity of bacterial communities on lobster shells.

| Factor | Unweighted community similarity, Jaccard |  | Weighted community similarity, Bray Curtis |  |
| --- | --- | --- | --- | --- |
|  | F-value | p-value | F-value | p-value |
| Timepoint | 9.417 | 0.001 | 24.566 | 0.001 |
| Location (not including baseline timepoint) | 5.018 | 0.009 | 11.263 | 0.003 |
| Shell Disease Index | 3.405 | 0.001 | 11.650 | 0.001 |
| Shell (molting) stage | 2.615 | 0.001 | 5.673 | 0.001 |
| Timepoint x Location | 2.443 | 0.001 | 4.285 | 0.001 |
| Timepoint x Shell Disease Index | 1.566 | 0.004 | 2.823 | 0.002 |
| Timepoint x Shell Stage | 1.256 | 0.008 | 1.952 | 0.044 |
| Location x Shell Stage | 1.168 | 0.014 | 1.722 | 0.013 |

Contrarily, even though there were more cultured isolates in warmer tanks, the diversity of the isolated community was not changed, which may reflect the limited capacity of culture-dependent techniques. The Shannon-Weiner diversity index was calculated for each group of lobsters at each timepoint (Table 2) and indicates that the diversity indices of all groups are similar between healthy versus diseased and between sampling timepoints.

**Table 2.**
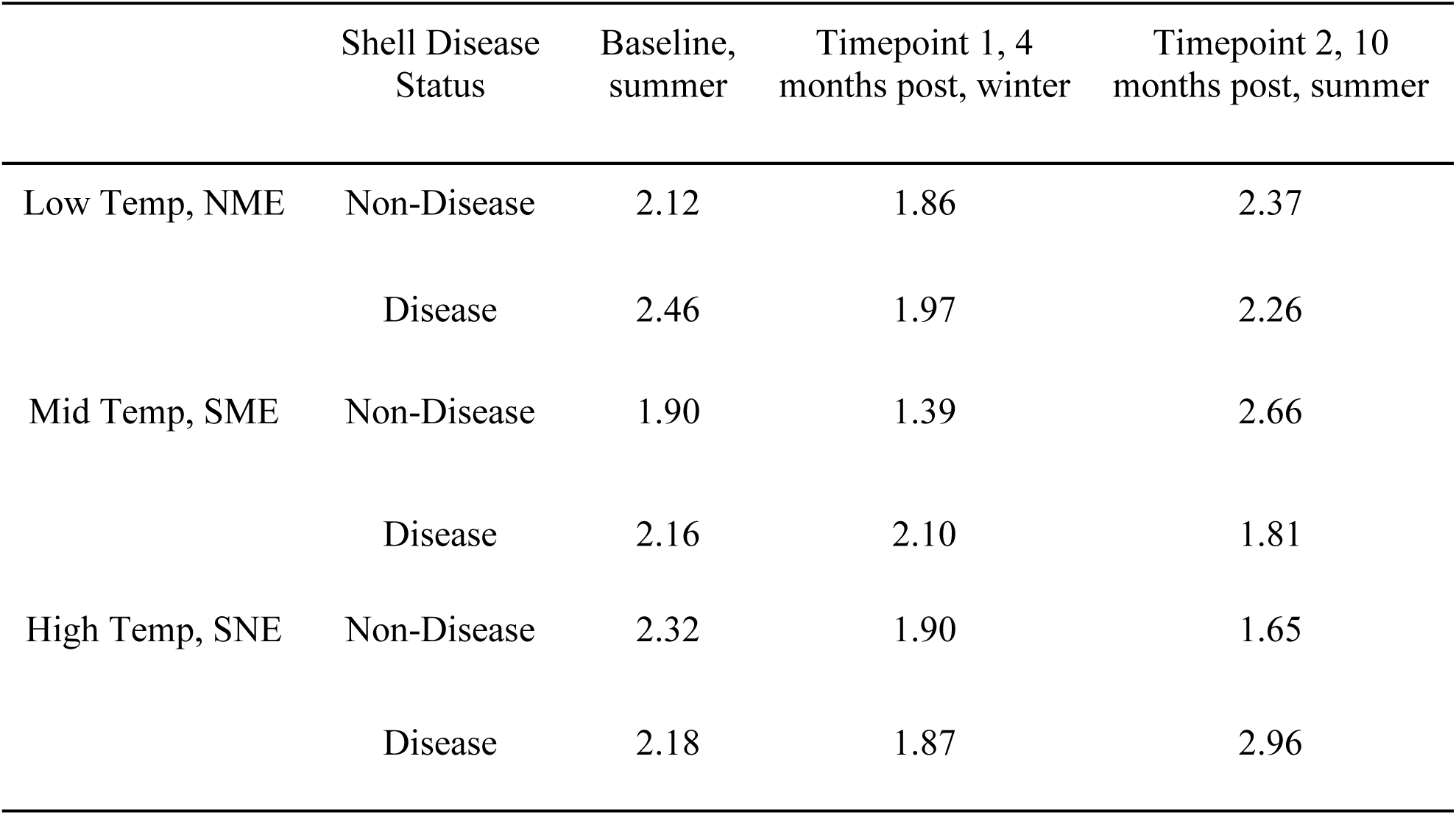
Shannon-Weiner diversity indices of cultured isolates calculated for each group of lobsters at different temperature regimes, timepoints, and health status. Baseline, with all three temperature treatment systems at an average temperature of 12.5°C. B; Time 1, average system temperatures: Low (NME) = 7.5°C, Mid (SME) = 8.5°C, and High (SNE) = 11.0°C. C; Time 2, average system temperatures: Low = 10°C, Mid = 15°C, and High = 21.0°C.

|  | Shell Disease Status | Baseline, summer | Timepoint 1, 4 months post, winter | Timepoint 2, 10 months post, summer |
| --- | --- | --- | --- | --- |
| Low Temp, NME | Non-Disease | 2.12 | 1.86 | 2.37 |
|  | Disease | 2.46 | 1.97 | 2.26 |
| Mid Temp, SME | Non-Disease | 1.90 | 1.39 | 2.66 |
|  | Disease | 2.16 | 2.10 | 1.81 |
| High Temp, SNE | Non-Disease | 2.32 | 1.90 | 1.65 |
|  | Disease | 2.18 | 1.87 | 2.96 |

Random forest feature prediction was used to identify bacterial taxa which were contributing to significant differences between microbial communities at different timepoints by temperature treatment (Figure 4). Certain taxa were absent from baseline and abundant on all shell samples at later timepoints regardless of treatment, including several SVs in the Flavobacteriaceae family. These were likely enriched in the tank aquaculture environment and not related to treatments. Others, such as SVs in the Rhodobacteraceae family, were likewise absent at baseline but appeared at later timepoints in Southern Maine and Southern New England treatments, implicating temperature as enriching for those.

**Figure 4.**
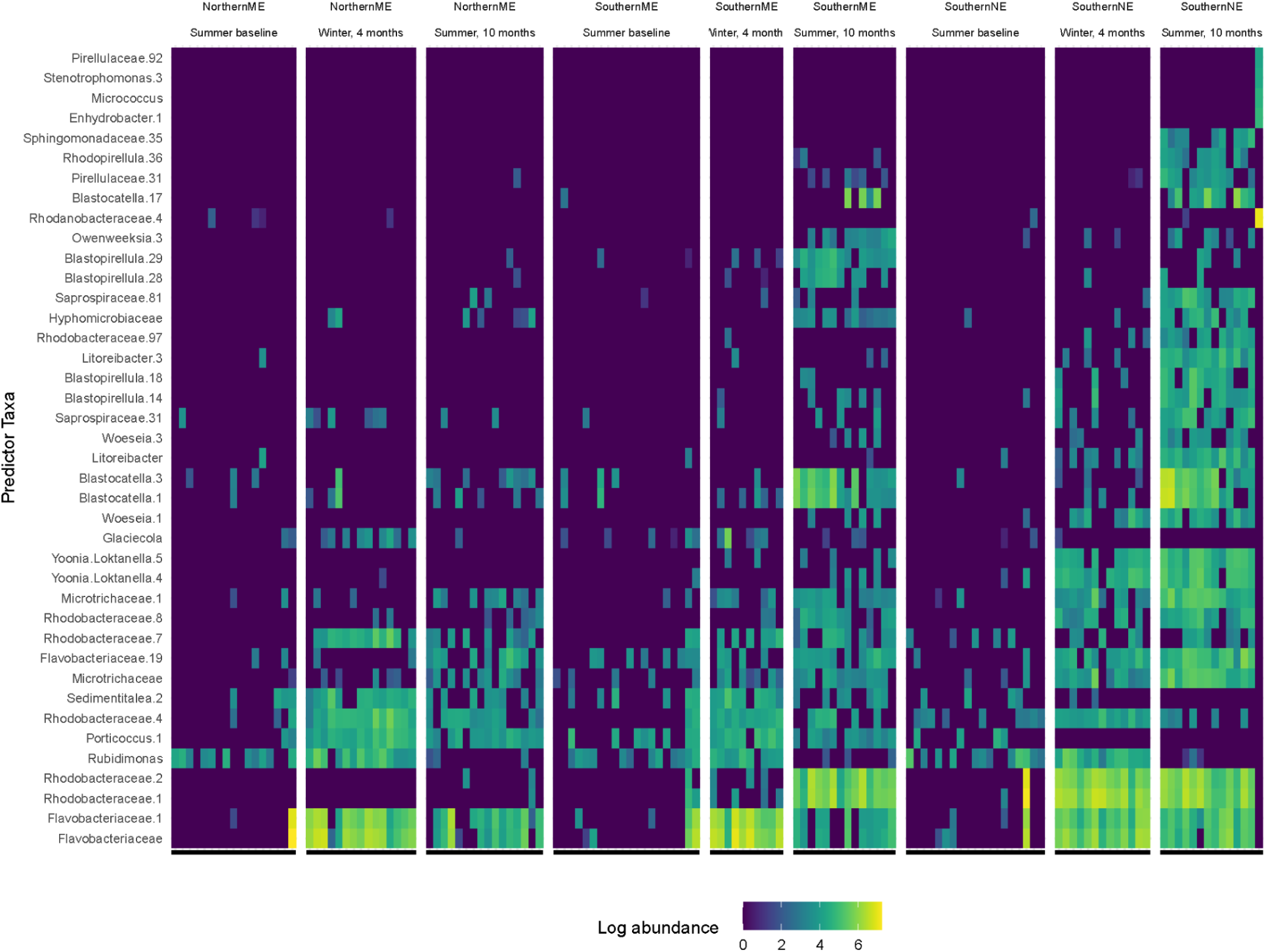
Relative abundance of bacterial taxa which were identified as important members of the lobster shell community associated with tank temperatures that simulate three geographic locations off the coast of New England. Taxa that were significantly important (*p* < 0.05) to the community structure with respect to tank temperature were identified using a permutational random forest algorithm. Log abundance is shown in the color scale, and columns are individual lobster shell samples. Columns are paneled by geographic location and timepoint, which reflects respective seasonal temperatures simulated in the tank. Baseline: temperature of 12.5°C on average for all three systems. Winter, 4 months: average system temperatures were Northern ME = 7.5°C, Southern ME = 8.5°C, and Southern NE = 11.0°C. Time 2: average system temperatures were Northern ME = 10°C, Southern ME = 15°C, and Southern NE = 21.0°C. Model accuracy was 79%, and the top 40 of the 207 significant bacterial SVs are shown.

### Shell bacterial diversity was not affected by weight or growth

There was a significant increase in weight for healthy and diseased lobsters across the baseline and last sampling periods (healthy lobsters: paired *t* test: *t*_24_ = -6.086, *p* < 0.001; shell diseased lobsters: paired *t* test: *t*_16_ = -4.328, *p* = 0.001) indicating lobster growth during the study duration. Non-shell diseased lobsters grew on average 13.9% of their original body weight (range = -9 to 30%) and shell diseased lobsters grew on average 10.2% of the original body weight (range = -11 to 22%). Shell length was associated with slightly lower bacterial richness (-2 SVs, lm t = - 2.566, *p* = 0.0113), although this effect is likely conflated with the trend of reduced bacterial diversity over time in the tanks during the experimental treatments. Bacterial richness was not affected (linear regression, *p* > 0.05) by lobster weight.

While in tanks for the year, 61% of the healthy lobsters and 59% of the diseased lobsters molted. Shell molt stage progression was associated with higher bacterial richness (Figure 5; Figure S1), but we posit that this is conflated with the high richness at baseline when lobsters were pulled from warm ocean waters in the summer and were nearing their first molt. The postmolt stages (A, B, C1/C2), when the carapace is soft and beginning to harden, exhibited the smallest increases in bacterial richness compared to stage A; stage B, +38 SVs (lm, t = 2.070, *p* = 0.040); stage C1/C2, not significant (*p* > 0.050). The intermolt stages (stages C3/C4), in which the shell is hardened and supports well-defined layers, exhibited much more bacterial richness compared to stage A; stage C3/C4, +61 SVs (lm, t = 3.374, *p* < 0.001). The greatest increase compared to Stage A was seen in the premolt stage (D) +114 SVs (lm, t = 6.613, *p* < 0.001), when a new carapace begins forming underneath and before molting occurs. Random forest models could not predict molt stage from bacterial community features with more than 57% accuracy (data not shown).

**Figure 5.**
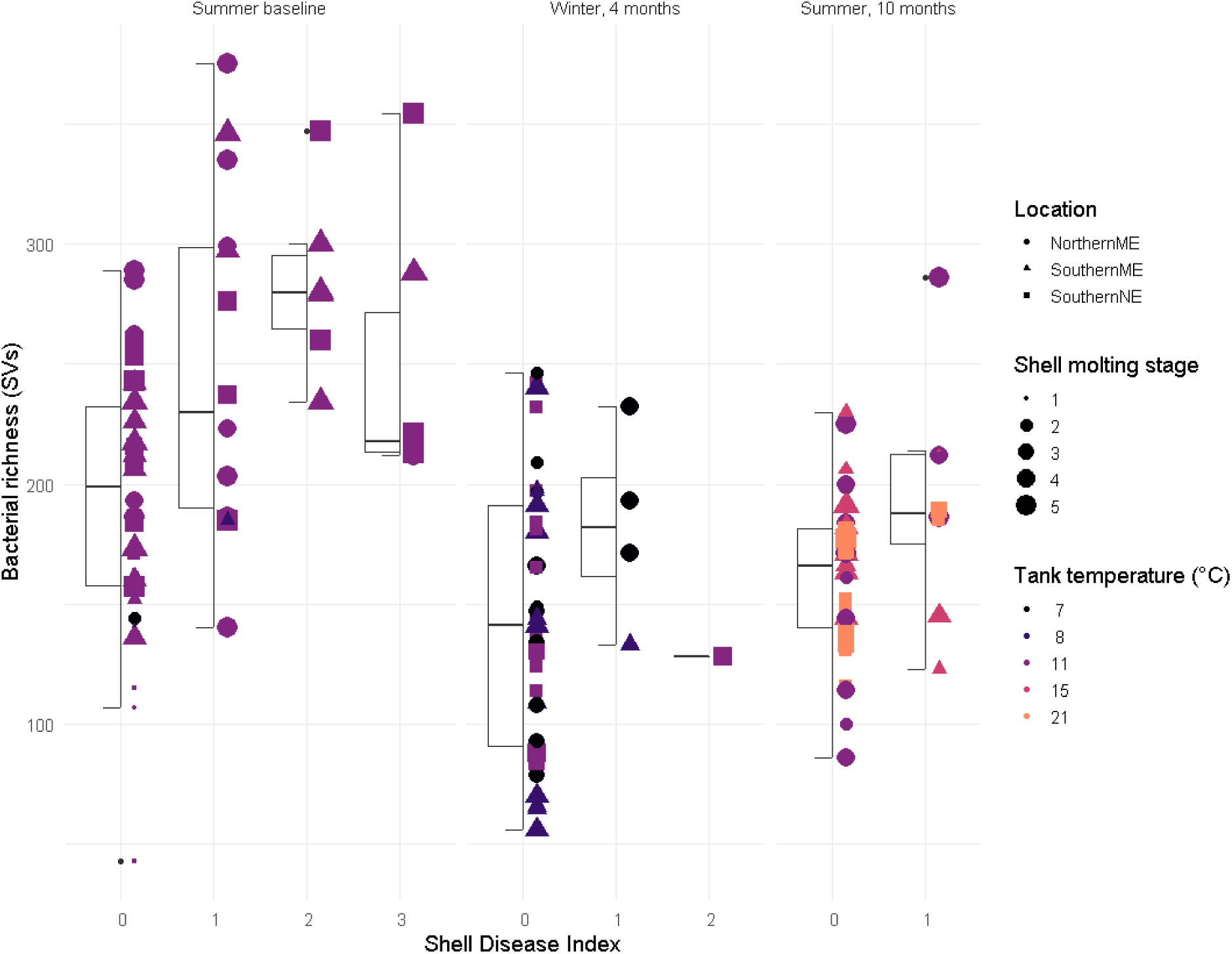
Bacterial richness (sequence variants; SVs) on lobster shells by Shell Disease Index under three simulated seasonal ocean temperatures over time.

### Shell bacteria and tank temperature were correlated with ESD progression, but not antimicrobial activity of plasma

Shell disease index scores indicated the percentage of the lobster shell covered by visible disease symptoms (Smolowitz et al., 2005). Scores for the entire cohort were higher at baseline than after 4 months in tanks undergoing temperature treatments, -0.60 (linear model, t = -4.337, p-value < 0.001), or after10 months, -0.55 (lm, t = -3.861, p-value < 0.001). This may reflect that lobsters which successfully molted were able to recover, and lobsters which did not successfully molt died, and only a few lobsters with shell disease had worsening symptoms. Shell disease index was not affected by tank water temperature (lm, p-value >0.5), confirming the complex etiology of the disease. Shell disease index was 1 rank higher in lobsters in the premolt stage (lm, t = 4.946, p-value < 0.001).

We examined the antimicrobial activity of the plasma based on two contrasting protocols (Homerding et al., 2012; Noga et al., 1994), to determine if ESD lobsters displayed reduced resistance to bacterial growth *in vitro*. At baseline, despite the higher severity indices of shell disease, there was no *E. coli* D31 growth inhibition from any samples and therefore no differences between the antimicrobial activity of plasma collected from healthy and diseased lobsters at 12.5**°**C (ANOVA: F_2,47_ = 1.428, *p* = 0.25). At 4 months in tanks, at the winter sampling, again there was no demonstrated *E. coli* D31 growth inhibition and the antimicrobial activity of plasma from healthy lobsters was not affected by temperature at 7.5**°**C, 8.5**°**C, or 11.0**°**C (ANOVA: F_3,23_ = 1.56, *p* = 0.23). There was also no *E. coli* D31 growth inhibition or altered plasma antimicrobial activity observed in diseased lobsters held at 7.5**°**C, 8.5**°**C, or 11.0**°**C (ANOVA: F_3,18_ = 1.083, *p* = 0.468). When comparing the antimicrobial activity of plasma collected from pooled healthy and diseased lobsters across temperatures 7.5**°**C, 8.5**°**C, and 11.0**°**C, there were no differences (ANOVA: F_2,44_ = 1.261, *p* = 0.29). After 10 months in tanks, in the summer sampling, the antimicrobial activity of plasma from healthy lobsters was not affected by temperature at 10.0**°**C, 15**°**C, or 21**°**C (ANOVA: F_3,23_ = 0.0612, *p* = 0.98). Results of antimicrobial activity of plasma from diseased lobsters was also unaffected by temperature at 10.0**°**C, 15**°**C, and 21**°**C (ANOVA: F_3,16_ = 0.897, *p* = 0.46). When comparing the pooled antimicrobial activity of plasma collected from healthy and diseased lobsters across temperatures 10.0**°**C, 15**°**C, and 21**°**C there were no differences (ANOVA: F_2,42_ = 0.0582, *p* = 0.94).

Shell disease index was associated with significantly increased bacterial community richness on shells compared to apparently healthy shells, and was somewhat progressive by severity (Figures 1A, 5): shells with an index score of 1 had 58 SVs more than apparently healthy (lm, t = 4.697, *p* < 0.001); index 2 had 98 more SVs (lm, t = 4.399, *p* < 0.001); and index 3 had 84 more SVs than apparently healthy lobsters (lm, t = 43.528, *p* < 0.001). Shell disease also appears to increase the evenness of bacterial SVs (0 - 1 scale; Figure 1B); at index level 1 (lm, increase = 0.08, t = 4.58, *p* < 0.001), level 2 (lm, increase = 0.09, t = 2.91, *p* = 0.004), and level 3 (lm, increase = 0.09, t = 2.72, *p* = 0.007).

Epizootic shell disease is associated with altered bacterial community total richness, as well as the membership of that community. However, when assessing the core community shared by 80% of lobsters with either apparently healthy (Figure 6A) or diseased shells (Figure 6B), it is apparent that many of the same core bacteria and putative agents are present on shells regardless of health status. When comparing the bacterial communities in lobsters at baseline, the time when the largest number of lobsters displayed the greatest spread of disease symptom progression, there were several bacterial species which were significantly correlated with disease progression (Figure 7). They included multiple SVs in the genera *Aquimarina*, *Kiloniella, Arenicella,* and in the families Flavobacteriaceae and Rhodobacteraceae, which were more prevalent and more abundant in lobsters with more advanced disease states (Figure 7). The model explained 44% of the variation in lobster shell bacteria at baseline.

**Figure 6.**
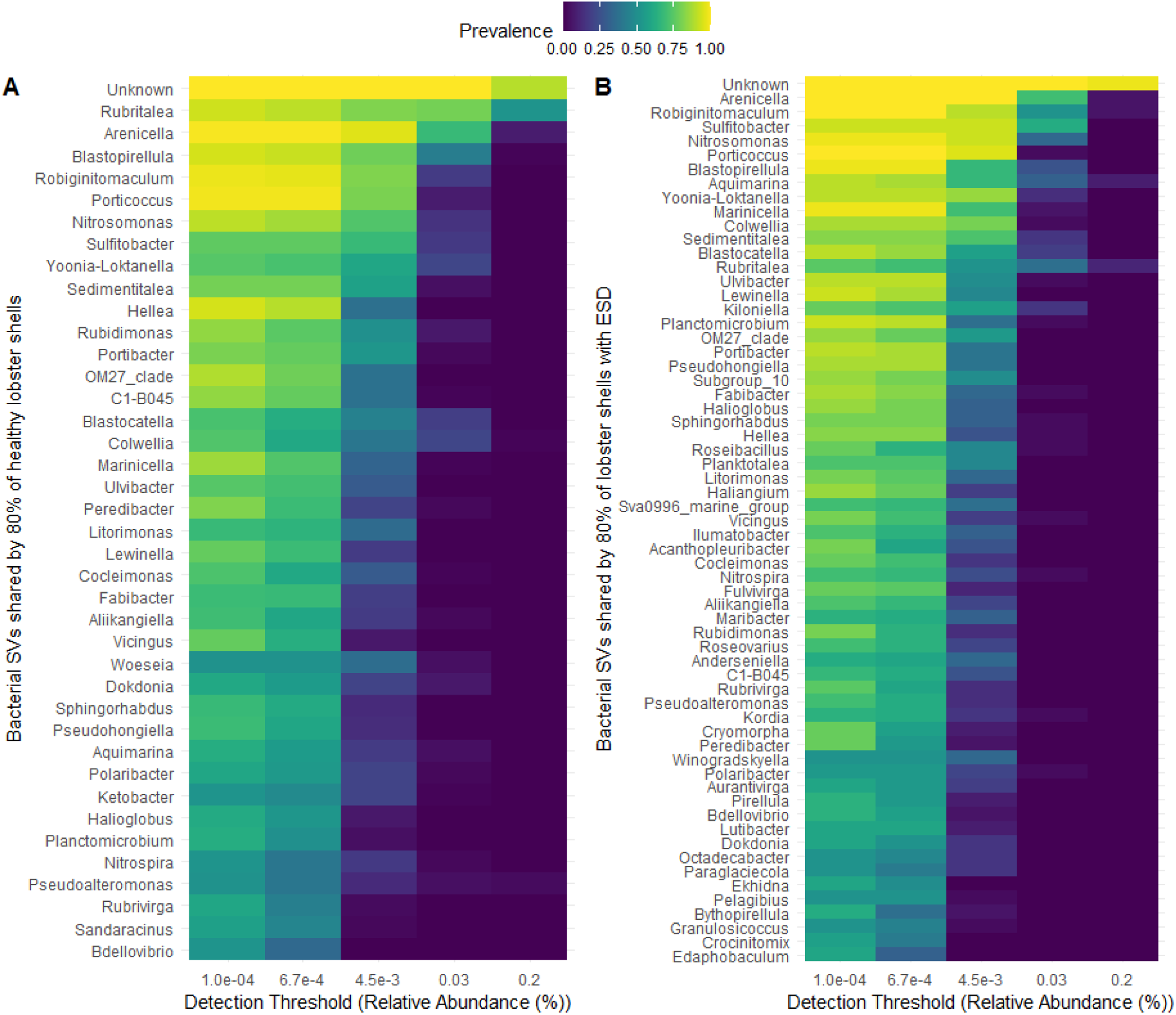
Core shell carapace bacterial sequence variants (SVs) shared by 80% of lobsters which are A) healthy (no visible disease signs) or B) epizootic shell diseased. Prevalence of bacterial SVs across 0 - 100% of lobster shells is shown in color, and SVs are ordered by the most to least prevalent, based on percent relative abundance thresholds.

**Figure 7.**
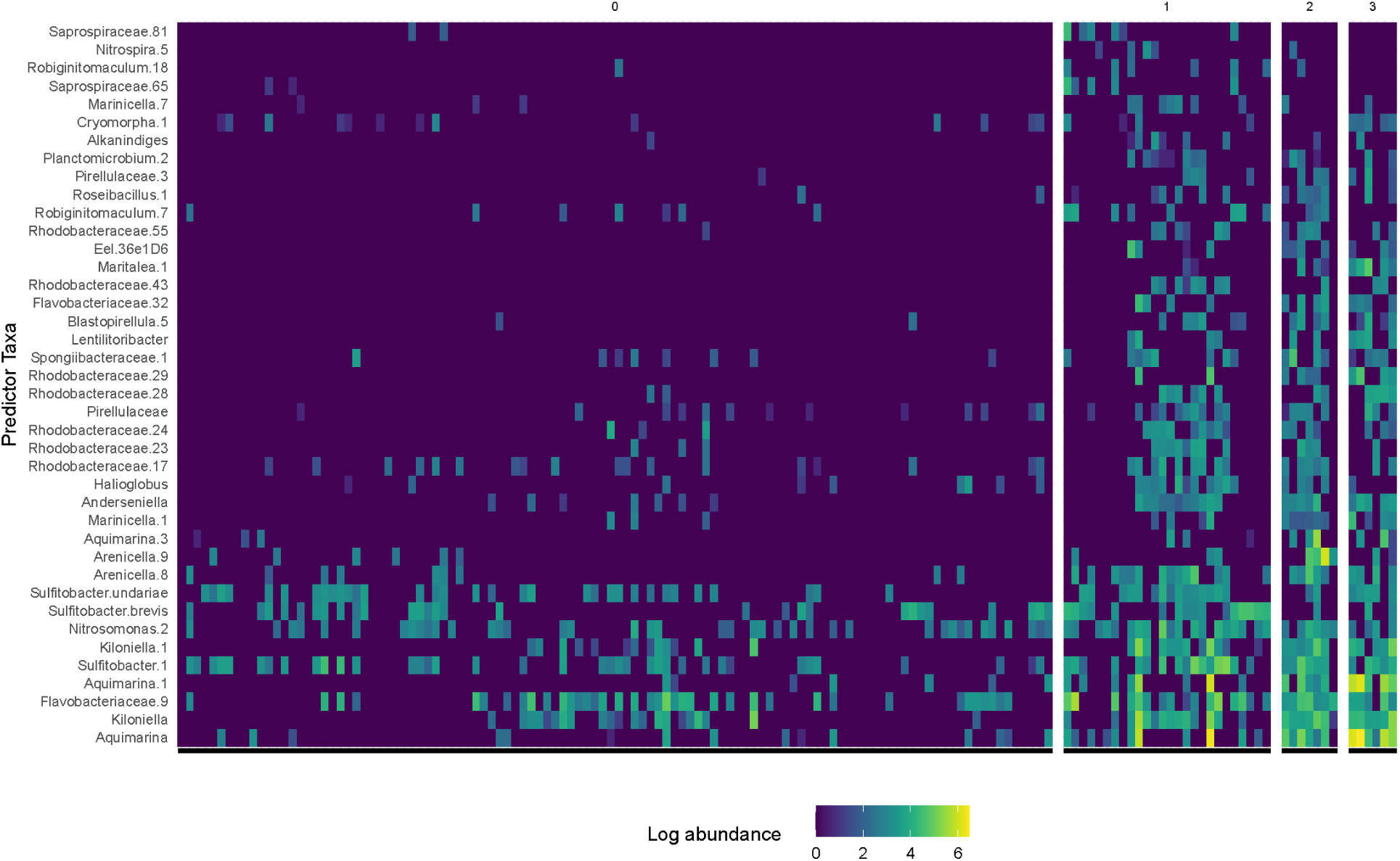
Relative abundance of bacterial taxa which were identified as important members of the lobster shell community associated with epizootic shell disease in fresh-caught lobsters off the coast of New England. Significantly important (*p* < 0.05) taxa were identified using a permutational random forest algorithm, performed on a data subset consisting only of the fresh-caught, baseline lobster shell samples prior to temperature treatments. Log relative abundance is shown in the color scale, and columns are individual lobster shell samples. Columns are paneled by the Shell Disease Index ranking: 0, no observable disease; 1, disease on 1 - 10% of the shell; 2, disease on 11 - 50% of the shell; and 3, disease on >50% of the shell. Model explained 44% of the variation in lobster shell bacteria at baseline.

### Tank temperature was not associated with mortality, but some bacteria were

Of the 57 lobsters at the beginning of the experiment, 15 died (Figure 8): 1 from NME healthy group, 1 from NME diseased group, 1 from SME healthy group, 5 from SME diseased group, 2 from SNE healthy group, 4 from SNE diseased group. A Pearson chi-square contingency analysis was used to determine the relationship between shell disease and the likelihood of mortality (JMP v13). Lobsters with shell disease were more likely to die (*P* = 0.043) than apparently non-shell diseased lobsters.

Bacterial richness on shells (number of SVs) does not appear associated with mortality events (Figure 8), and while there were sequences from putative pathogens found on ESD shells (Figure S2), there was not a correlation between mortality and extinguishing potentially pathogenic genera on shells (Figure S3). Lobsters with ESD or which died did not appear to have bacteria communities on their shells which were more similar to the bacterial communities found in tank water control samples (data not shown). This might have implied a loss of host-associated bacteria; however, this could not be thoroughly assessed as there were only single-timepoint samples for tank water community controls.

**Figure 8.**
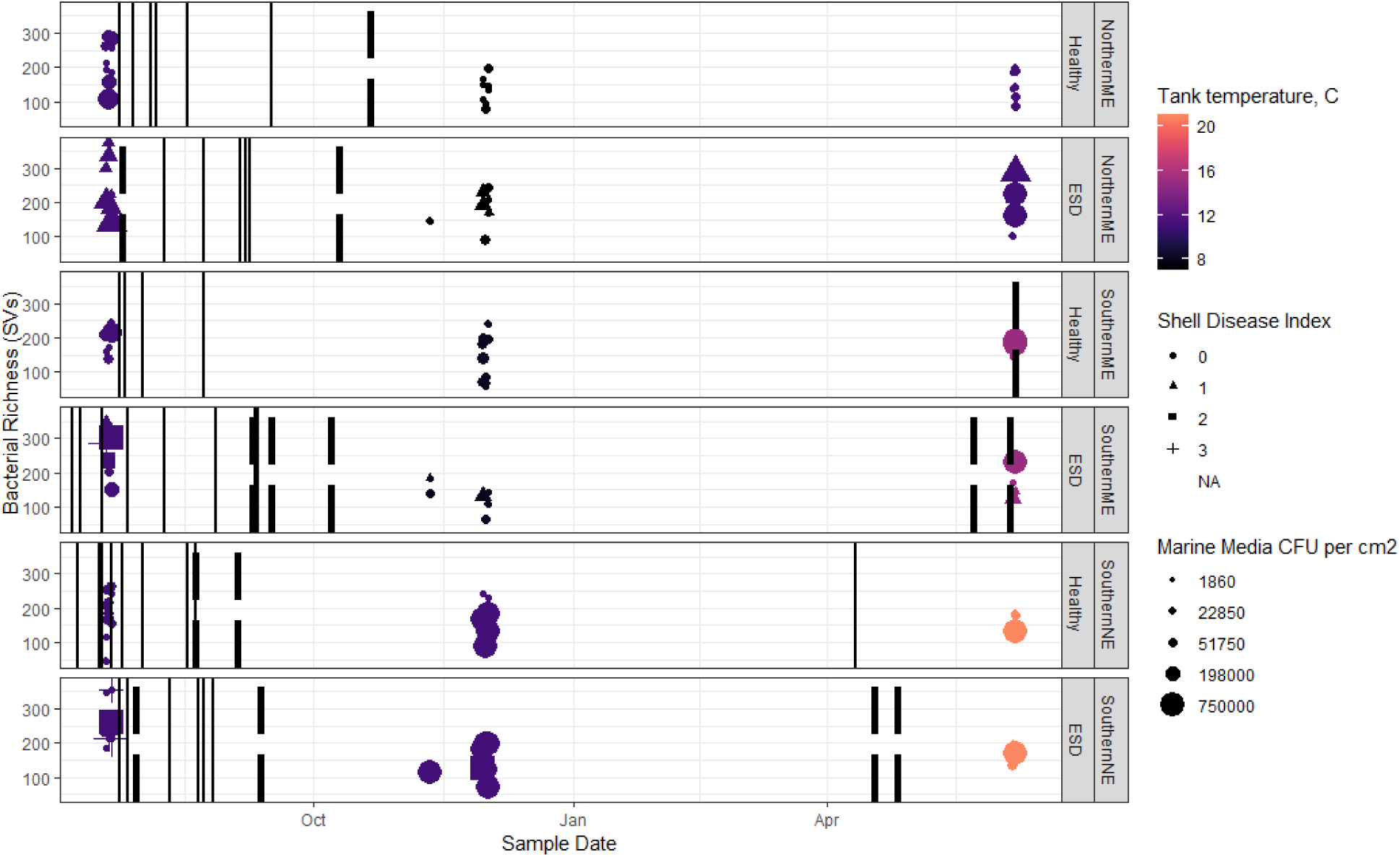
Bacterial richness (sequence variants) in relation to the number of bacterial colony forming units (CFUs) isolated from lobster shells on marine media over time. Shape denotes Shell Disease Index, size of points denotes colony forming units per centimeter squared on general Marine Media, color denotes the tank temperature at the time of sampling. Sampling date is along the x-axis; solid lines denote when a lobster molted, and dashed lines denote when a lobster died. Vertical panels represent each experimental tank and the geographic region its seasonal temperatures were modeled after. Shell-diseased lobsters at the start of the experiment were groups in tanks 2, 3, and 6.

Yet, these putative agents may play a role in the mortality rate of lobsters, rather than discerning between a healthy and diseased state. Important bacterial members for defining the community state on shells were identified in a subset of samples which included only those shells with visible signs of disease (Disease Index 1-3). In this subset of 39 samples, lobsters which died during the experiment were compared to those which didn’t, revealing different abundances of bacteria of interest. These are visualized by mortality and by shell disease index in Figure 9, a model with 86% accuracy which identified 112 bacterial taxa which were significantly different by groups.

**Figure 9.**
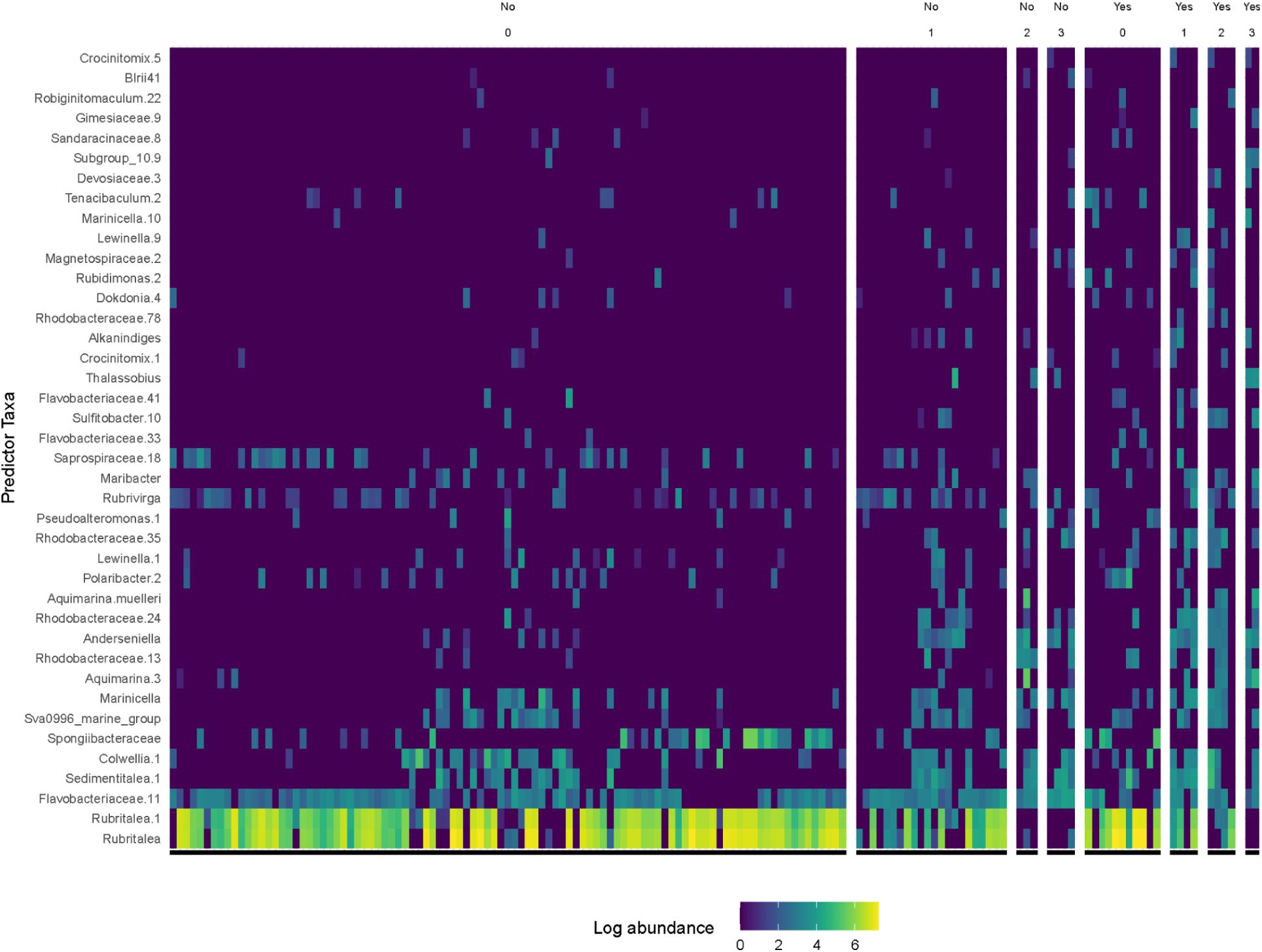
Abundance of bacterial taxa which were identified as important members of the lobster shell community associated with lobsters which died or not. Log abundance is shown in the color scale, and columns are individual lobster shell samples. Columns are paneled by mortality (yes/no) and the Shell Disease Index ranking: 0, no observable disease; 1, disease on 1 - 10% of the shell; 2, disease on 11 - 50% of the shell; and 3, disease on >50% of the shell. Only lobsters with visible shell disease were used in the model. The top 40 of 112 total statistically significant (*p* < 0.05) SVs are shown, as designated by permutational random forest analysis. Model accuracy was 86%.

Using the two media, 725 isolates were identified from lobster shells and 10 from tanks by phenotypic identification and/or 16S rRNA gene sequencing. The purpose of this work was to determine the microbial ‘stability’ as in changes to community profile over time and temperature. For review, Biolog Microbial Identification System (BMIS) provides identification by means of phenotypic/biochemical characteristics. BMIS results range in probability of identification and can provide high probability to the species level while some results provide a genus with moderate to low probability of species and others to genus only. There were 4 phyla, 20 families and 36 genera of bacteria identified (Table S1), inclusive of the genera most frequently associated with ESD such as *Aquimarina, Flavobacterium, Polaribacter, Sulfitobacter, Pseudoalteromonas*. We did not isolate *Thalassobius* species, a putative agent for ESD.

The five predominant isolates from two media types were grown from lobster shells at the three sampling timepoints, yielding 725 bacterial isolates (Figure 10): 100 from the 15 lobsters that died during the study, and 625 from the 42 lobsters that did not die. Of the 100 isolates from lobsters which died, 48% were oxidase positive (n = 48), 17% were catalase positive (n = 17), 63% were gram negative rods (n = 63), 3% were gram negative cocci (n = 3), 15% were gram positive rods (n = 15), and 4% were gram positive cocci (n = 4). Of the 625 isolates from lobsters which survived, 58% were oxidase positive (n = 363), 8.8% were catalase positive (n = 55), 70% were gram negative rods (n = 441), 3% were gram negative cocci (n = 21), 12.8% were gram positive rods (n = 80), and 2.7% were gram positive cocci (n = 17).

**Figure 10.**
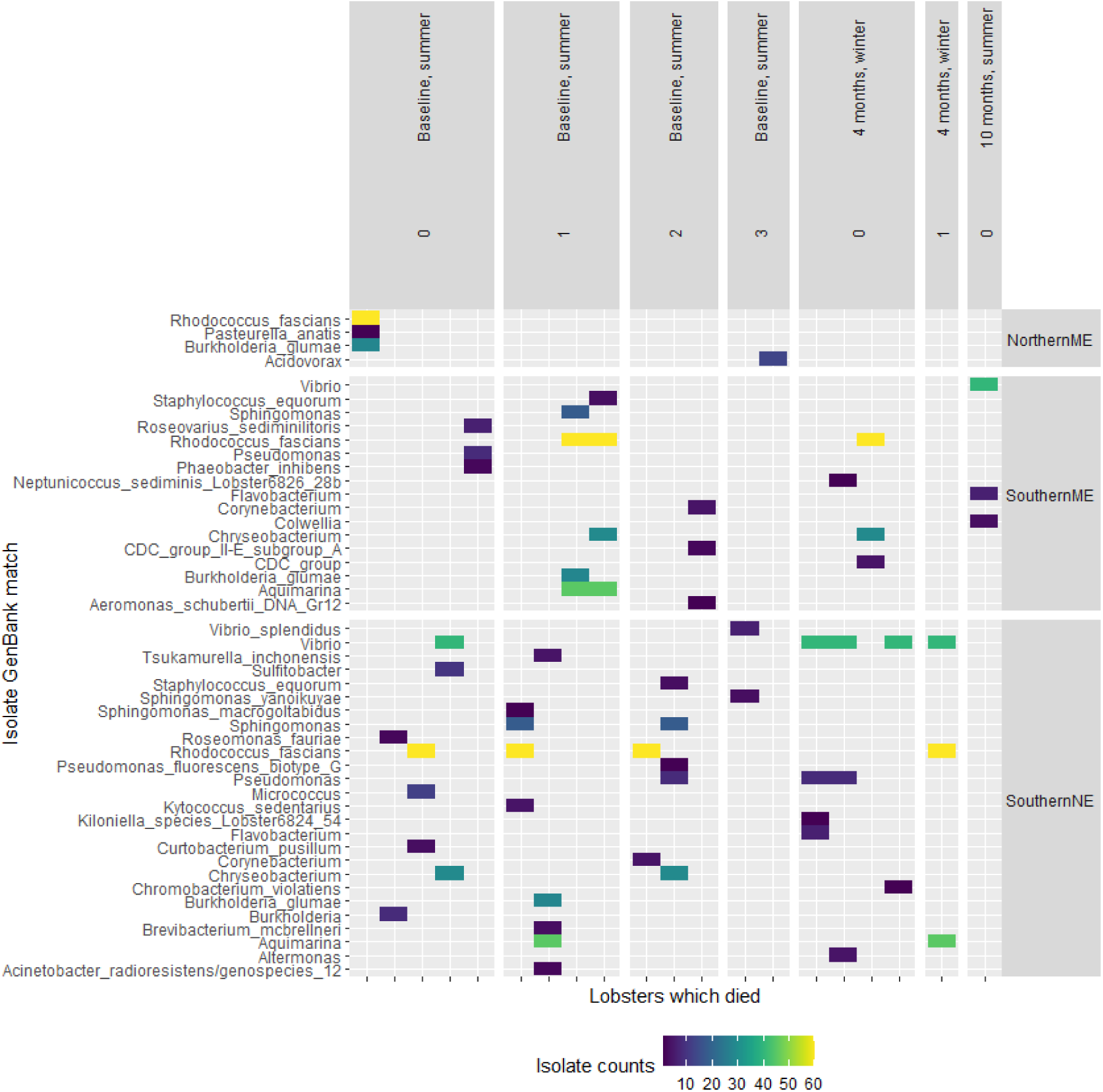
Comparison of bacteria isolated from shells of lobsters which survived or died during the experiment. Bacterial isolates were identified using 16S rRNA sequencing and NCBI GenBank sequence reference database. Columns are paneled by mortality (yes/no) and the Shell Disease Index ranking: 0, no observable disease; 1, disease on 1 - 10% of the shell; 2, disease on 11 - 50% of the shell; and 3, disease on >50% of the shell. GenBank match of “n/a” indicates isolate could not be identified at any the genus level of taxonomy.

Visual comparisons on isolates which were found on healthy and diseased lobsters showed no obvious differences, in that most isolates were found on both populations, however, there were some differences in lobsters which died during the experiment (Figure 10), as compared to those which survived, by geographic location (Figures S4 - S6). There was some agreement between the identity of cultured isolates and what was found in the bacterial community sequencing data, including *Aquimarina, Sulfitobacter, Colwellia*, and others (Figure S7).

## Discussion

Our goals were to 1) determine the effect of temperature on ESD progression, mortality, and immune health; 2) assess the stability of the lobster shells’ microbial community over time and temperature, and 3) identify correlations between temperature, lobster health, cultured bacteria from shells, and whole bacterial communities on shells. This study obtained 57 female lobsters from coastal waters off southern Maine, and maintained them in tanks for nearly a year with three geographical temperature profiles simulating seasonal water temperatures of the location lobsters were sourced from, a warmer location, or a colder location.

In all groups, bacterial community diversity was significantly reduced after baseline. It is not surprising that going from biodiverse ocean waters to environmentally static aquaria would reduce bacterial communities on lobster shells, regardless of water temperature or shell disease status. For example, tropical marine water lost bacterial diversity over time when stored in a ballast tank (Ng et al., 2015), and aquacultured tank systems typically have lower microbial diversity than seawater because microorganisms are filtered or killed to prevent disease and parasite load in animals. Marine animals acquire many of their associating microorganisms from their environments (Boscaro et al., 2022; Ishaq et al., 2022; Sousa et al., 2021), and even with a diverse shell microbiota at baseline it was expected that all lobsters would lose some bacterial diversity over time without a complex environment from which to replenish microorganisms.

### Temperature in tanks was not solely responsible for ESD symptoms or mortality

Based on our tank-based study, temperature alone does not affect the progression of ESD and indicates the complexity of disease occurrence and progression in diverse natural marine environments. We observed minimal increase in ESD progression over the 11 months, similar to other studies observing lobsters in aquaria (Stevens, 2009). During the course of our study, 61% of the healthy lobsters and 59% of the diseased lobsters molted, and lobsters may recover from ESD if they survive molting (Stevens, 2009). Temperature, food availability, and reproductive status can impact the timing of molt, which can alter lobsters’ susceptibility to bacteria or the environment, as discussed previously (Ishaq et al., 2022). In this study, the diseased lobsters that molted and re-developed signs of shell disease were in the mid and high range temperature groups. There was a significant weight gain for all lobster groups with no significant difference between non-shell diseased and shell diseased lobsters over the course of the study, implying that lobsters did not get sicker while in tanks. This was different from Stevens (2009) who suggested slower growth rates due to ESD. Low mortality was observed in healthy lobsters in all temperature regimes, and while there were more mortalities in the shell diseased lobsters the difference was not significant. In the SME temperature regime, there was a trend for the ESD lobsters to have higher mortality than non-shell diseased, and if our study had higher numbers of animals per treatment group, this trend might have been better defined.

We examined the antimicrobial activity of the plasma based on previous literature (Homerding et al., 2012; Noga et al., 1994). Noga et al. used a turbidimetric assay to determine that antibacterial activity of plasma was higher in healthy compared to shell-diseased blue crabs, *Callinectes sapidus,* and that there were differences based on populations at various geographical sites. Homerding et al. used this assay but did not identify a difference in the antibacterial activity of plasma from *H. americanus* lobsters with and without ESD, although did find a difference based on geographic locations of the lobsters. Our study results did not demonstrate significant bacterial growth inhibition; thus, no measurable antibacterial activity was seen in any of the lobster hemolymph over any of the timepoints when evaluating a particular group. The optical densities of individuals at testing timepoints had high inter-individual variability but the overall population results indicated no antibacterial response. The lack of antimicrobial activity in our study may indicate that in a controlled laboratory environment the stress factor of temperature alone did not result in a measurable plasma antibacterial immune response. In regard to the study by Homerding et al. 2012, there may have been other environmental stressors contributing to plasma response. Of note, Noga et al., used serum (coagulated hemolymph with clot removed) and our procedure followed Homerding et al. and used plasma (centrifuged hemolymph prior to clotting to remove hemocytes). The innate immune response of decapod crustaceans has integrated cellular and humoral components, both cellular and plasma responses work in coordination (Hauton, 2012). The procedure of immediately separating the hemocytes from the plasma may have reduced or eliminated antimicrobial proteins secreted from hemocytes during the clotting process. Investigating the serum antimicrobial activity may have provided measurable responses.

### Temperature altered bacterial communities on shells

Microbial isolate enumeration and identification for each lobster shell was performed 3 times approximately 3 months apart reflecting seasonal temperatures of summer, winter and peak summer for each regional area described. The same anatomical location was swabbed on each lobster. For clarity, swabbing was not targeted specifically at diseased areas on lobsters with signs consistent with ESD but rather the same anatomical site on the lobster, no-shell disease or diseased.

Baseline bacterial enumeration indicated that the bacterial loads on shell diseased lobsters were significantly higher than on healthy lobsters. As seasonal temperature cycles progressed, the bacterial counts at the 4-months post winter sampling, and the 10-months post summer sampling, were significantly higher in the high (SNE) temperature range, whether healthy or ESD, over counts from lobsters held at the mid (SME) and low (SNE) temperature ranges. However, in the high (SNE) and (mid) SME range there was no difference in bacterial load on healthy versus ESD animals. This may reflect that warmer water is a general selective pressure acting against most bacteria present, as warming decreases bacterial diversity in the marine environment (Traving et al., 2021) and organisms (Li et al., 2018; Minich et al., 2018). In the low (NME) temperature system significantly higher bacterial loads were still observed on the diseased versus healthy lobsters, which may reflect that colder water is a specific selective pressure affecting some of the bacteria present. It may also reflect that bacteria conditioned to warmer temperatures are faster growing and simply performed better in the lab.

### Certain bacteria were associated with healthy or ESD status

We compared the frequency of identified microorganisms associated with ESD and compared the frequency of isolation over time and temperature on apparently healthy and diseased lobsters. Using a combination of BMIS phenotypic identification and 16S rRNA gene Sanger sequencing, we isolated and identified organisms from the *Aquimarina, Chryseobacterium, Empedobacter, Flavobacterium, Polaribacter, Pseudoalteromonas, Roseomonas, Roseovarius, Sulfitobacter* and *Vibrio* genera which have all been reported as associated with ESD (Bell et al., 2012; Chistoserdov et al., 2012, 2005; Meres et al., 2012). *Aquimarina* and *Thalassobius* are frequently examined as pathogens, although more recently have been considered as opportunistic commensals already living on lobster shells (Meres, 2016). We routinely isolated *Aquimarina* and found them in the bacterial community sequencing data, but we did not identify *Thalassobius* in culture or in sequencing. We did, however, identify closely related genera from the *Rhodobacteraceae* family. There were no distinct patterns of frequency of isolation, the isolates most associated with ESD were isolated from both healthy and diseased lobsters present in all temperature groups at all sampling timepoints, and the Shannon-Weiner diversity indices for cultured isolates were similar for all groups. Our study results are similar to a study that examined the bacterial species in the cuticles of European lobsters (*Homarus gammarus*) and American lobsters sharing the same aquarium systems (Whitten et al., 2014).

There were several bacterial taxa from the sequencing data which were associated with ESD progression or absence in this study, which have been identified in previous studies, e.g., (Feinman et al., 2017; Meres et al., 2012; Whitten et al., 2014) which noted that many of these taxa were present in healthy and visibly-diseased lobsters in different amounts. In our samples, *Aquimarina*, which has long been implicated as a commensal-turned-pathogen in ESD (Quinn et al., 2017, 2012), increased in abundance in lobsters with lesions >11% of the shell (Index levels 2 and 3). We identified *Rubritalea* in the bacterial community data as abundant in healthy lobsters and reduced in those with ESD. We isolated *Gordonia* only from lobsters without shell disease and which did not die, which agrees with other findings in which *Gordonia* was only present on red rock lobsters (*Jasus edwardsii*) without tail fin necrosis (Pande et al., 2021). *Rubritalea* (Song et al., 2018; Yoon et al., 2011) and *Gordonia (Pande et al., 2021; Silva et al., 2022)* species have been identified in marine systems and associated commensals on the shells of marine animals where they produce carotenoids, which might act as antimicrobials. Carotenoids in lobster shells are typically sourced from plankton and other dietary components, but it is possible that microbially-produced carotenoids on the surface of shells are serving to drive shell communities. ESD lobsters are known to have less carotenoid content in their shells, including areas with or without lesions (Prince and Bayer, 2005).

### ESD could reflect the loss of a mutualist microorganism

Our findings agree with the previously published hypothesis (Meres, 2016) that ESD is not caused by the addition of an infectious agent, so much as a state of dysbiosis. We hypothesize that this may be due to the loss of a commensal which triggers unrestrained growth in the rest of the shell community. For example, evenness increased between bacterial SV abundance on diseased shells, implying that a keystone microorganism might be lost from lobster shells in the process of transitioning from a healthy to a diseased state, and without this keystone species the other bacteria present are able to thrive. This is similar to microbial ecological patterns in the other host ecosystems with keystone species, such as the human vagina when the loss of a *Lactobacillus* sp. allows for increased bacterial diversity which alters the function of the community and can lead to disease states, e.g., (Jašarević et al., 2015).

Specific symbiotic bacteria have been identified as critical to marine animals, for example to protect shrimp larvae from fungal infections (Gil-Turnes et al., 1989), provide nutrition and secondary molecules for defense in isopods (Lindquist et al., 2005), or to form the intricate connections between sponges and their microbiota (Pita et al., 2018). In deep sea squat lobsters, *Munidopsis alvisca*, bacteria and archaea on the shells of lobsters living near hydrothermal vents helped process potentially detrimental chemicals and may support the health of shells (Leinberger et al., 2022). While many of the bacteria in that study were novel and could not be identified, there were some found by Leinberger et al. that were also isolated in this study. These were isolates from the genera *Formosa* and *Pasteurella* from the shells of apparently healthy lobsters which survived the experiment.

Culture dependent and independent analyses are each subject to benefits and technological limitations, and even though the datasets could not be combined, their pairing allowed us deeper insight than either independently. For example, *Gordonia* were culturable but not dominant in the sequencing data and otherwise might have been overlooked. Conversely, *Rubritalea* was sequenced but not culturable under these conditions. The insight gained by the community analysis can be used to broaden culturing efforts to select for potential commensals or pathogens, which could then be used for whole-genome or transcriptomic analysis to assess activities, as well as in infection or rescue/probiotic trials with lobsters.

### Limitations of the study

The tank design and aquarium system limit the microbial diversity in tank water, which cannot accurately mimic the diversity and variability of ocean microbiomes, thus all microbial diversity on lobsters is reduced over time. Lobsters notoriously recover from ESD in tank systems, likely due to the lack of multiple and realistic stressors, making it difficult to ascertain the environmental conditions or microbial affiliations which exacerbate ESD symptoms. Further, because climate change has already been affecting microorganisms, hosts, and ecosystems, it is difficult to determine if the bacterial community found on healthy lobster shells represents their typical associated community. Culturing protocols can never be broad enough to isolate all bacterial diversity from samples, but by selecting the top five most predominant organisms observed by culture, we were able to determine frequency of isolation but not the quantity of each species isolated. Finally, we used amplicon sequencing analysis of total bacterial DNA fractions, which included DNA from living active/inactive and from dead/relic cells, thus we are unable to comment on the functionality of the bacterial community found.

## Author Contributions

Conceptualization, H.H., D.B.; Methodology, D.B.; Software, S.L.I., G.L.; Formal Analysis, S.L.I., G.L., M.S.T., S.M.T., D.B.; Investigation, H.H., D.B.; Resources, D.B.; Data Curation, S.L.I., S.M.T., D.B.; Writing - Original Draft, S.L.I., G.L., S.M.T., D.B.; Writing - Review and Editing; S.L.I., G.L., S.M.T., M.S.T., J.D.M., H.H., D.B.; Visualization, S.L.I., G.L., D.B.; Supervision, S.L.I., H.H., D.B.; Project Administration, D.B.; Funding Acquisition, D.B..

## Supporting information

R script

## Acknowledgments

We are grateful to the previous efforts of those who enabled us to conduct lobster research which led to the development of these concepts. This includes the State of Maine Department of Marine Resources, Maine Lobster Research Education and Development (RED) board, University of Maine Aquatic Animal Health Laboratory Staff, Emily Thomas, and Dawna Beane, as well as University of Maine undergraduate students Laurel Anderson, Emily Tarr, and Helen Reese.

## List of abbreviations

ANOVA: analysis of variance
Df: degrees of freedom
ESD: epizootic shell disease
I.e.: Latin for id est, “that is”
Lm: linear regression model
NME: northern Maine
PCOA: principal coordinate analysis
rRNA: ribosomal ribonucleic acid
SME: southern Maine
SNE: southern New England
SV: sequence variant

## STAR Methods

### RESOURCE AVAILABILITY

#### Lead contact

Further information and requests for resources and reagents should be directed to and will be fulfilled by the lead contact, Suzanne Ishaq.

#### Materials availability

This study did not generate new unique reagents.

#### Data and code availability

Raw sequencing reads from the V3-V4 region of the 16S rRNA gene from bacterial community sequencing data (Illumina MiSeq) and sample data, including about the sequencing run, lobsters, and environmental conditions, are publicly available through NCBI Sequence Read Archive (SRA). Trimmed and assembled reads from the 16S rRNA gene from cultured isolates sequence data (Sanger sequencing) publicly available through NCBI GenBank. All NCBI data are associated together under BioProject Accession PRJNA887451. Code for bacterial community sequence data is provided as Supplemental Materials.

### EXPERIMENTAL MODEL AND SUBJECT DETAILS

#### System Design and Function

Aquaria tank systems were maintained at three temperature regimes that compare to recorded seasonal ocean temperatures for Southern New England (SNE), Southern Maine (SME) and Northern Maine (NME) regions (Figure S8). Average monthly temperatures were obtained through NOAA’s National Oceanographic Data Center (NOAA, 2022).

The aquaria systems consisted of three individual 1400 L recirculating artificial sea water units with two 365 L tanks in each unit. For each system, water was mechanically and chemically filtered and passed through a bank of two 65 W ultraviolet sterilizers before flowing back into the lobster holding tanks, at a minimum of one full tank change per hour. Twenty-five percent of tank water was exchanged twice a week with freshly made and sterile artificial sea water. Temperature and dissolved oxygen in water was tested daily. Levels of ammonia, nitrogen, and pH were tested twice a week. Water was maintained at levels of total ammonia and nitrogen less than 1.0 mg/L; nitrite, less than 0.1 mg/L; pH from 8.0 to 8.1; 33–35 ppt salinity; and oxygen, > 7.0 mg/L. Initially, all tank systems were held at a temperature 11± 2 °C for a two-week adjustment/baseline period, as this was representative of the ocean water temperature from where lobsters had been sourced.

#### Lobsters

Fifty-seven female lobsters were collected during the Maine State ventless trap survey (Maine Department of Marine Resources, 2006) in late June 2016 from Maine’s lobster management zones F and G (42°05.5’ N, 70°14’ W) which stretch from the southern border of Maine to Small Point, (Figure S9). Non-shell-diseased lobsters (no apparent shell disease; n = 28) and shell diseased lobsters (apparent signs of ESD; n=29) were randomly distributed between the three tank systems, therefore, each system held a minimum of nine apparently non-shell-diseased (referred to as healthy) and nine shell-diseased lobsters (referred to as diseased). Healthy versus diseased lobsters were held in separate tanks within the system. Lobsters were individually compartmentalized within the tanks to reduce agonistic behavior and cannibalism, their claws were un-banded to allow for normal grooming, and they were fed previously frozen fish, shrimp, and green crabs throughout the study. Over the course of the study, 15 lobsters died.

Lobsters were acclimated for approximately three weeks before seasonal temperature regimes were initiated. The three systems were phased into current seasonal temperatures at no more than 2°C per day. Two weeks allowed for lobsters to acclimate to laboratory aquaria systems, eliminating stress response from capture and transport and allowed lobsters to come to a ‘resting’ state (Basti et al., 2010).

### METHOD DETAILS

#### Molt Stage Evaluation

The relative molt stage of lobsters was determined by estimating shell rigidity (Aiken 1980) and grouped into postmolt (stages A, B, C1 and C2); intermolt (stages C3 and C4); and premolt (stage D).

#### Shell Disease Evaluation

The extent of shell disease on lobsters was visually assessed (Figure S10) and scored using the shell disease index: 0, no observable signs of disease; 1+, shell disease signs on 1-10% of the shell surface; 2+, shell disease signs on 11-50% of the shell surface; 3+, shell disease signs on > 50% of the shell surface (Smolowitz et al., 2005).

#### Hemolymph Sampling and Plasma Antimicrobial Assay

Approximately 1.5 ml of hemolymph was collected aseptically from the dorsal abdominal artery into a sterile 3.0 cc syringe with a 23-gauge needle and then discharged into a 2 ml microfuge tube. Microfuge tubes were held on ice and centrifuged at 10,000g within 10 minutes of sample collection to avoid coagulation of the hemolymph. The plasma (cell-free hemolymph) was then transferred to a new microfuge tube. All plasma samples were stored at -80॰ C until the anti-microbial assay was performed.

The antimicrobial activity of the lobsters’ plasma was measured using a modified turbidimetric assay (Noga et al., 1994) where *Escherichia coli* D31 (Monner et al., 1971) is treated with plasma, and bacterial growth is compared among treatments after incubation. The bacterium *E. coli* D31 was acquired from Yale University’s bacterial culture collection. The stored frozen lobster plasma samples were thawed on ice and filter sterilized using a 0.22 µm filter prior to testing. The procedure was performed as described (Homerding et al., 2012) but was modified by performing the entire assay in a 96-well plate. For each plasma sample, 10 μl of the filter sterilized plasma was incubated with 10 μl of the prepared bacterial suspension and 30 μl of PBS for 30 minutes at room temperature. Next, 450 μl of cold Trypticase Soy Broth containing additional 1% NaCl and 0.1 mg/L streptomycin was added. Each plasma sample was then immediately plated in triplicate onto a 96-well plate (Whytrigg Close, Falcon Scientific, United Kingdom). Microbial growth was measured by absorbance at 570 nm on a BioTek spectrophotometer plate reader (BioTek, now Agilent Technologies, Santa Clara, California, U.S.) every 4 hour for 22 hours. Antimicrobial activity, the bacterial growth inhibition, of plasma was calculated as the mean optical density and compared to the *E. coli* D31 growth control without lobster plasma. Increasing optical densities (OD) indicate increasing bacterial growth.

#### Carapace Sampling for Microbial Analyses

Carapace sampling was performed by swabbing an approximate 4 cm^2^ square surface area of the dorsolateral region of the cephalothorax (Figure S11) with a sterile cotton-tipped applicator. Each lobster was sampled in the same anatomical location at each sampling timepoint. The right side of the dorsolateral area of the cephalothorax was sampled for the baseline sampling, the left side for the Time 1, and the right side again for Time 2. The applicator was repeatedly rolled over the approximate 4 cm^2^ square surface area for 30 seconds and then placed in a tube containing 0.9 ml of sterile artificial sea water (ASW) (Cold Spring Harbor Laboratory Press, 2012). These samples were held on ice and then processed for microbial community sampling.

#### Microbial Enumeration and Culture

The swab placed in 0.9 ml of sterile ASW was vortexed vigorously for 1 minute to dislodge cellular and microbial material. The swab was aseptically removed and discarded, and a subsample of the solution (400 μl) was set aside for DNA extraction. This sample containing the dislodged material was considered the undiluted sample for the microbial enumeration and culture assays. A serial dilution was performed out to 10^-3^ for the baseline and Time 1 and to 10^-5^ for Time 2 by serially transferring 100 μl into 900 μl of sterile ASW. All dilutions were plated on 2 media by adding 100 μl to each plate and spreading using a flame sterilized glass spreader. The two media used were Trypticase Soy Agar with 5% sheep red blood cells +1.5% NaCl (BA) (Northeast Laboratory, Waterville, Maine, USA), and a Marine Sea Salt Agar plate (MA), prepared in house with 10.0 g Trypticase Soy Agar (Becton, Dickinson and Co. Sparks, Maryland. USA), 20.0 g agar (Becton, Dickinson and Co. Sparks, Maryland, USA), 33.8 g instant ocean (Crystal Sea® Marinemix, Baltimore, MD), 1000 ml deionized water, and then autoclaved for sterility. All inoculated plates were incubated at 15±2 °C for 10 days. Ten days allowed for maximum growth of all bacterial types and allowed for a greater degree of colony morphology differentiation. Culturable microbial growth was enumerated by standard plate count methods and predominant colonies selected per media type for phenotypic identification using the Biolog® Microbial Identification System Gen II (BMIS) and Sanger 16S rRNA sequencing on selected isolates. The five most predominant colonies on each media type were selected for identification, resulting in 724 isolates across all samples. Isolated colonies were re-streaked onto the corresponding medium for re-isolation and purity and the isolates incubated again for 48 - 72 hours at 16° C. Preliminary biochemical profiling included Gram stain, oxidase, catalase reaction and carbohydrate utilization and H_2_S production with triple sugar iron (TSI) medium. Isolates were then filtered according to preliminary similar biochemical reactions prior to final identification using the BMIS. Those isolates that could not be identified by BMIS were identified using 16S rRNA sequencing.

#### DNA extraction

A 400 μl subsample of each of the ASW swab solutions as previously described was transferred to a clean 1.0 ml microcentrifuge tube. Samples were centrifuged at 10,000 rpm for 10 minutes at 4°C. Supernatant was discarded and the remaining cell pellets were frozen at -80°C in 500 μl 95% Molecular Grade Ethanol for DNA extraction.

DNA extraction was performed using the DNeasy Blood and Tissue kit (QIAGEN^®^, Hilden, Germany) with a modified pre-treatment lysis for Gram-Positive bacteria. Samples from the -80°C were placed on ice and ethanol was removed. Pellets were air dried for approximately 10 minutes and resuspended in 180 μl enzymatic lysis buffer (20mM Tris-Cl pH 8.0, 2 mM sodium EDTA, 1.2% Triton X-100 with 20 mg/ml lysozyme added immediately before use) and incubated for 2 hours at 37°C. A pre-treatment lysis protocol for Gram-Negative bacteria was then performed in which 25 μl Proteinase K and 200 μl Buffer AL were added and mixed by vortexing. Tubes were incubated at 56°C overnight for a minimum of 12 hours. The binding and washing steps were followed per manufacturer’s spin-column instructions for purification of total DNA from animal tissue. DNA was eluted in a final volume of 50 μl. Total genomic DNA was quantified and qualified using a NanoDrop^®^ 1000 spectrophotometer (NanoDrop Technologies, Wilmington, Delaware, USA).

#### Culture identification using whole 16S rRNA sequencing

Individual colonies cultured on agar as previously described were selected for 16S rRNA amplification and sequencing in 50 μl nuclease-free water and boiled at 99°C for 5 minutes to lyse cells and denature DNase. Samples were centrifuged at 12,000 x G for 1 minute to separate cellular debris and 10 μl of the supernatant was transferred to a 500 μl tube. PCR amplification was performed using Promega GoTaq^®^ Flexi DNA polymerase and PCR nucleotide mix (Promega, Madison, Wisconsin, USA) for final concentrations of 1.5 mM MgCl_2_, 0.2 μm of each dNTP, 0.5 μm forward primer (5’-TCCTACGGGAGGCAGCAGT-3’) and 0.5 μm reverse primer (5’-GGACTACCAGGGTATCTAATCC-3’) (Nadkarni et al., 2002). Samples were amplified on a BioMetra thermocycler with PCR conditions consisting of an initial denaturation for 2 minutes at 95°C followed by 35 cycles of denaturation for 1 minute at 95°C, annealing for 1 minute at 60°C, and extension for 45 seconds at 72°C and a final extension for 5 minutes at 72°C.

Gel electrophoresis was performed on 1.5% agarose in 1x TAE buffer at 85 volts for 40 minutes with 20 μl PCR product and 100bp ladder (Promega). QIAQuick Gel Purification & Extraction Kit (QIAGEN^®^) was used to purify PCR products per manufacturer’s instructions. Total genomic DNA was quantified and qualified using a NanoDrop^®^ 1000 spectrophotometer. Purified PCR products were prepared and submitted for sequencing performed by the University of Maine DNA Sequencing Facility using an ABI model 3730 DNA Sequencer. Sequences were trimmed using Geneious R10.0 software and identified using the NCBI BLAST database.

#### Bacterial community library preparation for 16S rRNA amplicon sequencing

A total of 131 lobster shell samples, 6 tank controls consisting of a 15 cm^2^ area swabbed from tank surfaces at the second time point, and 4 no-template negative controls were successfully sequenced across 4 sequencing runs. DNA extract was sent to Jonah Ventures, LLC (Boulder, Colorado, U.S), which conducted 16S rRNA gene PCR amplification, amplicon quantification and purification, Illumina MiSeq ver. 4 sequencing, and initial quality-control trimming, according to their proprietary standard operating procedures. Data preprocessing steps included demultiplexing sequence files using the barcode in the header, using python code generated by Joy-El Talbot at Iris Data Solutions (Supplemental materials, splitFASTQByHeader.py).

Data processing, analysis, and visualization was performed in RStudio using the R software platform versions 3.6.3 through 4.2.1 (RCoreTeam, 2022), using a modified version of the DADA2 pipeline (Callahan et al., 2016), available in Supplemental materials, DADA2_Ishaq_16S_lobster_shell_bacteria_example_code.R. Forward and reverse reads were processed in batches based on 4 total sequencing runs, for quality assurance steps through to amplicon sequence variant identification, before being compiled into a single dataset for rarefaction and statistical analysis. Raw reads and sample data, including about the sequencing run, lobsters, and environmental conditions, are publicly available through NCBI Sequence Read Archive (SRA), under BioProject Accession PRJNA887451.

For each of the four sequencing runs, the quality of forward and reverse reads (>900,000 paired read) was initially examined with quality scores plotted with DADA2, and forward/reverse sequences were processed together using filterAndTrim command parameters: trimming the first and last 10 bases, 2 max expected errors for forward reads and 3 for reverse reads, no ambiguous bases, and removing any reads which matched to the positive control, phiX. The DADA2 package in R was used to calculate error rates of the filtered output, dereplicate reads (combining identical reads), pick amplicon sequence variants (SVs) and make sequence tables, and remove chimeras using a *de novo* approach. The workflow was verified by tracking reads through the analysis with ggplot to see how many were lost at each QA step. Taxonomy was assigned to sequences with Silva ver. 138 taxonomy database (Pruesse et al., 2007) formatted for DADA2. Sequences which matched as eukaryotic mitochondria or chloroplast were removed using the dplyr package (Wickham et al., 2015).

A phyloseq object was created for each sequencing run with the resulting sequence variant table, taxonomy, and additional sample data, using the phyloseq package (McMurdie and Holmes, 2013). Initial data exploration using unweighted Jaccard distance and ordination plots showed no sample clustering by extraction batch, indicating no residual batch-effect of sequencing in sample bacterial profiles. The no-template negative control samples specific to each sequencing run were used to identify and remove additional contaminating sequences from their respective runs (Ishaq, 2017). The cleaned dataset was rarefied to the size of the smallest sample, yielding 5,808 sequences per sample.

### QUANTIFICATION AND STATISTICAL ANALYSIS

#### Growth and weight analysis

Baseline weights of non-shell diseased and shell diseased lobsters were compared using a *t* test. Growth of the lobsters during the experiment was assessed by comparing the lobster weight at the baseline sampling to weight at the third sampling using a paired *t*-test. In addition, the percentage growth for each lobster was calculated, then growth of non-shell diseased lobsters was compared to shell diseased lobsters. The total number of mortalities throughout the experiment from non-shell diseased and shell diseased lobsters were compared for each temperature regime (i.e., SNE, SME, and NME) using Fisher’s exact tests.

#### Antimicrobial assay analysis

Antimicrobial assay analyses were conducted using GraphPad Prism v. 7 (Dotmatics, 2022) and all analyses were conducted using SPSS v. 24.0 (IBM, 2021). Nonparametric tests were used where the data were not normally distributed. The mean optical density readings at 22 hours for each treatment were compared with ANOVAs.

#### Cultured isolate analysis

At the baseline sampling, the bacterial loads of non-shell diseased and shell diseased lobsters were compared with a *t* test. For each sampling period, the bacterial loads within the non-shell disease and shell disease were compared across timepoints (i.e., baseline, lower and higher seasonal temperatures) using Kruskal-Wallis tests. Then the bacterial loads between non-shell disease and shell disease lobsters at each temperature regime (i.e., SNE, SME, and NME) were compared with Mann-Whitney tests. Shannon-Weiner (H’) equitability index was used to assess the diversity in the bacterial communities for all treatment groups using the following formula, where N is the total number of species found and n_i_ is the number of individuals of one particular species found (Beisel and Moreteau, 1997): H’=-∑[(n_1_/N) ln(n_1_/N)]

#### Bacterial community library analysis for 16S rRNA amplicon sequencing data

Alpha diversity was calculated using observed SV richness, and Shannon evenness derived from Shannon Diversity. A Shapiro-Wilkes test was used to test if diversity data were normally distributed. Differences in observed richness and evenness between sample groups were tested using linear mixed models for single-factor comparisons, and generalized linear mixed models to incorporate fixed and random (date) effects, as well as nesting within tank or accounting for repeated measures of lobsters, with the lme4 package (Bates et al., 2015) and visualized using the ggplot package (Wickham, 2016). When considering experimental and host factors individually, bacterial richness on shells was not affected (linear regression, *p* > 0.05), by individual tank, or tank temperature at the time of sampling indicate that shell bacteria are not simply responding to water temperature. Tank temperature fluctuates during the experiment and across the geographic treatments, and different tanks are occasionally the same temperature at the same or different timepoints, thus it was important to consider the effect of temperature independent of other factors.

Principal Coordinate Analysis (PCoA) was performed to explore similarity of samples based on host species identity and host infection status. Calculations were made using two different sample similarity algorithms (Jaccard and Bray-Curtis) using the phyloseq package to calculate and visualize. Jaccard Similarity uses presence/absence of bacterial SV and can be interpreted as a measure of whether taxa membership of the original community has changed. Bray-Curtis uses presence/absence of bacterial SVs and their relative abundance, and can be interpreted as a measure of whether different taxa are present under new conditions or if the original taxa remain but at different levels of abundance. Sample clustering was tested with permutational ANOVA (i.e., permANOVA), with 1000 permutations per test, using the adonis function in the vegan package of R (Oksanen et al., 2020). The data subsets are specified in the results section.

Core taxa were identified using the microbiome package (Lahti and Shetty, 2020), designated as shared across 80% of lobsters in the subset group comparisons (healthy and diseased), and at least 0.001% abundance of the bacterial SV. The permutational random forest algorithm was used to identify important community features, i.e., bacterial SVs which were indicative of the community associated with a designated state, using the rfpermute package (Archer, 2022). Each random forest was performed with 500 trees and 100 permutations, resulting in an out-of-the-box error rate of the model from which accuracy may be calculated (1-OOB error), and a p-value for each taxon, corrected for multiple comparisons. The designated states (experimental factors), data subsets, and model accuracies are specified in the results section.

