## Supplemental Figures and Tables for "Warmer water temperature and epizootic shell disease reduces diversity but increases cultivability of bacteria on the shells of American Lobster (*Homarus americanus*)"

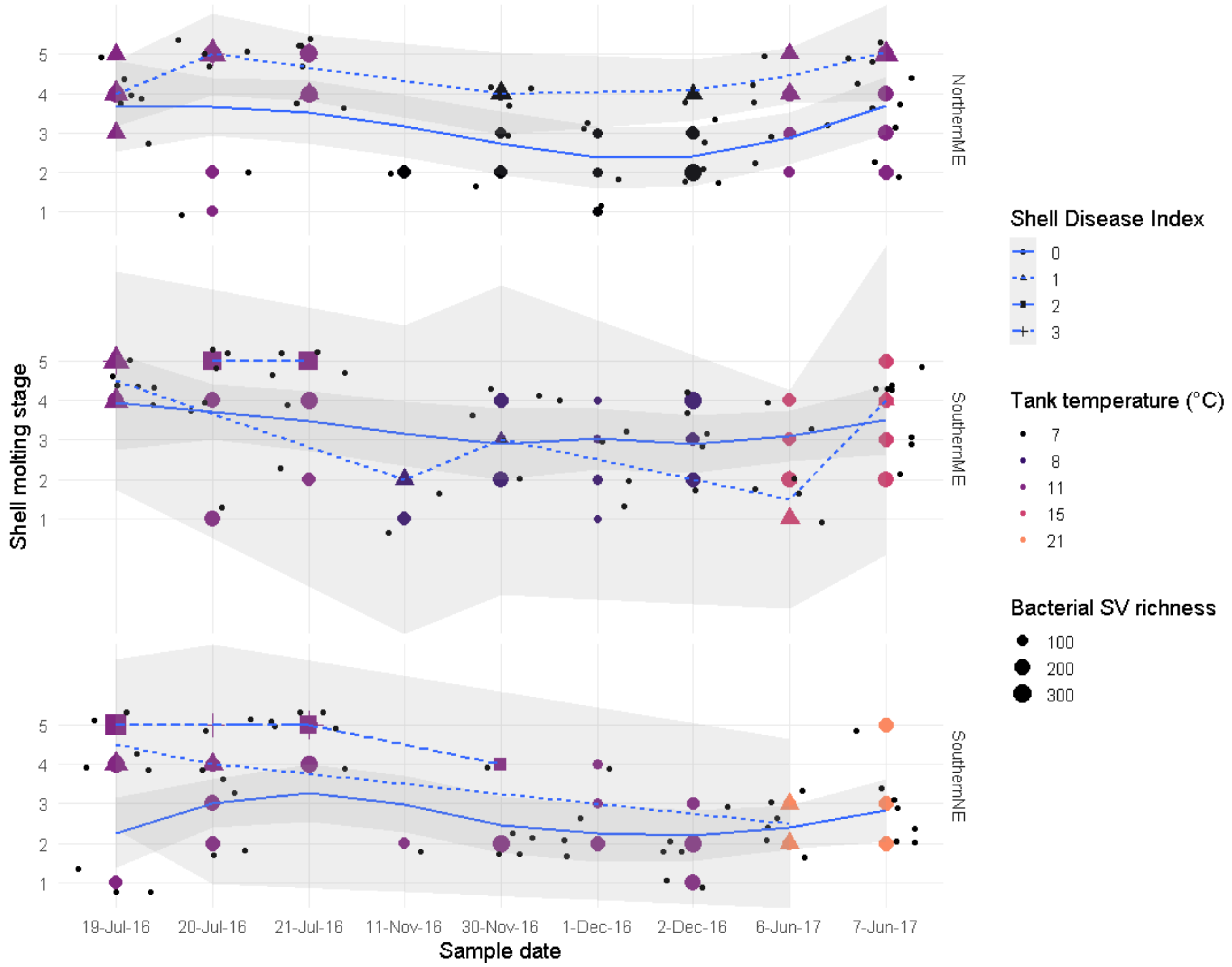


**Figure S1 Sequence variant richness for lobster shell bacterial communities at different shell molting stages under three simulated seasonal ocean temperatures over time.** Average seasonal ocean temperatures were simulated (i.e., tank temperature denoted by point color) for Northern Maine (ME), Southern Maine, and Southern New England (NE). Point size indicates bacterial richness (number of sequence variants).

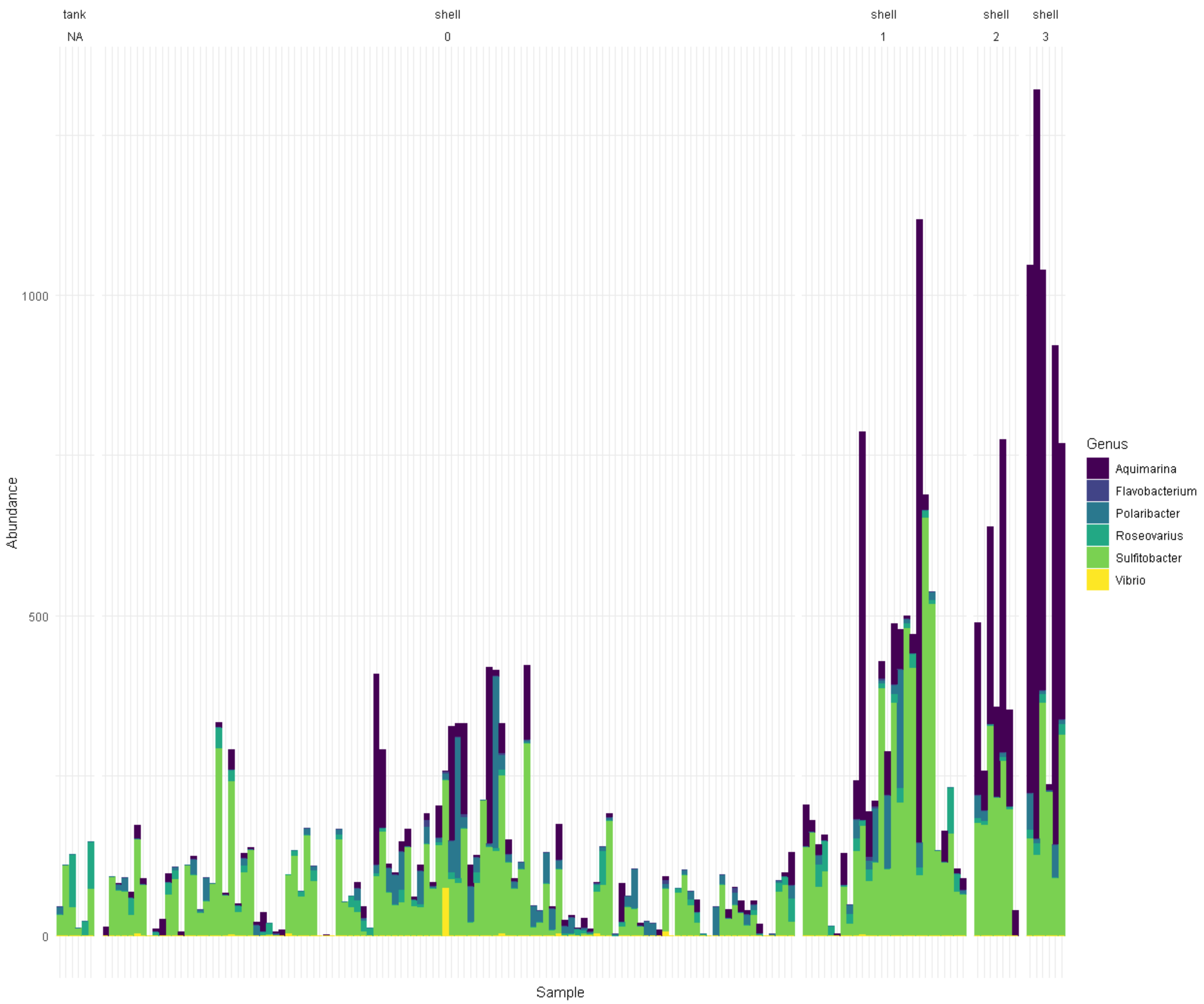


**Figure S2 Abundance (sequence variants) of putative pathogenic bacteria which have been implicated in epizootic shell disease, by Shell Disease Index.**

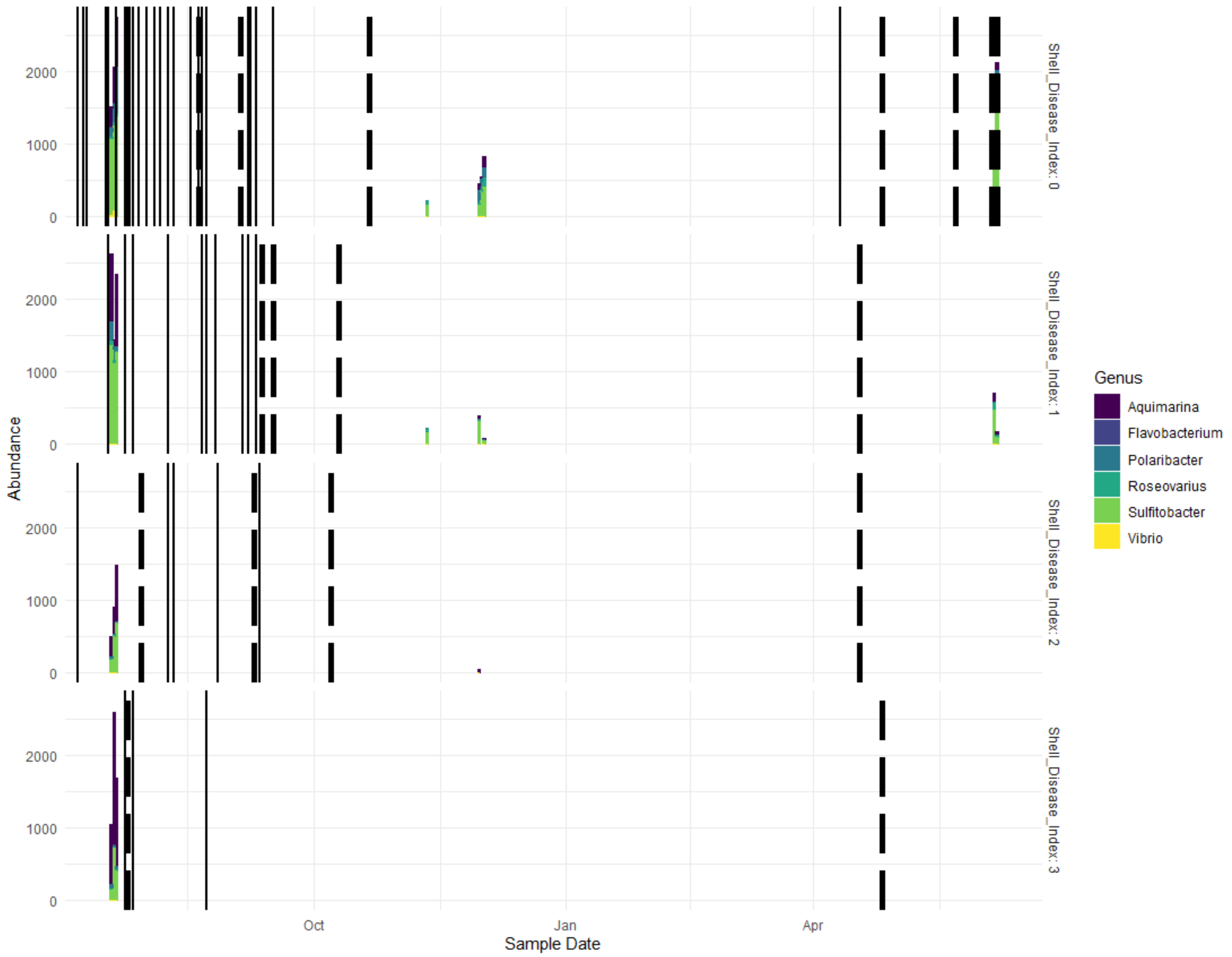


**Figure S3 Abundance (sequence variants) of putative pathogenic bacterial genera in relation to molting, mortality, and Shell Disease Index.** Color denotes the genus of putative pathogens. Sampling date is along the x-axis; solid lines denote when a lobster molted, and dashed lines denote when a lobster died. Vertical panels denote Shell Disease Index from apparently healthy (0), to visual disease >50% of the shell (3).

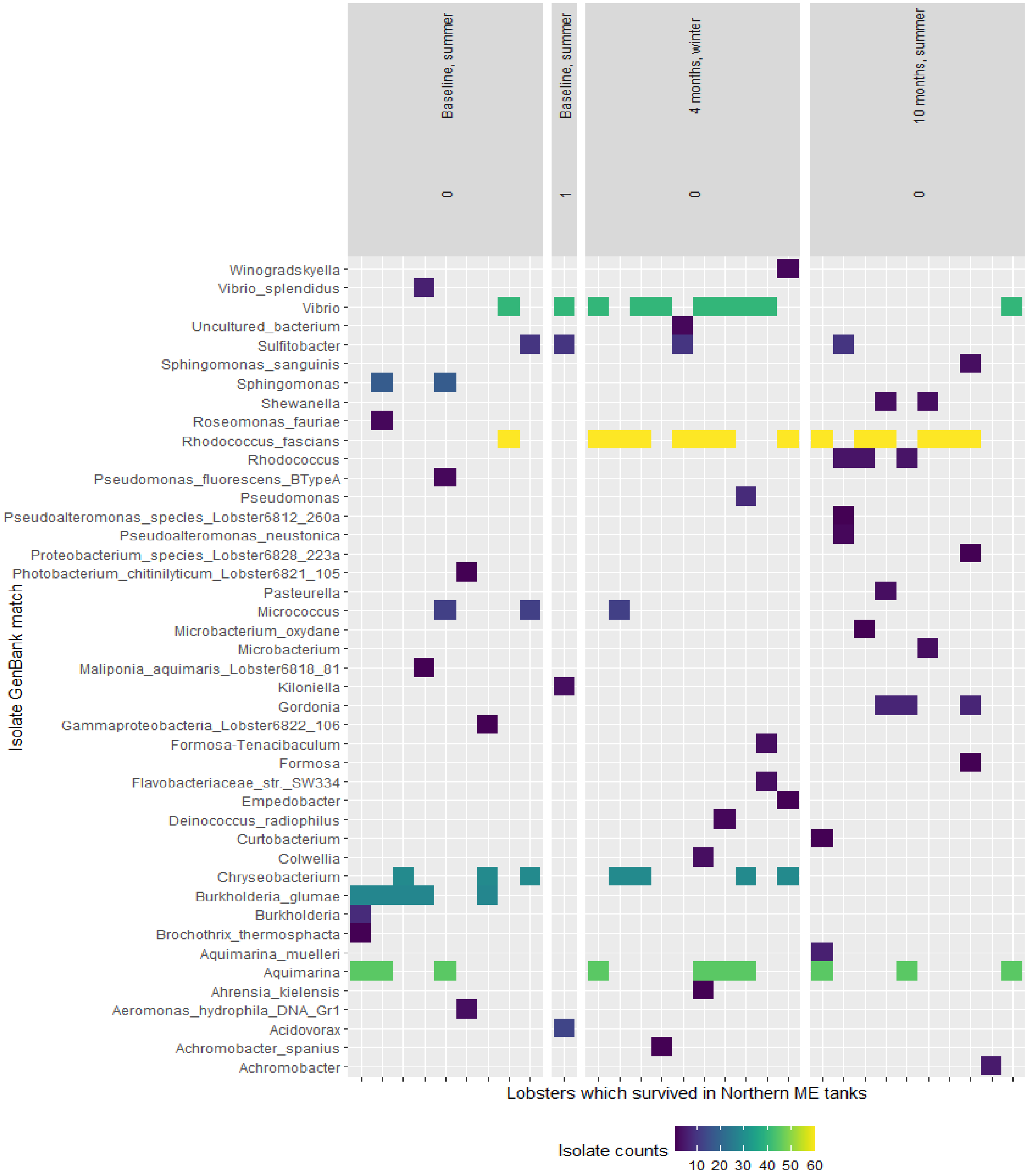


**Figure S4 Comparison of bacteria isolated from shells of lobsters which survived the experiment in the Northern ME tanks.** Bacterial isolates were identified using BMIS and 16S rRNA gene sequencing with the GenBank sequence reference database. Only isolates which could be identities are included. Columns are paneled by Timepoint and the Shell Disease Index ranking: 0, no observable disease; 1, disease on 1 - 10% of the shell; 2, disease on 11 - 50% of the shell; and 3, disease on >50% of the shell. GenBank match of “n/a” indicates isolate could not be identified at any the genus level of taxonomy.

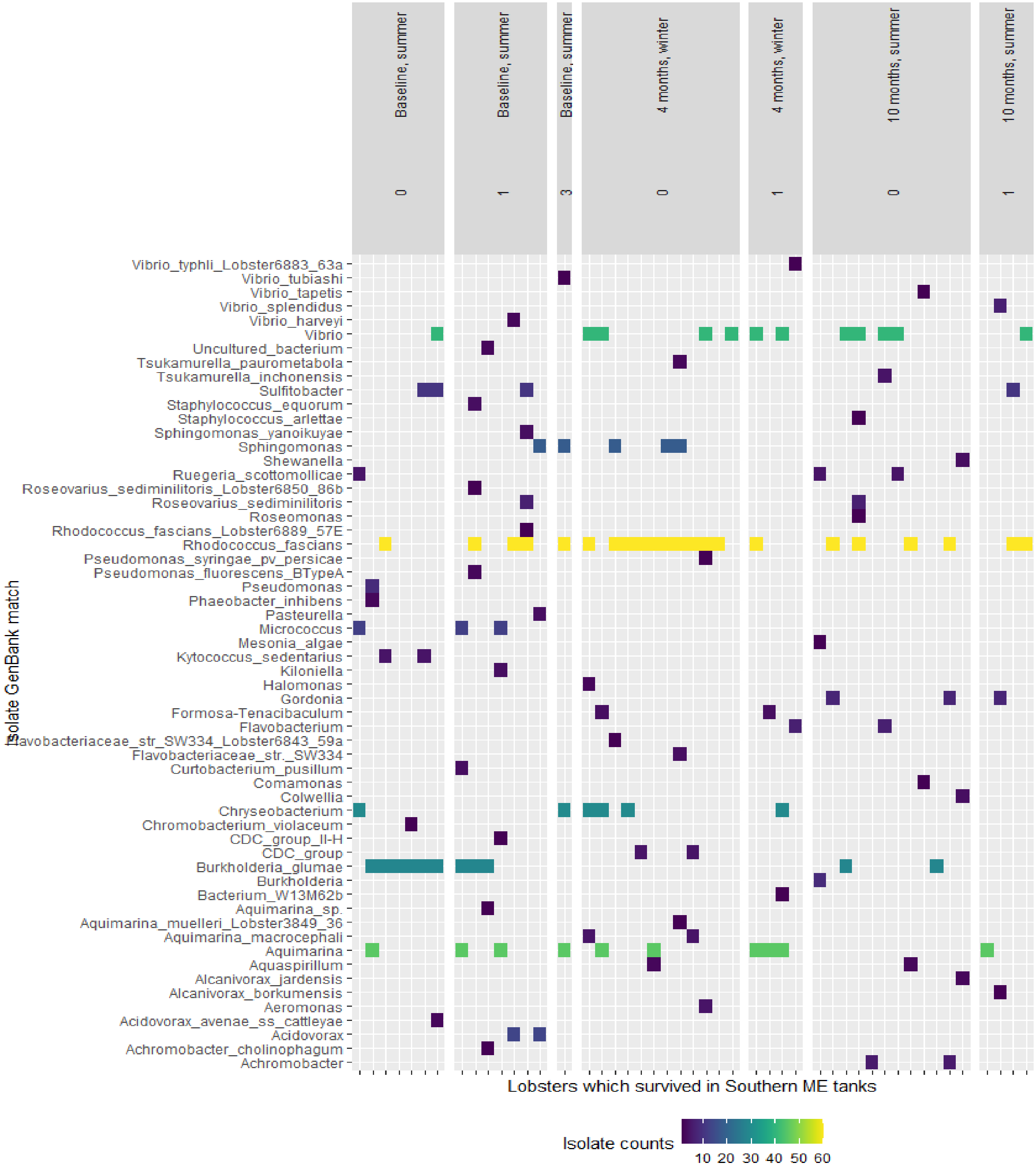


**Figure S5 Comparison of bacteria isolated from shells of lobsters which survived the experiment in the Southern ME tanks.** Bacterial isolates were identified using BMIS and 16S rRNA gene sequencing with the GenBank sequence reference database. Only isolates which could be identities are included. Columns are paneled by Timepoint and the Shell Disease Index ranking: 0, no observable disease; 1, disease on 1 - 10% of the shell; 2, disease on 11 - 50% of the shell; and 3, disease on >50% of the shell. GenBank match of “n/a” indicates isolate could not be identified at any the genus level of taxonomy.

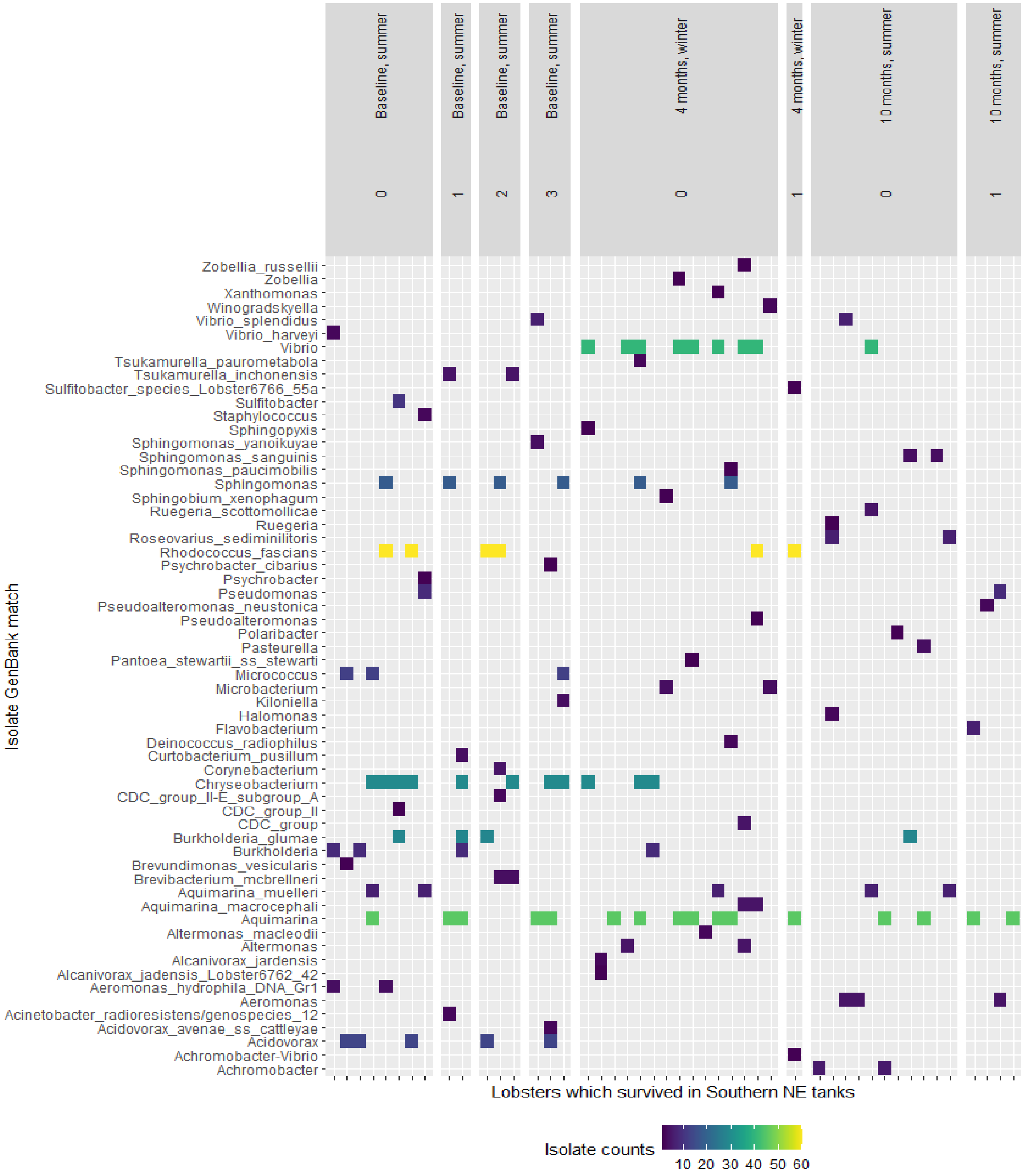


**Figure S6 Comparison of bacteria isolated from shells of lobsters which survived the experiment in the Southern NE tanks.** Bacterial isolates were identified using BMIS and 16S rRNA gene sequencing with the GenBank sequence reference database. Only isolates which could be identities are included. Columns are paneled by Timepoint and the Shell Disease Index ranking: 0, no observable disease; 1, disease on 1 - 10% of the shell; 2, disease on 11 - 50% of the shell; and 3, disease on >50% of the shell. GenBank match of “n/a” indicates isolate could not be identified at any the genus level of taxonomy.

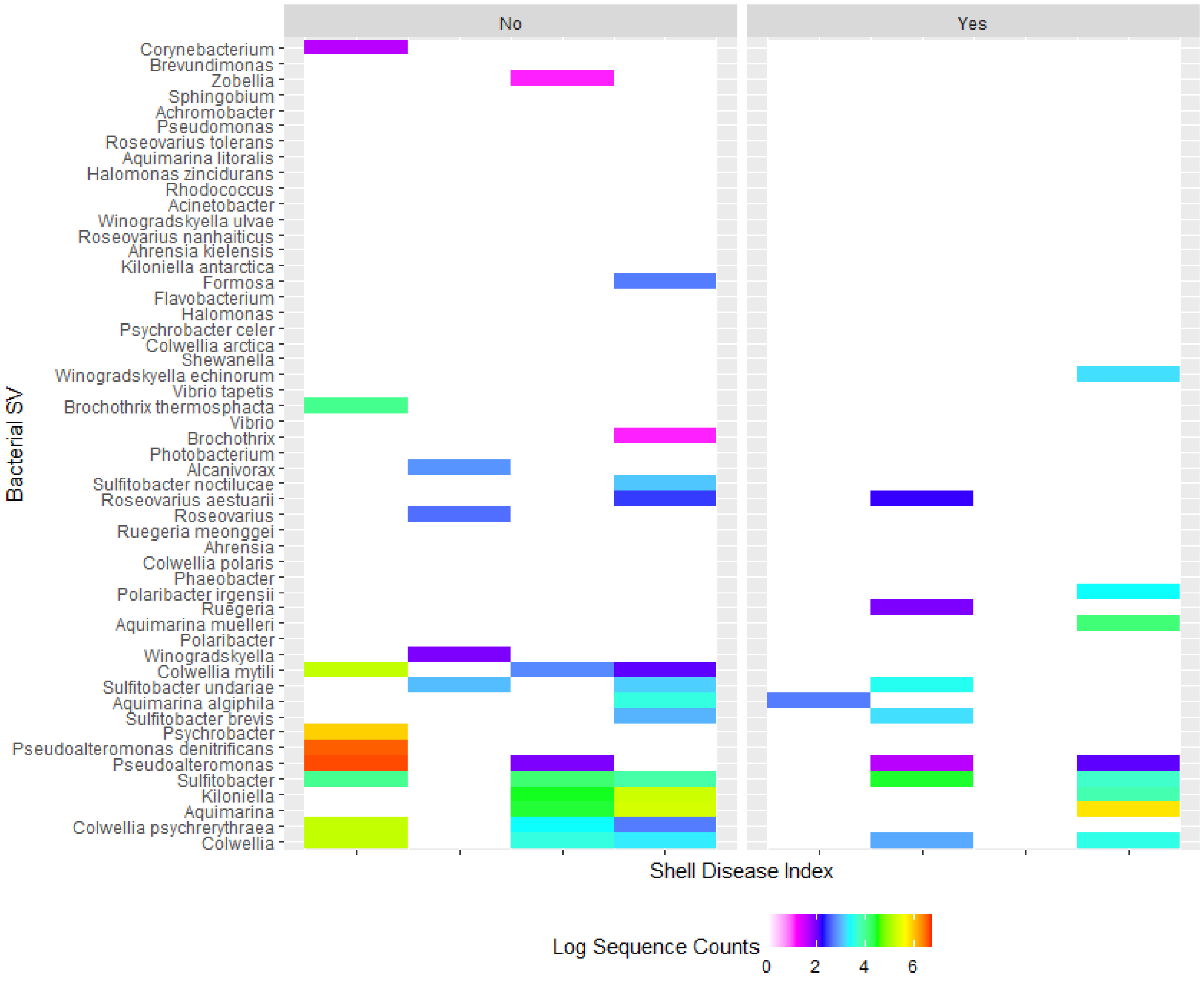


**Figure S7. Sequence read counts from bacterial SVs which matched the genus identity of the cultured bacterial isolates from lobster shells.** Color scale represents log-transformed sequence read counts. Columns are paneled by mortality (yes/no) and the Shell Disease Index ranking: 0, no observable disease; 1, disease on 1 - 10% of the shell; 2, disease on 11 - 50% of the shell; and 3, disease on >50% of the shell.

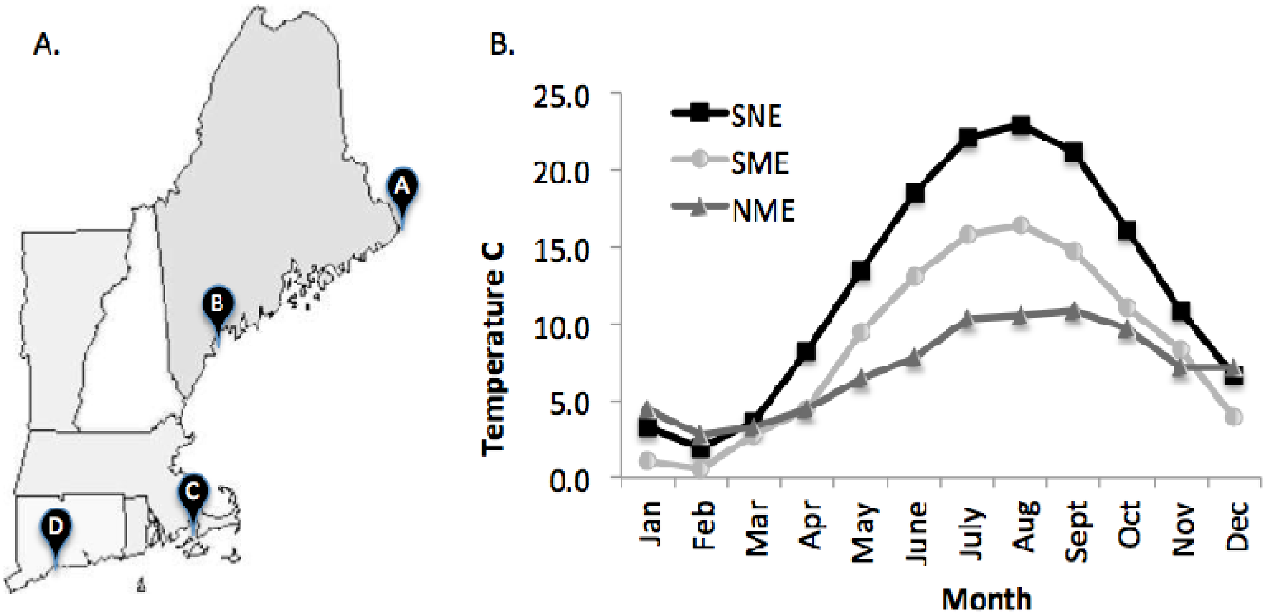


**Figure S8. Water temperature regimes**. A. Temperatures were obtained through the National Oceanographic Data Center (NODC). NODC temperatures reflect those recorded near Eastport, ME (A); Portland, ME (B); and an average of temperatures from Woods Hole, MA (C) and New Haven, CT (D) was used to represent Southern New England. B. Annual temperature cycles used in this project to represent Southern New England (SNE), Southern Maine (SME) and Northern Maine (NME).

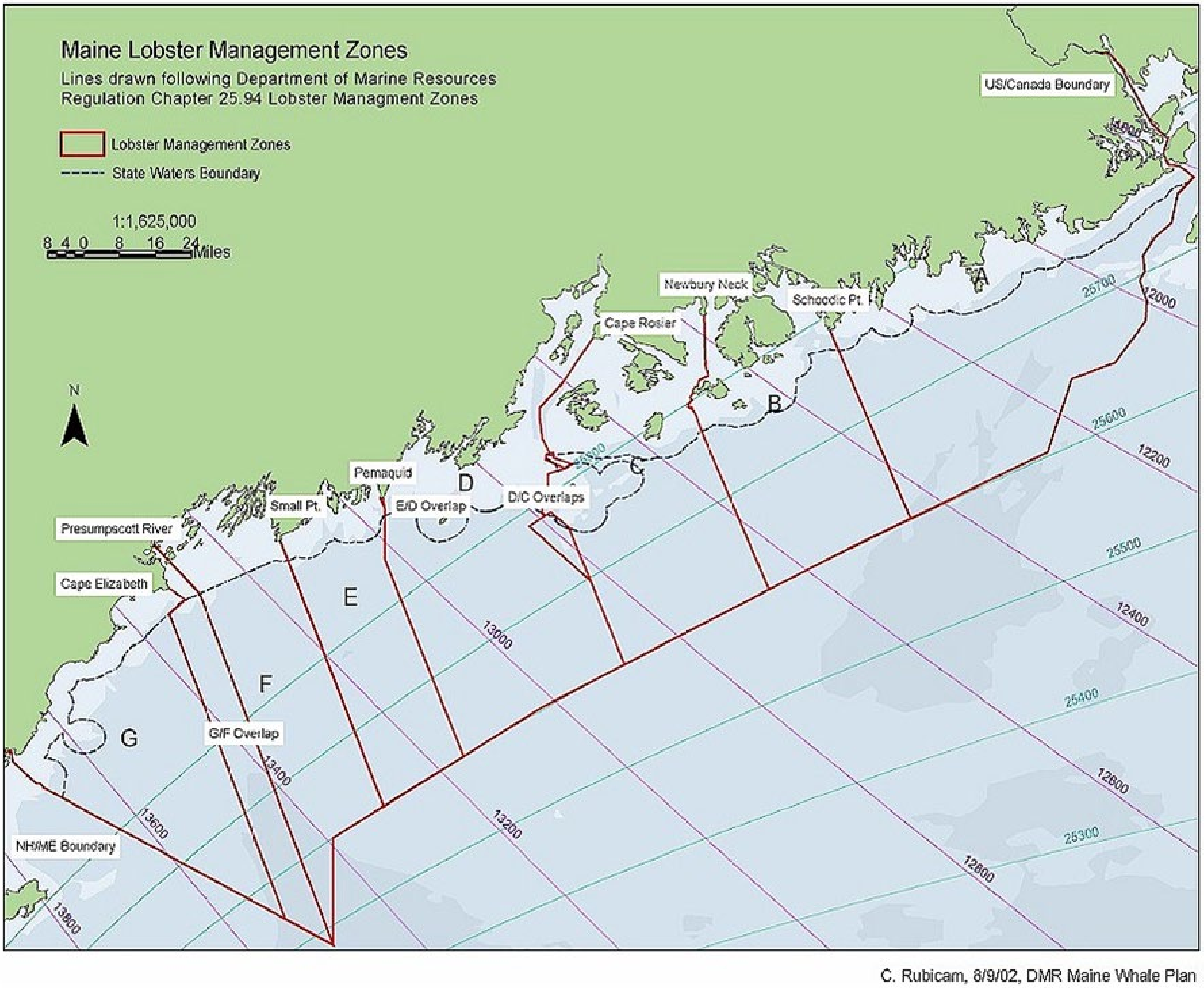


**Figure S9. Maine lobster management zones.** Lobsters for the study were trapped in management zones F and G.

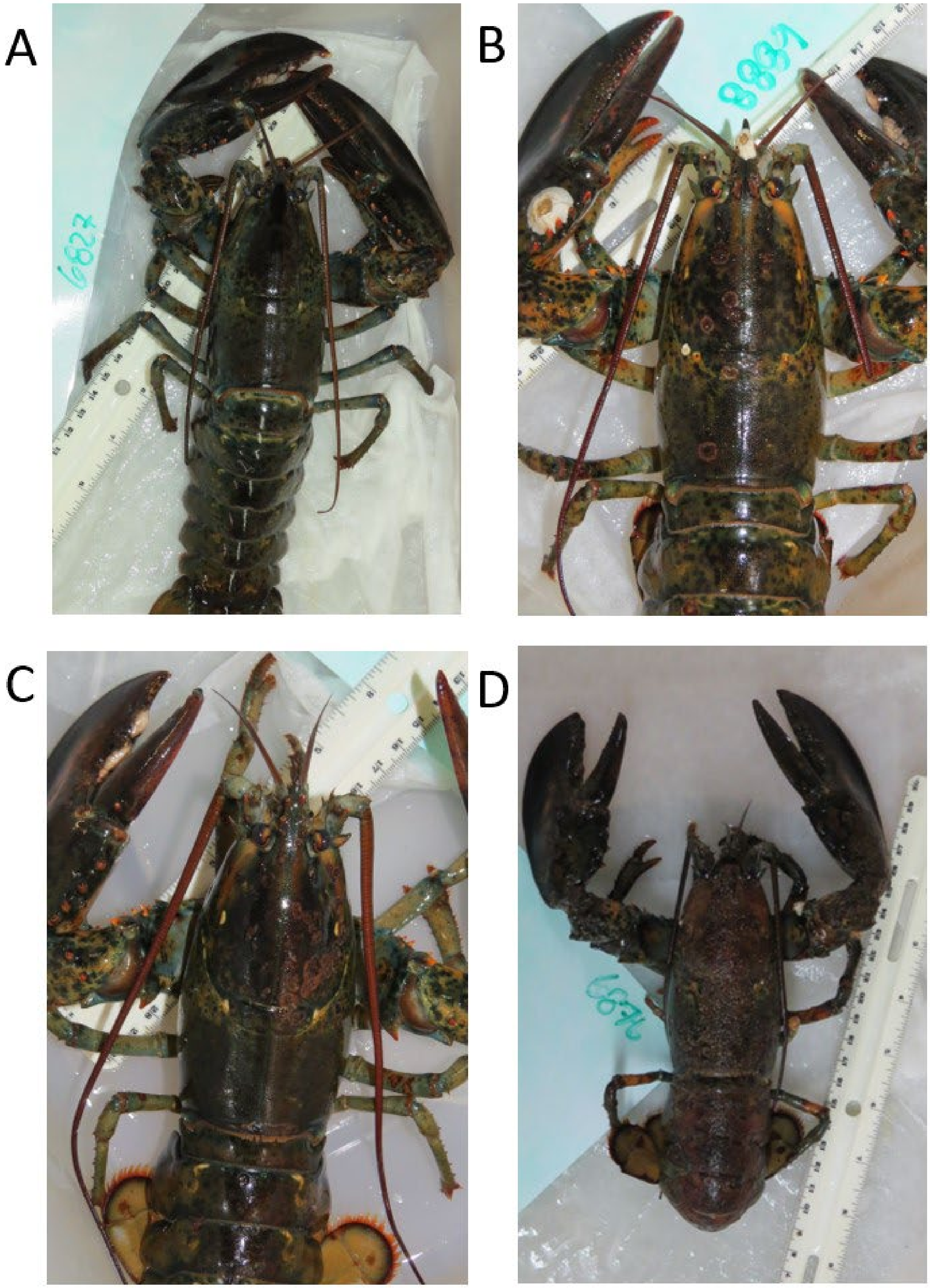


**Figure S10. Examples of lobster shell disease indices.** A) 0, no observable signs of disease, B) 1+, shell disease signs on 1-10% of the shell surface, C) 2+, shell disease signs on 11-50% of the shell surface, D) 3+, shell disease signs on > 50% of the shell surface.

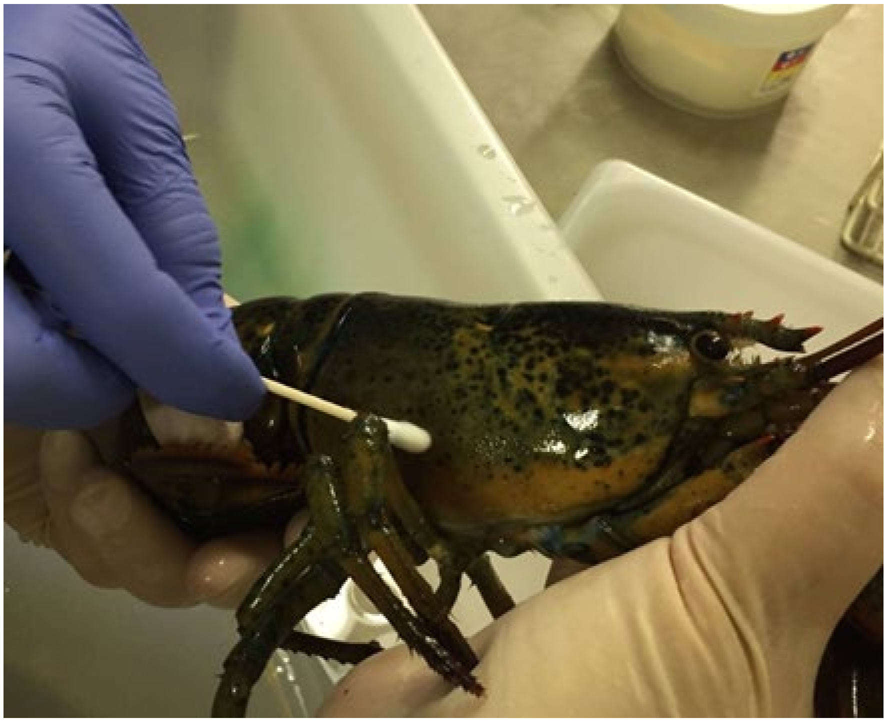

**Figure S11. Lobster carapace sampling using a sterile cotton swab to obtain bacterial communities from the shell surface**. The right side of the dorsolateral area of the cephalothorax was sampled for the baseline sampling, the left side for the Time 1, and the right side again for Time 2.

**Supplemental Tables**

**Table S1 Taxonomy of lobster shell bacterial isolates**

| Class | Order | Family | Genus |
|---|---|---|---|
| ***Actinomycetota (formerly Actinobacteria)*** | | | |
| *Actinobacteria* | *Actinomycetales* | *Gordoniaceae* | *Gordonia* |
| *Actinomycetales* | | *Intrasporangiaceae* | *Kytococcus* |
| | | *Tsukamurellaceae* | *Tsukamurella* |
| | *Corynebacterineae* | *Corynebacteriaceae* | *Corynebacterium* |
| | | *Nocardiaceae* | *Rhodococcus* |
| | *Micrococcales* | *Brevibacteriaceae* | *Brevibacterium* |
| | | *Micrococcaceae* | *Micrococcus* |
| | *Micrococcineae* | *Microbacteriaceae* | *Curtobacterium* |
| | | | *Microbacterium* |
| ***Bacillota (formerly Firmicutes)*** | | | |
| *Bacilli* | *Bacillales* | *Listeriaceae* | *Brochothrix* |
| | | *Staphylococcaceae* | *Staphylococcus* |
| ***Bacteroidota (formerly Bacteroidetes)*** | | | |
| *Flavobacteria* | *Flavobacteriales* | *Flavobacteriaceae* | *Aquimarina* |
| | | | *Chryseobacterium* |
| | | | *Empedobacter* |
| | | | *Flavobacterium* |
| | | | *Tenacibaculum* |
| | | | *Formosa* |
| | | | *Mesonia* |
| | | | *Polaribacter* |
| | | | *Winogradskyella* |
| | | | *Zobellia* |
| ***Deinococcus- Thermus*** | | | |

| Class | Order | Family | Genus |
|---|---|---|---|
| *Deinococci* | *Deinococcales* | *Deinococcalaceae* | *Deinococcus* |
| ***Pseudomonadota (formerly Proteobacteria)*** | | | |
| *Alphaproteobacteria* | *Caulobacterales* | *Caulobacteraceae* | *Brevundimonas* |
| | *Hyphomicrobiales* | *Ahrensiaceae* | *Ahrensia* |
| | | | *Neptunicoccus* |
| | | | *Phaeobacter* |
| | | | *Pukyongiella* |
| | | | *Roseovarius* |
| | | | *Ruegeria* |
| | | | *Sulfitobacter* |
| | *Rhodobacterales* | *Rhodobacteraceae* | *Maliponia* |
| | *Rhodospirillales* | *Acetobacteraceae* | *Roseomonas* |
| | | *Comamonadaceae* | *Acidovorax* |
| | | | *Comamonas* |
| | | *Kiloniellaceae* | *Kiloniella* |
| | *Sphingomonadales* | *Sphingomonadaceae* | *Sphingomonas* |
| | | | *Sphingopyxis* |
| | | | *Sphingobium* |
| *Betaproteobacteria* | *Burkholderiales* | *Burkholderiaceae* | *Burkholderia* |
| | *Neisseriales* | *Neisseriaceae* | *Aquaspirillum* |
| | | | *Chromobacterium* |
| *Gammaproteobacteria* | *Aeromonadales* | *Aeromonadaceae* | *Aeromonas* |
| | *Alteromonadales* | *Alteromonadaceae* | *Achromobacter* |
| | | | *Alteromonas* |
| | | | *Pseudoalteromonas* |
| | | *Idiomarinaceae* | *Idiomarina* |
| | | *Colwelliaceae* | *Colwellia* |

| Class | Order | Family | Genus |
|---|---|---|---|
| | | *Shewanellaceae* | *Shewanella* |
| | *Enterobacterales* | *Erwiniaceae* | *Pantoea* |
| | *Pasteurellales* | *Pasteurellaceae* | *Pasteurella* |
| | *Oceanospirillales* | *Alcanivoracaceae* | *Alcanivorax* |
| | | *Halomonadaceae* | *Halomonas* |
| | *Pseudomonadales* | *Moraxellaceae* | *Acinetobacter* |
| | | | *Psychrobacter* |
| | | *Pseudomonadaceae* | *Pseudomonas* |
| | *Spongibacteriales* | *Sphingobacteriaceae* | *Spongibacter* |
| | *Vibrionales* | *Vibrionaceae* | *Photobacterium* |
| | | | *Proteobacterium* |
| | | | *Vibrio* |
| | *Xanthomonadales* | *Xanthomonadaceae* | *Xanthomonas* |
