## Supplementary material for "Warmer water temperature and epizootic shell disease reduces diversity but increases cultivability of bacteria on the shells of American Lobster (*Homarus americanus*)": R script

## Introduction and preliminary processing -----------------------------
# This script processes 16S rRNA amplicon sequencing data, for the lobster tank dataset generated by Deborah Bouchard et al. at the University of Maine.
# This workflow developed by Sue Ishaq, spring 2020 - fall 2022. There are 163 samples, including negative controls, across 4 sequencing runs.

## Preprocessing steps included demultiplexing sequence files using the barcode in the header, using python code generated by Joy-El Talbot at Iris Data Solutions.
## the "undetermined" files were not parsed because sequences contained at least 1 ambiguous base (N), other errors


## Install packages into R -----------------------------
install.packages("BiocManager") # installation manager package to help you download/install other packages
BiocManager::install("dada2") #the data analysis program, more info here: https://benjjneb.github.io/dada2/dada-installation.html
BiocManager::install("GenomeInfoDbData") 
BiocManager::install('phyloseq') #the data interpretation program
install.packages("vegan") # statistical package
install.packages("ape") # reading, writing, plotting, and manipulating phylogenetic trees
install.packages('plyr')
install.packages("dplyr") # data manipulation 
install.packages("lme4") # linear mixed models
install.packages("lmerTest") # and permutational capacity to go along with it
install.packages("emmeans")
install.packages("permute") # permutational capacity
install.packages("randomForest") # random forest feature selection
install.packages("rfPermute") # random forest with permutational capacity
install.packages("ggplot2") # graphics package 
install.packages("RColorBrewer") # graphics color customization
install.packages("ggpubr")
install.packages("viridis")

## Set directory and load packages EACH SESSION ----------------------------------------------
#setwd('~/Lobster_16S/') #CHANGE ME

library(dada2); packageVersion('dada2')
library(GenomeInfoDbData)
library(phyloseq); packageVersion("phyloseq")
library(vegan)
library(ape); packageVersion("ape")
library(plyr)
library(dplyr)
library(lme4)
library(lmerTest)
library(emmeans)
library(permute)
library(randomForest)
library(rfPermute)
library(ggplot2)
library(RColorBrewer)
library(ggpubr)
library(viridis)


## Load raw data ---------------------------

# set the folder paths to each of the locations of the raw sequencing files
JV31_raw_F <- '~/JV31/JV31_forward_raw/' #CHANGE ME
JV31_raw_R <- '~/JV31/JV31_reverse_raw/' #CHANGE ME

JV35_raw_F <- '~/JV35/JV35_forward_raw/' #CHANGE ME
JV35_raw_R <- '~/JV35/JV35_reverse_raw/' #CHANGE ME

JV39_raw_F <- '~/JV39/JV39_forward_raw/' #CHANGE ME
JV39_raw_R <- '~/JV39/JV39_reverse_raw/' #CHANGE ME

JV41_raw_F <- '~/JV41/JV41_forward_raw/' #CHANGE ME
JV41_raw_R <- '~/JV41/JV41_reverse_raw/' #CHANGE ME


## specify what the file names (fastqs) are to look for, this looks for only zipped (gz) files
JV31fnsF <- list.files(JV31_raw_F)
JV31fastqsF <- JV31fnsF[grepl('.gz$', JV31fnsF)] # CHANGE if different file extensions

JV31fnsR <- list.files(JV31_raw_R)
JV31fastqsR <- JV31fnsR[grepl('.gz$', JV31fnsR)] # CHANGE if different file extensions


JV35fnsF <- list.files(JV35_raw_F)
JV35fastqsF <- JV35fnsF[grepl('.gz$', JV35fnsF)] # CHANGE if different file extensions

JV35fnsR <- list.files(JV35_raw_R)
JV35fastqsR <- JV35fnsR[grepl('.gz$', JV35fnsR)] # CHANGE if different file extensions


JV39fnsF <- list.files(JV39_raw_F)
JV39fastqsF <- JV39fnsF[grepl('.gz$', JV39fnsF)] # CHANGE if different file extensions

JV39fnsR <- list.files(JV39_raw_R)
JV39fastqsR <- JV39fnsR[grepl('.gz$', JV39fnsR)] # CHANGE if different file extensions


JV41fnsF <- list.files(JV41_raw_F)
JV41fastqsF <- JV41fnsF[grepl('.gz$', JV41fnsF)] # CHANGE if different file extensions

JV41fnsR <- list.files(JV41_raw_R)
JV41fastqsR <- JV41fnsR[grepl('.gz$', JV41fnsR)] # CHANGE if different file extensions


## fully specify the path, and file names (fns) for the reads.  
JV31fnsF <- file.path(JV31_raw_F, JV31fastqsF)
JV31fnsR <- file.path(JV31_raw_R, JV31fastqsR)

JV35fnsF <- file.path(JV35_raw_F, JV35fastqsF)
JV35fnsR <- file.path(JV35_raw_R, JV35fastqsR)

JV39fnsF <- file.path(JV39_raw_F, JV39fastqsF)
JV39fnsR <- file.path(JV39_raw_R, JV39fastqsR)

JV41fnsF <- file.path(JV41_raw_F, JV41fastqsF)
JV41fnsR <- file.path(JV41_raw_R, JV41fastqsR)


# pull the sequence names using the forward sequenes and take the sequencing name chunk
  # Assumes filename = samplename_XXX.fastq.gz, splits the string based on the first "_" it finds, and takes chunk 1
  # example, take the back section: sample.names <- sapply(strsplit(basename(fastqsF), '_'), `[`, 2)
  # example, take the middle section: sample.names <- sapply(strsplit(basename(fastqsF), '_', "end.fastq"), `[`, 2)
JV31sample.names <- sapply(strsplit(basename(JV31fastqsF), '_'), `[`, 1) 
JV35sample.names <- sapply(strsplit(basename(JV35fastqsF), '_'), `[`, 1) 
JV39sample.names <- sapply(strsplit(basename(JV39fastqsF), '_'), `[`, 1) 
JV41sample.names <- sapply(strsplit(basename(JV41fastqsF), '_'), `[`, 1) 

# Assign the sample names to the list of Forward/R1 fastq files
names(JV31fastqsF) <- JV31sample.names 
names(JV35fastqsF) <- JV35sample.names 
names(JV39fastqsF) <- JV39sample.names 
names(JV41fastqsF) <- JV41sample.names 

## Check sequence quality ---------------------------
# Typically in paired-end datasets, R1 looks great and R2 less so. It may be better to use only R1 rather than merge and trim off a lot of R2.
# Peek inside one or two files to see what the data looks like instead.  This is helpful for understanding sequence file formatting and how to search/manipulate info within.

# Look at quality profile plot for aggregated F/RR1 or R/R2 reads for each sequencing run. 
  # This will show the quality scores for every base in the amplicon for all the sequences. Quality score is the sequencer's confidence score in that base call, and corresponds to how many errors can be expected. >30 is good, < 20 and you should trim it out (next section)
  # The green line is the average quality score for all sequences in that sample, black/grey are relative abundance of that score at each base.

plotQualityProfile(JV31fnsF, aggregate=T) # save this plot for future reference
plotQualityProfile(JV31fnsR, aggregate=T) # save this plot for future reference

plotQualityProfile(JV35fnsF, aggregate=T) # save this plot for future reference
plotQualityProfile(JV35fnsR, aggregate=T) # save this plot for future reference

plotQualityProfile(JV39fnsF, aggregate=T) # save this plot for future reference
plotQualityProfile(JV39fnsR, aggregate=T) # save this plot for future reference

plotQualityProfile(JV41fnsF, aggregate=T) # save this plot for future reference
plotQualityProfile(JV41fnsR, aggregate=T) # save this plot for future reference


## Filter and trim raw sequences ---------------------------
# Filer your sequences and trim sequence length based on various parameters.  There are two ways to do this, on the Forward, R1 reads only (they tend to be better quality than Reverse, R2s), or you can do it on Forward/Reverse at the same time with the intent to merge them together into contigs later.

# create folders for your filtered data, only do this once (unless you delete the folders).  You'll need to make one set for each dataset to facilitate a DADA2 function we'll do later
dir.create('~/JV31/JV31_forward_filtered/') #CHANGE ME
dir.create('~/JV31/JV31_reverse_filtered/') #CHANGE ME

dir.create('~/JV35/JV35_forward_filtered/') #CHANGE ME
dir.create('~/JV35/JV35_reverse_filtered/') #CHANGE ME

dir.create('~/JV39/JV39_forward_filtered/') #CHANGE ME
dir.create('~/JV39/JV39_reverse_filtered/') #CHANGE ME

dir.create('~/JV41/JV41_forward_filtered/') #CHANGE ME
dir.create('~/JV41/JV41_reverse_filtered/') #CHANGE ME


# Specify the folder path to place filtered files in a filtered subdirectory
JV31_filt_F <- '~/JV31/JV31_forward_filtered' #CHANGE ME
JV31_filt_R <- '~/JV31/JV31_reverse_filtered' #CHANGE ME

JV35_filt_F <- '~/JV35/JV35_forward_filtered' #CHANGE ME
JV35_filt_R <- '~/JV35/JV35_reverse_filtered' #CHANGE ME

JV39_filt_F <- '~/JV39/JV39_forward_filtered' #CHANGE ME
JV39_filt_R <- '~/JV39/JV39_reverse_filtered' #CHANGE ME

JV41_filt_F <- '~/JV41/JV41_forward_filtered' #CHANGE ME
JV41_filt_R <- '~/JV41/JV41_reverse_filtered' #CHANGE ME


# specify where to send the filtered files, and name them "samplename_F_filt.fastq.qz
JV31filtFs <- file.path(JV31_filt_F, paste0(JV31sample.names, "_F_filt.fastq.gz"))
JV31filtRs <- file.path(JV31_filt_R, paste0(JV31sample.names, "_R_filt.fastq.gz"))

JV35filtFs <- file.path(JV35_filt_F, paste0(JV35sample.names, "_F_filt.fastq.gz"))
JV35filtRs <- file.path(JV35_filt_R, paste0(JV35sample.names, "_R_filt.fastq.gz"))

JV39filtFs <- file.path(JV39_filt_F, paste0(JV39sample.names, "_F_filt.fastq.gz"))
JV39filtRs <- file.path(JV39_filt_R, paste0(JV39sample.names, "_R_filt.fastq.gz"))

JV41filtFs <- file.path(JV41_filt_F, paste0(JV41sample.names, "_F_filt.fastq.gz"))
JV41filtRs <- file.path(JV41_filt_R, paste0(JV41sample.names, "_R_filt.fastq.gz"))


# associate the same names with the filtered files
names(JV31filtFs) <- JV31sample.names
names(JV31filtRs) <- JV31sample.names

names(JV35filtFs) <- JV35sample.names
names(JV35filtRs) <- JV35sample.names

names(JV39filtFs) <- JV39sample.names
names(JV39filtRs) <- JV39sample.names
	
names(JV41filtFs) <- JV41sample.names
names(JV41filtRs) <- JV41sample.names

  
# Filter and trim Forward/Reverse for the use of contigs filter that sample pair and output as new same names	  
    filtoutputJV31 <- filterAndTrim( #filter command
      JV31fnsF, JV31filtFs, JV31fnsR, JV31filtRs,
	  	trimLeft=c(10, 10), # cuts off the first XX bases from the F reads (tend to be low quality). 10 for illumina, 15 for 454 pyrosequencing
	  	trimRight=c(10,10), # cuts off last XX bases of F and R reads (tend to be low quality).  If have 454 data, can instead use #maxLen=XXX
	  	maxEE=c(2,3), ## The maxEEparameter sets the maximum number of expected errors allowed in a read.
	  	maxN= 0, # max number of ambiguous bases (N) allowed, DADA2 doesn't allow any!!
	  	rm.phix=TRUE, # remove any PhiX DNA (used as positive control)
	  	verbose=TRUE
	  	) # verbose = print to screen
    
dim(filtoutputJV31) ## check the dimensions of the variable you created, will tell you how many samples it ran to double check you got them all

# take a look
filtoutputJV31

saveRDS(filtoutputJV31, "filtered_output_JV31.RDS") # save the filtered variable.  your filtered files are already saved, this saves the R item to work with later without filtering again so you can track read numbers at each stem
  
#---- repeat with JV35
filtoutputJV35 <- filterAndTrim( #filter command
  JV35fnsF, JV35filtFs, JV35fnsR, JV35filtRs,
  trimLeft=c(10, 10), # cuts off the first XX bases from the F reads (tend to be low quality). 10 for illumina, 15 for 454 pyrosequencing
  trimRight=c(10,10), # cuts off last XX bases of F and R reads (tend to be low quality).  If have 454 data, can instead use #maxLen=XXX
  maxEE=c(2,3), ## The maxEEparameter sets the maximum number of expected errors allowed in a read.
  maxN= 0, # max number of ambiguous bases (N) allowed, DADA2 doesn't allow any!!
  rm.phix=TRUE, # remove any PhiX DNA (used as positive control)
  verbose=TRUE
) # verbose = print to screen

dim(filtoutputJV35) ## check the dimensions of the variable you created, will tell you how many samples it ran to double check you got them all

# take a look
filtoutputJV35

saveRDS(filtoutputJV35, "filtered_output_JV35.RDS") # save the filtered variable.  your filtered files are already saved, this saves the R item to work with later without filtering again so you can track read numbers at each stem


#--- repeat with JV39
filtoutputJV39 <- filterAndTrim( #filter command
  JV39fnsF, JV39filtFs, JV39fnsR, JV39filtRs,
  trimLeft=c(10, 10), # cuts off the first XX bases from the F reads (tend to be low quality). 10 for illumina, 15 for 454 pyrosequencing
  trimRight=c(10,10), # cuts off last XX bases of F and R reads (tend to be low quality).  If have 454 data, can instead use #maxLen=XXX
  maxEE=c(2,3), ## The maxEEparameter sets the maximum number of expected errors allowed in a read.
  maxN= 0, # max number of ambiguous bases (N) allowed, DADA2 doesn't allow any!!
  rm.phix=TRUE, # remove any PhiX DNA (used as positive control)
  verbose=TRUE
) # verbose = print to screen

dim(filtoutputJV39) ## check the dimensions of the variable you created, will tell you how many samples it ran to double check you got them all

# take a look
filtoutputJV39

saveRDS(filtoutputJV39, "filtered_output_JV39.RDS") # save the filtered variable.  your filtered files are already saved, this saves the R item to work with later without filtering again so you can track read numbers at each stem


#--- repeat with JV41
filtoutputJV41 <- filterAndTrim( #filter command
  JV41fnsF, JV41filtFs, JV41fnsR, JV41filtRs,
  trimLeft=c(10, 10), # cuts off the first XX bases from the F reads (tend to be low quality). 10 for illumina, 15 for 454 pyrosequencing
  trimRight=c(10,10), # cuts off last XX bases of F and R reads (tend to be low quality).  If have 454 data, can instead use #maxLen=XXX
  maxEE=c(2,3), ## The maxEEparameter sets the maximum number of expected errors allowed in a read.
  maxN= 0, # max number of ambiguous bases (N) allowed, DADA2 doesn't allow any!!
  rm.phix=TRUE, # remove any PhiX DNA (used as positive control)
  verbose=TRUE
) # verbose = print to screen

dim(filtoutputJV41) ## check the dimensions of the variable you created, will tell you how many samples it ran to double check you got them all

# take a look
filtoutputJV41

saveRDS(filtoutputJV41, "filtered_output_JV41.RDS") # save the filtered variable.  your filtered files are already saved, this saves the R item to work with later without filtering again so you can track read numbers at each stem


## DADA2 learn error rates for JV31 ----------------------------
# DADA2 learns error rates for each sequencing run separately, because some error is inherent to the run.  Uses a random subset of bases from the data, DADA2 pipeline recommends learning the error rates prior to dereplication (combining identical sequences into one).

set.seed(1234) # set the seed for using random number generators.  setting a seed means it can be reproduced because you start in the same place.

# If starting a new session, specify the folder path to place filtered files in a filtered sub directory
JV31_filt_F <- '~/JV31/JV31_forward_filtered' #CHANGE ME
JV31_filt_R <- '~/JV31/JV31_reverse_filtered' #CHANGE ME


## set the file names for Forwards
JV31fnsF <- list.files(JV31_filt_F, full.names = TRUE)
JV31filtsF <- JV31fnsF[grepl('.gz$', JV31fnsF)] # CHANGE if different file extensions
sample.namesJV31F <- sapply(strsplit(basename(JV31filtsF), '_'), `[`, 1) # Assumes filename = samplename_XXX.fastq
names(JV31filtsF) <- sample.namesJV31F 

## set the file names for Reverses
JV31fnsR <- list.files(JV31_filt_R, full.names = TRUE)
JV31filtsR <- JV31fnsR[grepl('.gz$', JV31fnsR)] # CHANGE if different file extensions
sample.namesJV31R <- sapply(strsplit(basename(JV31filtsR), '_'), `[`, 1) # Assumes filename = samplename_XXX.fastq
names(JV31filtsR) <- sample.namesJV31R

# dereplicate, which compresses identical sequences into one to make computing faster.  DADA2 will retain the quality info for all seqs that were compressed. 
derepJV31F <- derepFastq(JV31filtsF, verbose=TRUE)
derepJV31R <- derepFastq(JV31filtsR, verbose=TRUE)


## learn the error rates from your sequencing run. Some error is inherant to sequencing run, so this should be done on each run separately, and you can combine multiple datasets later at the sequence table step. Be sure to include controls or low quality samples for learning error. When using smaller datasets, DADA2 recommended nbases=1e6 reads, but for the big data workflow, it recommended nbases=2e6. Multithread = TRUE if you are using a mac that can handle it.
 errJV31F <- learnErrors(derepJV31F, nbases = 1e6, multithread=FALSE, randomize=TRUE)
 errJV31R <- learnErrors(derepJV31R, nbases = 1e6, multithread=FALSE, randomize=TRUE)


# save error profiles to output files
saveRDS(errJV31F, 'error_profile_JV31F_lobster.RDS')
saveRDS(errJV31R, 'error_profile_JV31R_lobster.RDS')

# OPTIONAL: look at error plots to make sure correlation is good, optional
 plotErrors(errJV31F, nominalQ=TRUE)
 plotErrors(errJV31R, nominalQ=TRUE)


 
## DADA2 sample inference, a.k.a. pick the sequence variants ----------------------
##  Do not pool different sequencing runs as they have their own inherent error rates. Can set the error rate as a matrix that has already been learned, or learn now as selfConsist=TRUE

 # Sample inference and merger of PAIRED-END reads.  dereplicates, picks SVs, and merges pairs all in one loop
 
 # this is a loop, run the whole chunk
JV31mergers <- vector("list", length(sample.namesJV31F))
names(JV31mergers) <- sample.namesJV31F
for(sam in sample.namesJV31F) {
  cat("Processing:", sam, "\n")
  JV31derepF <- derepFastq(JV31filtsF[[sam]])
  JV31dadaFs <- dada(JV31derepF, err=errJV31F, multithread=TRUE) # sample inference on the forwards. if using 454 pyrosequencing data, add HOMOPOLYMER_GAP_PENALTY=-1, BAND_SIZE=32
  JV31derepR <- derepFastq(JV31filtsR[[sam]])
  JV31dadaRs <- dada(JV31derepR, err=errJV31R, multithread=TRUE) #sample inference on the reverse. if using 454 pyrosequencing data, add HOMOPOLYMER_GAP_PENALTY=-1, BAND_SIZE=32
  JV31merger <- mergePairs(JV31dadaFs, JV31derepF, JV31dadaRs, JV31derepR) # merge the forwards and reverses
  JV31mergers[[sam]] <- JV31merger
}
# end of loop


## DADA2 Make a sequence table and remove chimeras ----------------------------
# Seqtab is comparable to OTU table. Chimeras are artificially created sequences caused during amplification steps that go wrong.

# Construct sequence table, using != will remove rows of unwanted data
JV31seqtab <- makeSequenceTable(JV31mergers) #if you merged F/R

dim(JV31seqtab) 

	saveRDS(JV31seqtab, '~/JV31/JV31seqtab_lobster.rds') #CHANGE ME
	
	JV31seqtab.nochim <- removeBimeraDenovo(JV31seqtab, method="consensus", multithread=TRUE, verbose=TRUE) 
# record # 2972 chimeras out of # 16,465 of seqs

dim(JV31seqtab.nochim) 

	saveRDS(JV31seqtab.nochim, '~/JV31/JV31seqtab.nochim_lobster.rds') #CHANGE ME
	

	
#### ----- repeat for JV35 ------
	## DADA2 learn error rates  ----------------------------
	# DADA2 learns error rates for each sequencing run separately, because some error is inherent to the run.  Uses a random subset of bases from the data, DADA2 pipeline recommends learning the error rates prior to dereplication (combining identical sequences into one).
	
	set.seed(1234) # set the seed for using random number generators.  setting a seed means it can be reproduced because you start in the same place.
	
	JV35_filt_F <- '~/JV35/JV35_forward_filtered' #CHANGE ME
	JV35_filt_R <- '~/JV35/JV35_reverse_filtered' #CHANGE ME
	
	
	## set the file names for Forwards
	JV35fnsF <- list.files(JV35_filt_F, full.names = TRUE)
	JV35filtsF <- JV35fnsF[grepl('.gz$', JV35fnsF)] # CHANGE if different file extensions
	sample.namesJV35F <- sapply(strsplit(basename(JV35filtsF), '_'), `[`, 1) # Assumes filename = samplename_XXX.fastq
	names(JV35filtsF) <- sample.namesJV35F 
	
	## set the file names for Reverses
	JV35fnsR <- list.files(JV35_filt_R, full.names = TRUE)
	JV35filtsR <- JV35fnsR[grepl('.gz$', JV35fnsR)] # CHANGE if different file extensions
	sample.namesJV35R <- sapply(strsplit(basename(JV35filtsR), '_'), `[`, 1) # Assumes filename = samplename_XXX.fastq
	names(JV35filtsR) <- sample.namesJV35R
	
	
	# dereplicate, which compresses identical sequences into one to make computing faster.  DADA2 will retain the quality info for all seqs that were compressed. 
	derepJV35F <- derepFastq(JV35filtsF, verbose=TRUE)
	derepJV35R <- derepFastq(JV35filtsR, verbose=TRUE)
	
	
	## learn the error rates from your sequencing run. Some error is inherant to sequencing run, so this should be done on each run separately, and you can combine multiple datasets later at the sequence table step. Be sure to include controls or low quality samples for learning error. When using smaller datasets, DADA2 recommended nbases=1e6 reads, but for the big data workflow, it recommended nbases=2e6. Multithread = TRUE if you are using a mac that can handle it.
	errJV35F <- learnErrors(derepJV35F, nbases = 1e6, multithread=FALSE, randomize=TRUE)
	errJV35R <- learnErrors(derepJV35R, nbases = 1e6, multithread=FALSE, randomize=TRUE)
	
	
	# save error profiles to output files
	saveRDS(errJV35F, 'error_profile_JV35F_lobster.RDS')
	saveRDS(errJV35R, 'error_profile_JV35R_lobster.RDS')
	
	# OPTIONAL: look at error plots to make sure correlation is good, optional
	plotErrors(errJV35F, nominalQ=TRUE)
	plotErrors(errJV35R, nominalQ=TRUE)
	
	
	
	## DADA2 sample inference, a.k.a. pick the sequence variants ----------------------
	##  Do not pool different sequencing runs as they have their own inherent error rates. Can set the error rate as a matrix that has already been learned, or learn now as selfConsist=TRUE
	
	# Sample inference and merger of PAIRED-END reads.  dereplicates, picks SVs, and merges pairs all in one loop
	
	# this is a loop, run the whole chunk
	JV35mergers <- vector("list", length(sample.namesJV35F))
	names(JV35mergers) <- sample.namesJV35F
	for(sam in sample.namesJV35F) {
	  cat("Processing:", sam, "\n")
	  JV35derepF <- derepFastq(JV35filtsF[[sam]])
	  JV35dadaFs <- dada(JV35derepF, err=errJV35F, multithread=TRUE) # sample inference on the forwards. if using 454 pyrosequencing data, add HOMOPOLYMER_GAP_PENALTY=-1, BAND_SIZE=32
	  JV35derepR <- derepFastq(JV35filtsR[[sam]])
	  JV35dadaRs <- dada(JV35derepR, err=errJV35R, multithread=TRUE) #sample inference on the reverse. if using 454 pyrosequencing data, add HOMOPOLYMER_GAP_PENALTY=-1, BAND_SIZE=32
	  JV35merger <- mergePairs(JV35dadaFs, JV35derepF, JV35dadaRs, JV35derepR) # merge the forwards and reverses
	  JV35mergers[[sam]] <- JV35merger
	}
	# end of loop
	
	
	## DADA2 Make a sequence table and remove chimeras ----------------------------
	# Seqtab is comparable to OTU table. Chimeras are artificially created sequences caused during amplification steps that go wrong.
	
	# Construct sequence table, using != will remove rows of unwanted data
	JV35seqtab <- makeSequenceTable(JV35mergers) #if you merged F/R
	
	dim(JV35seqtab) 

	saveRDS(JV35seqtab, '~/JV35/JV35seqtab_lobster.rds') #CHANGE ME
	
	JV35seqtab.nochim <- removeBimeraDenovo(JV35seqtab, method="consensus", multithread=TRUE, verbose=TRUE) 

	dim(JV35seqtab.nochim) 

	saveRDS(JV35seqtab.nochim, '~/JV35/JV35seqtab.nochim_lobster.rds') #CHANGE ME
	


#### ----- repeat for JV39 ------
	## DADA2 learn error rates  ----------------------------
	# DADA2 learns error rates for each sequencing run separately, because some error is inherent to the run.  Uses a random subset of bases from the data, DADA2 pipeline recommends learning the error rates prior to dereplication (combining identical sequences into one).
	
	set.seed(1234) # set the seed for using random number generators.  setting a seed means it can be reproduced because you start in the same place.
	
	JV39_filt_F <- '~/JV39/JV39_forward_filtered' #CHANGE ME
	JV39_filt_R <- '~/JV39/JV39_reverse_filtered' #CHANGE ME
	
	
	## set the file names for Forwards
	JV39fnsF <- list.files(JV39_filt_F, full.names = TRUE)
	JV39filtsF <- JV39fnsF[grepl('.gz$', JV39fnsF)] # CHANGE if different file extensions
	sample.namesJV39F <- sapply(strsplit(basename(JV39filtsF), '_'), `[`, 1) # Assumes filename = samplename_XXX.fastq
	names(JV39filtsF) <- sample.namesJV39F 
	
	## set the file names for Reverses
	JV39fnsR <- list.files(JV39_filt_R, full.names = TRUE)
	JV39filtsR <- JV39fnsR[grepl('.gz$', JV39fnsR)] # CHANGE if different file extensions
	sample.namesJV39R <- sapply(strsplit(basename(JV39filtsR), '_'), `[`, 1) # Assumes filename = samplename_XXX.fastq
	names(JV39filtsR) <- sample.namesJV39R
	
	
	# dereplicate, which compresses identical sequences into one to make computing faster.  DADA2 will retain the quality info for all seqs that were compressed. 
	derepJV39F <- derepFastq(JV39filtsF, verbose=TRUE)
	derepJV39R <- derepFastq(JV39filtsR, verbose=TRUE)
	
	
	## learn the error rates from your sequencing run. Some error is inherant to sequencing run, so this should be done on each run separately, and you can combine multiple datasets later at the sequence table step. Be sure to include controls or low quality samples for learning error. When using smaller datasets, DADA2 recommended nbases=1e6 reads, but for the big data workflow, it recommended nbases=2e6. Multithread = TRUE if you are using a mac that can handle it.
	errJV39F <- learnErrors(derepJV39F, nbases = 1e6, multithread=FALSE, randomize=TRUE)
	errJV39R <- learnErrors(derepJV39R, nbases = 1e6, multithread=FALSE, randomize=TRUE)
	
	
	# save error profiles to output files
	saveRDS(errJV39F, 'error_profile_JV39F_lobster.RDS')
	saveRDS(errJV39R, 'error_profile_JV39R_lobster.RDS')
	
	# OPTIONAL: look at error plots to make sure correlation is good, optional
	plotErrors(errJV39F, nominalQ=TRUE)
	plotErrors(errJV39R, nominalQ=TRUE)
	
	
	
	## DADA2 sample inference, a.k.a. pick the sequence variants ----------------------
	##  Do not pool different sequencing runs as they have their own inherent error rates. Can set the error rate as a matrix that has already been learned, or learn now as selfConsist=TRUE
	
	# Sample inference and merger of PAIRED-END reads.  dereplicates, picks SVs, and merges pairs all in one loop
	
	# this is a loop, run the whole chunk
	JV39mergers <- vector("list", length(sample.namesJV39F))
	names(JV39mergers) <- sample.namesJV39F
	for(sam in sample.namesJV39F) {
	  cat("Processing:", sam, "\n")
	  JV39derepF <- derepFastq(JV39filtsF[[sam]])
	  JV39dadaFs <- dada(JV39derepF, err=errJV39F, multithread=TRUE) # sample inference on the forwards. if using 454 pyrosequencing data, add HOMOPOLYMER_GAP_PENALTY=-1, BAND_SIZE=32
	  JV39derepR <- derepFastq(JV39filtsR[[sam]])
	  JV39dadaRs <- dada(JV39derepR, err=errJV39R, multithread=TRUE) #sample inference on the reverse. if using 454 pyrosequencing data, add HOMOPOLYMER_GAP_PENALTY=-1, BAND_SIZE=32
	  JV39merger <- mergePairs(JV39dadaFs, JV39derepF, JV39dadaRs, JV39derepR) # merge the forwards and reverses
	  JV39mergers[[sam]] <- JV39merger
	}
	# end of loop
	
	
	## DADA2 Make a sequence table and remove chimeras ----------------------------
	# Seqtab is comparable to OTU table. Chimeras are artificially created sequences caused during amplification steps that go wrong.
	
	# Construct sequence table, using != will remove rows of unwanted data
	JV39seqtab <- makeSequenceTable(JV39mergers) #if you merged F/R
	
	dim(JV39seqtab) 

	saveRDS(JV39seqtab, '~/JV39/JV39seqtab_lobster.rds') #CHANGE ME
	
	JV39seqtab.nochim <- removeBimeraDenovo(JV39seqtab, method="consensus", multithread=TRUE, verbose=TRUE) 

	dim(JV39seqtab.nochim) 

	saveRDS(JV39seqtab.nochim, '~/JV39/JV39seqtab.nochim_lobster.rds') #CHANGE ME
	

	

#### ----- repeat for JV41 ------
	## DADA2 learn error rates - ----------------------------
	# DADA2 learns error rates for each sequencing run separately, because some error is inherent to the run.  Uses a random subset of bases from the data, DADA2 pipeline recommends learning the error rates prior to dereplication (combining identical sequences into one).
	
	set.seed(1234) # set the seed for using random number generators.  setting a seed means it can be reproduced because you start in the same place.
	
	JV41_filt_F <- '~/JV41/JV41_forward_filtered' #CHANGE ME
	JV41_filt_R <- '~/JV41/JV41_reverse_filtered' #CHANGE ME
	
	
	## set the file names for Forwards
	JV41fnsF <- list.files(JV41_filt_F, full.names = TRUE)
	JV41filtsF <- JV41fnsF[grepl('.gz$', JV41fnsF)] # CHANGE if different file extensions
	sample.namesJV41F <- sapply(strsplit(basename(JV41filtsF), '_'), `[`, 1) # Assumes filename = samplename_XXX.fastq
	names(JV41filtsF) <- sample.namesJV41F 
	
	## set the file names for Reverses
	JV41fnsR <- list.files(JV41_filt_R, full.names = TRUE)
	JV41filtsR <- JV41fnsR[grepl('.gz$', JV41fnsR)] # CHANGE if different file extensions
	sample.namesJV41R <- sapply(strsplit(basename(JV41filtsR), '_'), `[`, 1) # Assumes filename = samplename_XXX.fastq
	names(JV41filtsR) <- sample.namesJV41R
	
	
	# dereplicate, which compresses identical sequences into one to make computing faster.  DADA2 will retain the quality info for all seqs that were compressed. 
	derepJV41F <- derepFastq(JV41filtsF, verbose=TRUE)
	derepJV41R <- derepFastq(JV41filtsR, verbose=TRUE)
	
	
	## learn the error rates from your sequencing run. Some error is inherant to sequencing run, so this should be done on each run separately, and you can combine multiple datasets later at the sequence table step. Be sure to include controls or low quality samples for learning error. When using smaller datasets, DADA2 recommended nbases=1e6 reads, but for the big data workflow, it recommended nbases=2e6. Multithread = TRUE if you are using a mac that can handle it.
	errJV41F <- learnErrors(derepJV41F, nbases = 1e6, multithread=FALSE, randomize=TRUE)
	errJV41R <- learnErrors(derepJV41R, nbases = 1e6, multithread=FALSE, randomize=TRUE)
	
	
	# save error profiles to output files
	saveRDS(errJV41F, 'error_profile_JV41F_lobster.RDS')
	saveRDS(errJV41R, 'error_profile_JV41R_lobster.RDS')
	
	# OPTIONAL: look at error plots to make sure correlation is good, optional
	plotErrors(errJV41F, nominalQ=TRUE)
	plotErrors(errJV41R, nominalQ=TRUE)
	
	
	
	## DADA2 sample inference, a.k.a. pick the sequence variants ----------------------
	##  Do not pool different sequencing runs as they have their own inherent error rates. Can set the error rate as a matrix that has already been learned, or learn now as selfConsist=TRUE
	
	# Sample inference and merger of PAIRED-END reads.  dereplicates, picks SVs, and merges pairs all in one loop
	
	# this is a loop, run the whole chunk
	JV41mergers <- vector("list", length(sample.namesJV41F))
	names(JV41mergers) <- sample.namesJV41F
	for(sam in sample.namesJV41F) {
	  cat("Processing:", sam, "\n")
	  JV41derepF <- derepFastq(JV41filtsF[[sam]])
	  JV41dadaFs <- dada(JV41derepF, err=errJV41F, multithread=TRUE) # sample inference on the forwards. if using 454 pyrosequencing data, add HOMOPOLYMER_GAP_PENALTY=-1, BAND_SIZE=32
	  JV41derepR <- derepFastq(JV41filtsR[[sam]])
	  JV41dadaRs <- dada(JV41derepR, err=errJV41R, multithread=TRUE) #sample inference on the reverse. if using 454 pyrosequencing data, add HOMOPOLYMER_GAP_PENALTY=-1, BAND_SIZE=32
	  JV41merger <- mergePairs(JV41dadaFs, JV41derepF, JV41dadaRs, JV41derepR) # merge the forwards and reverses
	  JV41mergers[[sam]] <- JV41merger
	}
	# end of loop
	
	
## DADA2 Make a sequence table and remove chimeras ----------------------------
	# Seqtab is comparable to OTU table. Chimeras are artificially created sequences caused during amplification steps that go wrong.
	
	# Construct sequence table, using != will remove rows of unwanted data
	JV41seqtab <- makeSequenceTable(JV41mergers) #if you merged F/R
	
	dim(JV41seqtab) 

	saveRDS(JV41seqtab, '~/JV41/JV41seqtab_lobster.rds') #CHANGE ME
	
	JV41seqtab.nochim <- removeBimeraDenovo(JV41seqtab, method="consensus", multithread=TRUE, verbose=TRUE) 

	dim(JV41seqtab.nochim) 

	saveRDS(JV41seqtab.nochim, '~/JV41/JV41seqtab.nochim_lobster.rds') #CHANGE ME
	
		

## Workflow verification steps --------------------------

# Track reads through the analysis here to see how many were lost at each QA step, in case you were too stringent. Only need to count F reads, even if you did paired F/R
getN <- function(x) sum(getUniques(x)) #set function to count the unique reads
	
# load the filtered output file as needed
filtoutput <- readRDS("filtered_output_JV41.rds")
	
	# load the metadata files so you can match factors to seq data
meta <- read.csv('~/JV41/JV41_metadata.csv', header = TRUE, row.names = 1) #row.names specifies which column to call the samples by, doesn't have to be the first column

JV41seqtab.nochim <- readRDS('~/JV41/JV41seqtab.nochim_lobster.rds') #CHANGE ME

track <- cbind(filtoutput, rowSums(JV41seqtab.nochim), meta$Sample_type) #bind columns from filtered output, # of seqs/sample from the no.chim seq table, and the treatment factor
colnames(track) <- c("reads.in","filtered", "nonchimeras", "Sample_type")
rownames(track) <- rownames(meta)

head(track)

# save the tracking data as an R file
saveRDS(track, 'tracked_seqs_JV41.RDS')


require(tidyr)
library(ggplot2)

plotData <- as.data.frame(track) %>% gather(type,totals, reads.in, filtered,nonchimeras)

#relabel the sample type variables
plotData$Sample_type <- factor(plotData$Sample_type, labels=c("negative_control", "tank_pH","tank_temperature")) #CHANGE ME

#set the count type order
plotData$type <- factor(plotData$type, levels=c("reads.in", "filtered","nonchimeras")) #CHANGE ME

ggplot(plotData,aes(x=type,y=as.numeric(totals))) + geom_point(aes(color=Sample_type)) + ylab("Sequences") + xlab("QA stage") #CHANGE ME


# randomly select a negative control to check
unqs.NC <- seqtab.nochim["NegControlbatch1",] #CHANGE ME, update with sample name of a negative control
unqs.NC <- sort(unqs.NC[unqs.NC>0], decreasing=TRUE) # Drop ASVs absent in the sample
cat("DADA2 inferred", length(unqs.NC), "sample sequences present in the selected sample.\n")
	# how many reads are inferred in the negative control?

plot(unqs.NC, ylab="Number of seqs/SV, Neg Control", xlab="SVs", main="Number of sequences/SV in Negative Control") #CHANGE ME

# OPTIONAL: add taxonomy and see what's in it
taxa.NC <- assignTaxonomy(unqs.NC, '~silva_nr_v138_train_set.fa.gz', minBoot = 75)
	write.csv(taxa.NC, '~taxa_silva_NC_EXAMPLE.csv') 

	
# randomly select a positive control to check
unqs.PC <- seqtab.nochim["sample_name_of_positive_control",] #CHANGE ME, update with sample name of a positive contol
unqs.PC <- sort(unqs.PC[unqs.PC>0], decreasing=TRUE) # Drop ASVs absent in the sample
cat("DADA2 inferred", length(unqs.PC), "sample sequences present in the selected sample.\n")
	# how many reads are inferred in the negative control?
	
plot(unqs.PC, ylab="Number of seqs/SV, Neg Control", xlab="SVs", main="Number of sequences/SV in Positive Control") #CHANGE ME
	
# OPTIONAL: add taxonomy and see what's in it
taxa.PC <- assignTaxonomy(unqs.mock, '~silva_nr_v138_train_set.fa.gz', minBoot = 75)
	write.csv(taxa.PC, '~taxa_silva_PC_EXAMPLE.csv') 
	

## DADA2 picking taxonomy ---------------------------------
## look at the number of SVs/samples they are found in
nsam <- colSums(seqtab.nochim>0)
plot(nsam, xlab="Number of SVs", ylab="Number of Samples the SVs are found in", main="Seqtab.all.nochim and how many samples each SV is found in")
# seqtab.all.nochim3 <- seqtab.all.nochim[,nsam>=3] ## only about 8,000 SVs are found in more than 3 samples
# nsam <- colSums(seqtab.all.nochim3>0)
# plot(nsam, xlab="Number of SVs", ylab="Number of Samples the SVs are found in")

# load datasets
seqtab.nochimJV31 <- readRDS('JV31seqtab.nochim_lobster.rds')

## assign taxonomy. this may be memory intensive.
all.taxa <- assignTaxonomy(seqtab.nochimJV31, '~/silva_nr_v138_train_set.fa.gz', minBoot = 75) #CHANGE file path
  saveRDS(all.taxa, 'taxa_silva138_JV31.rds') #CHANGE ME
  write.csv(all.taxa, 'taxa_silva138_JV31.csv') 

# try adding species designation to the table. this may be memory intensive.
all.taxa.sp <- addSpecies(all.taxa, 'silva_species_assignment_v138.fa.gz', allowMultiple = FALSE, verbose = FALSE) #CHANGE file path
  saveRDS(all.taxa.sp, 'taxa_species_silva138_JV31.rds') #CHANGE ME
  

  # load datasets
  seqtab.nochimJV39 <- readRDS('JV39seqtab.nochim_lobster.rds')
  all.taxa <- read.csv('taxa_silva_JV39.csv', header = T, row.names=1)
  
  
  # try adding species designation to the table. this may be memory intensive.
  all.taxa.sp <- addSpecies(all.taxa, 'silva_species_assignment_v138.fa.gz', allowMultiple = FALSE, verbose = FALSE) #CHANGE file path
  saveRDS(all.taxa.sp, 'taxa_species_silva138_JV31.rds') #CHANGE ME
  
  
  
  
  
## First look and QA with neg con in Phyloseq ------------------------------
library(phyloseq)
  
# load seqtab as needed
seqtab.nochim <- readRDS('~/JV31/JV31seqtab.nochim_lobster.rds') #CHANGE ME

## load taxa table as needed
all.taxa.sp <- read.csv('~/JV31/taxa_species_silva138_JV31.csv', header=T, row.names=1) #CHANGE ME

# load metadata table as needed
meta <- read.csv("~/JV31/JV31_metadata.csv", header=T, row.names=1) #CHANGE ME

## create a phyloseq object with all samples (subset out later)
Lobster_ps <- phyloseq(otu_table(seqtab.nochim, taxa_are_rows=FALSE), 
               sample_data(meta), 
               tax_table(as.matrix(all.taxa.sp)))

Lobster_ps
# how many samples made it this far? CHANGE ME


# If you have sequenced controls, it's a good idea to look at your data in comparison to them.

# Alpha diversity peek 
 plot_richness(Lobster_ps, x="Sample_type", #define the x-axis
               measures=c("Observed","Chao1", "Shannon"), 
               color="Covercrop_Treatment") + #color the dots by a factor
   theme_bw() #a ggplot theme to make the graph look nice
 
## Note: hopefully, diversity/richness is lower in the Negative controls than real samples

 
# Plot the taxa sums to see how populated each taxa is (do you have many rare taxa?
	plot(sort(taxa_sums(Lobster_ps), TRUE), 
	     type="h", 
	     ylim=c(0, 20)) #limit the y-axis to better see the long tail of rare taxa

	

# Create a simple ordination to look for clustering by extraction batch or confounding variables in your metadata 

### PCOA of all samples per run	
 EX.ord <- ordinate(Lobster_ps, #calculate similarities
                    method ="PCoA", #ordination type
                    "jaccard", binary = TRUE) #similarity type. Jaccard is binary, Bray can be binary (unweighted) or not (weighted)
 
 
 plot_ordination(Lobster_ps, EX.ord, type="samples", color="Sample_type")
	# Initial clustering by batch or confounding variables implies contamination issues (see next section)
 # Horse-shoe patterns indicate underlying patterns to the data


## Removing contamination using negative controls-----------------------------------
 ### This section can only be accomplished if you have sequenced negative controls and/or DNA quantification data from when the pool was created.  If you have these, CHOOSE ONE method to follow.  If you don't, skip this section and consider including sequencing controls in your next run.
 
# Dr. Ishaq's method: https://github.com/SueIshaq/Examples-DADA2-Phyloseq
# decontam in R: https://benjjneb.github.io/decontam/vignettes/decontam_intro.html
 
# Dr. Ishaq's method creates vectors out of the SV table data for negative controls, and subtracts those SVs from the sample data.  Depending on the type of negative control, these are removed from the whole data set or from subsets of batches. Remove PCR and sampling materials negative control SVs fully from all samples, and remove extraction kit SVs fully from each dna_extraction_batch, respectively.
 
 # For the lobster data, only 4 negative controls were included in 2 runs.  Since they were from the same experiments it might still help by removing biological contamination sourced from the reagents. Combine all 4 lobster sequencing runs together and use that as the phyloseq object here.
 
 ### PCOA of all samples per run before cleaning
 EX.ord <- ordinate(Lobster_ps, #calculate similarities
                            method ="PCoA", #ordination type
                            "jaccard", binary = TRUE) #similarity type. Jaccard is binary, Bray can be binary (unweighted) or not (weighted)
 
 plot_ordination(Lobster_ps, EX.ord, type="samples", color="sequencing run", title="All runs before cleaning out negative controls")
 
 

# Subset the controls and prune to be only those taxa	
	Lobster_ps_controls = subset_samples(Lobster_ps, Temp_geography == "NA") # number of samples CHANGE ME
	Lobster_ps_controls <- prune_taxa(taxa_sums(Lobster_ps_controls) > 0, Lobster_ps_controls)

# Make the taxa names into a vector so you can remove them. 
	control_vec <- as.vector(taxa_names(Lobster_ps_controls))
	vec <- as.vector(taxa_names(Lobster_ps))
	keep <- setdiff(vec, control_vec)

# Use the keep vector for the prune taxa argument, because it wants the argument to be true (matching).
	Lobster_ps_NCclean <- prune_taxa(keep, Lobster_ps)
	
# Then remove the samples which are now empty, namely the NegCon_PCR and NegCon_swab.
	Lobster_ps_NCclean <- prune_samples(sample_sums(Lobster_ps_NCclean) > 0, Lobster_ps_NCclean)
	
	# number of samples and SVs remaining CHANGE ME

# Did it work? Check your ordination again
	### PCOA of all samples per run	
	Lobster_cleaner.ord <- ordinate(Lobster_ps_NCclean, #calculate similarities
	                   method ="PCoA", #ordination type
	                   "jaccard", binary = TRUE) #similarity type. Jaccard is binary, Bray can be binary (unweighted) or not (weighted)
	
	plot_ordination(Lobster_ps_NCclean, Lobster_cleaner.ord, type="samples", color="sequencing run", title="All runs after cleaning out negative controls")
	
	
## Clean out unwanted taxa ------------------------------------
# Regardless of whether you included sequenced negative controls, you can remove taxa which are of no interest to you

# clean out chloroplast and mitochondria. can also elect to remove other contaminating domains or kingdoms as needed
  require(dplyr)
		Lobster_ps_NC_batch_taxa_clean <- Lobster_ps_NC_batch_clean %>% subset_taxa(Kingdom == "Bacteria" & Family != "Mitochondria" & Class != "Chloroplast") # or use Lobster_ps if you had no negative controls

## clean out taxa/SV columns that are no longer present
  Lobster_ps_NC_batch_taxa_clean <- prune_taxa(taxa_sums(Lobster_ps_NC_batch_taxa_clean) > 0, Lobster_ps_NC_batch_taxa_clean)
  
  Lobster_ps_clean <- prune_samples(sample_sums(Lobster_ps_NC_batch_taxa_clean) > 0, Lobster_ps_NC_batch_taxa_clean)
  
  # 183 samples of samples and 8106 SVs left 


# Save your clean phyloseq object
saveRDS(Lobster_ps_clean, 'Lobster_ps_clean_phyloseq_object.RDS') #CHANGE ME

# Reload as needed
Lobster_ps_clean <-readRDS('Lobster_ps_clean_phyloseq_object.RDS') 


## Rarefaction --------------------------------------------------
# To compare sequence data accurately, it is often necessary to rarefy/normalize SVs to even depth across all samples
  # Rarefication reportedly works best (compared to logUQ, CSS, DESeqVS, edgeR-TMM): https://microbiomejournal.biomedcentral.com/articles/10.1186/s40168-017-0237-y
  # Rarefication also not subject to library effect sizes

# take a look at rowsums, or total sequences per sample
rowSums(otu_table(Lobster_ps_clean))

Lobster_ps_clean.rar <- rarefy_even_depth(Lobster_ps_NCclean, 
                                     sample.size=4153, #SVs/sample you want. 5-10k is a good amount, more is better
                                     replace=FALSE, #sampling with or without replacement
                                     trimOTUs=TRUE, #remove OTUs left empty
                                     rngseed=711, 
                                     verbose=TRUE)
#4 samples were removed, and 1343 ASVs removed. 179 samples and 6763 taxa left

# Save your cleaned, rarefied phyloseq object  
saveRDS(Lobster_ps_clean.rar, 'Lobster_ps_clean_rarefied_phyloseq_object.RDS')


  sample_data(Lobster_ps_clean.rar)$Timepoint = factor(sample_data(Lobster_ps_clean.rar)$Timepoint, levels = c("Control", "Baseline", "4_months", "10_months"), labels = c("Control", "Baseline, Summer", "4 months, Winter", "10 months, Summer"))
  
## make a plot to look at lobster shell disease and mortality with all temperature samples
  ggplot(data=subset(sample_data(Lobster_ps_clean.rar), !is.na(Lobster_ID)), aes(x=Timepoint, y=Shell_Disease_Index, color=as.factor(Lobster_ID), group=(as.factor(Lobster_ID))))+
    theme_bw()+
    facet_grid(Tank~.) +
    geom_line() +
    geom_point(aes(shape = as.factor(Tank))) +
    ylab("Shell Disease Index") + labs(shape = "Tank", color= "Lobster ID")
  
  
  require(lubridate)
  ##### as.Date to import date into a usable format for continuous variable
    sample_data(Lobster_ps_clean.rar)$Sample_date = as.Date(sample_data(Lobster_ps_clean.rar)$Sample_date, format = "%d-%b-%y")
    sample_data(Lobster_ps_clean.rar)$Molt = as.Date(sample_data(Lobster_ps_clean.rar)$Molt, format = "%m/%d/%Y")
    sample_data(Lobster_ps_clean.rar)$Died = as.Date(sample_data(Lobster_ps_clean.rar)$Died, format = "%m/%d/%Y")
    
    
  ## make a plot to look at lobster shell disease and mortality with all temperature samples
  ggplot(data=subset(sample_data(Lobster_ps_clean.rar), !is.na(Lobster_ID)), aes(x=Sample_date, y=Shell_Disease_Index, color=as.factor(Lobster_ID), group=(as.factor(Lobster_ID))))+
    theme_bw()+
    scale_x_date(date_labels = "%y-%m-%d") +
    facet_grid(Tank~.) +
    geom_line(show.legend = FALSE) +
    geom_point(aes(shape = as.factor(Tank)), size =4) +
    geom_vline(aes(xintercept=Molt, color=as.factor(Lobster_ID)), size=1, show.legend=FALSE) +
    geom_vline(aes(xintercept=Died, color=as.factor(Lobster_ID)), linetype = "longdash", size=2, show.legend=FALSE) +
    ylab("Shell Disease Index") + xlab("Sample Date") + labs(shape = "Tank", color= "Lobster ID")
  
# view individual lobster plots by SDI with molt and death dates
  ggplot(data=subset(sample_data(Lobster_ps_clean.rar), !is.na(Lobster_ID)), aes(x=Sample_date, y=Shell_Disease_Index, color=as.factor(Lobster_ID), fill=as.factor(Lobster_ID), group=(as.factor(Lobster_ID))))+
    theme_bw()+
    scale_x_date(date_labels = "%y-%m-%d") +
    facet_wrap(Tank~Lobster_ID) +
    geom_line() +
    geom_point(aes(shape = as.factor(Tank)), size =4) +
    geom_vline(aes(xintercept=Molt), color="red", size=1.5) +
    geom_vline(aes(xintercept=Died), color="black", size =1.5) +
    ylab("Shell Disease Index") + xlab("Sample Date") +theme(legend.position = "none")
  

## Pick up the statistical analysis here -------------------------------------
# Once you are satisfied with your QA, you can start from here with your phyloseq object as needed

library(phyloseq)

#non-rarefied version of the temp experiment
Lobster_ps_clean <-readRDS("Lobster_clean_temponly_phyloseq_object.RDS")

# Rarefied version of temp experiment
Lobster_ps_clean.rar <-readRDS("Lobster_clean_rarefied_temponly_phyloseq_object.RDS")


## ------ Investigating molt and death date ----------

# view individual lobster plots by SDI with molt and death dates
ggplot(data=subset(sample_data(Lobster_ps_clean.rar), !is.na(Lobster_ID)), aes(x=Sample_date, y=Shell_Disease_Index, color=as.factor(Lobster_ID), fill=as.factor(Lobster_ID), group=(as.factor(Lobster_ID))))+
  theme_bw()+
  scale_x_date(date_labels = "%y-%m-%d") +
  facet_wrap(Tank~Lobster_ID) +
  geom_line() +
  geom_point(aes(shape = as.factor(Tank)), size =4) +
  geom_vline(aes(xintercept=Molt), color="red", size=1.5) +
  geom_vline(aes(xintercept=Died), color="black", size =1.5) +
  ylab("Shell Disease Index") + xlab("Sample Date") +theme(legend.position = "none")

require(lubridate)
##### as.Date to import date into a usable format for continuous variable
sample_data(Lobster_ps_clean.rar)$Sample_date = as.Date(sample_data(Lobster_ps_clean.rar)$Sample_date, format = "%d-%b-%y")
sample_data(Lobster_ps_clean.rar)$Molt = as.Date(sample_data(Lobster_ps_clean.rar)$Molt, format = "%m/%d/%Y")
sample_data(Lobster_ps_clean.rar)$Died = as.Date(sample_data(Lobster_ps_clean.rar)$Died, format = "%m/%d/%Y")


# Lobster_temp_richness_molt_died_LobsterID
plot_richness(Lobster_ps_clean.rar, x='Sample_date', measures = "Observed", color='Tank_temp_C') + facet_wrap(Lobster_ID~.) + theme_bw() + scale_fill_viridis(discrete=FALSE) + scale_color_viridis(discrete=FALSE) + geom_vline(aes(xintercept=Molt), size=1 ) + geom_vline(aes(xintercept=Died), linetype = "longdash", size=2) + xlab("Sample Date") + labs(color = "Tank temperature, C")

#Summary: richness does not appear to be associated with mortality, and since so many lobsters molted right after being put in tanks it's hard to say if the reduction in richness is associated with that molt or not. For the 2 lobsters that did molt between the 2nd and 3rd sampling, richness appeared unchanged

# culturing data correlation with richness
plot_richness(Lobster_ps_clean.rar, x='BAwNaCL_Per1cm2', measures = "Observed", color='Tank_temp_C') + facet_grid(Temp_geography~.) + scale_x_continuous(trans='log10') + theme_bw() + scale_fill_viridis(discrete=FALSE) + scale_color_viridis(discrete=FALSE) + geom_smooth() + xlab("CFU per cm2 on BA + NaCl media") + labs(color = "Tank temperature, C")

plot_richness(Lobster_ps_clean.rar, x='Marine_Per1cm2', measures = "Observed", color='Tank_temp_C') + facet_grid(Temp_geography~.) + scale_x_continuous(trans='log10') + theme_bw() + scale_fill_viridis(discrete=FALSE) + scale_color_viridis(discrete=FALSE) + geom_smooth() + xlab("CFU per cm2 on Marine media") + labs(color = "Tank temperature, C")
#Summary: they all appear to have a unimodal distribution

plot_richness(Lobster_ps_clean.rar, x='Sample_date', measures = "Observed", color='Tank_temp_C') + geom_point(aes(size =BAwNaCL_Per1cm2, shape = as.factor(Shell_Disease_Index))) + facet_grid(Temp_geography*Tank~.) + theme_bw() + scale_fill_viridis(discrete=FALSE) + scale_color_viridis(discrete=FALSE) + geom_vline(aes(xintercept=Molt), size=1 ) + geom_vline(aes(xintercept=Died), linetype = "longdash", size=2) + xlab("Sample Date") + labs(color = "Tank temperature, C") + scale_size(name   = "BA + NaCl Media CFU per cm2", breaks = fivenum(sample_data(Lobster_ps_clean.rar)$BAwNaCL_Per1cm2),labels = fivenum(sample_data(Lobster_ps_clean.rar)$BAwNaCL_Per1cm2))
# Summary: this does not seem to show a pattern, gets low and stays low

tank_lab <- c("Healthy", "ESD", "ESD", "Healthy", "Healthy", "ESD")
names(tank_lab) <- c("1", "2", "3", "4", "5", "6")
sample_data(Lobster_temp.rar)$Tank <- factor(sample_data(Lobster_temp.rar)$Tank, levels = c("1", "2", "4", "3", "5", "6"))


plot_richness(Lobster_temp.rar, x='Sample_date', measures = "Observed", color='Tank_temp_C') + geom_point(aes(size =Marine_Per1cm2, shape = as.factor(Shell_Disease_Index))) + facet_grid(Temp_geography*Tank~., labeller = labeller(Tank = tank_lab)) + theme_bw() + scale_fill_viridis(discrete=FALSE, option="magma",  begin = 0, end =0.75) + scale_color_viridis(discrete=FALSE, option="magma", begin = 0, end =0.75) + geom_vline(aes(xintercept=Molt), size=1 ) + geom_vline(aes(xintercept=Died), linetype = "longdash", size=2) + xlab("Sample Date") + ylab("Bacterial Richness (SVs)") +labs(color = "Tank temperature, C", shape= "Shell Disease Index") + scale_size(name   = "Marine Media CFU per cm2", breaks = fivenum(sample_data(Lobster_temp.rar)$Marine_Per1cm2),labels = fivenum(sample_data(Lobster_temp.rar)$Marine_Per1cm2))
#
# Summary: in tanks with disease, gets high again based on temp

# Summary: when looking at each individual lobster, did not see definable pattern


## What's in tank controls ---------------------------------
Lobster_temp_tank = subset_samples(Lobster_ps_clean.rar, Sample_type == "tank")
Lobster_temp_tank <- prune_samples(sample_sums(Lobster_temp_tank) > 0, Lobster_temp_tank)
Lobster_temp_tank <- prune_taxa(taxa_sums(Lobster_temp_tank) > 0, Lobster_temp_tank)

install.packages("viridis")
library(viridis)
library(ggpubr)

plotA <- plot_richness(Lobster_temp_tank, x="Sample_type", #define the x-axis
              measures=c("Observed", "Shannon"), 
              color="Temp_geography") + #color the dots by a factor
              theme_bw() + #a ggplot theme to make the graph look nice
              scale_fill_viridis(option="magma") +
              labs(color="Temperature geography")
  
plotB <- plot_heatmap(Lobster_temp_tank)


# plot together
ggarrange(plotA, plotB,
          ncol = 1, nrow=2, labels =c("A", "B"), common.legend = FALSE)

## Alpha diversity ---------------------------------
  # ----- Diversity metrics  --------------
# Phyloseq can measure and visualize alpha diversity: https://joey711.github.io/phyloseq/plot_richness-examples.html
# phyloseq doesn't do stats or more complex graphs

#library(phyloseq)

# use phyloseq to measure alpha diversity
Lobster_ps_clean.rar.rich <- estimate_richness(Lobster_ps_clean.rar, measure=c("Observed", "Shannon")) #change to whatever measures you want


#calculate evenness
# measure evenness for each sample
Lobster_ps_clean.rar.even <- Lobster_ps_clean.rar.rich$Shannon/log(Lobster_ps_clean.rar.rich$Observed)

# Coerce to data.frame and add the metadata for these samples
Lobster_ps_clean.rar.sd = as(sample_data(Lobster_ps_clean.rar), "matrix")
Lobster_ps_clean.rar.sd = as.data.frame(Lobster_ps_clean.rar.sd)
Lobster_ps_clean.rar.rich.df <- cbind(Lobster_ps_clean.rar.rich, Lobster_ps_clean.rar.even, Lobster_ps_clean.rar.sd)

#check the distribution of your data, which will change you approach
shapiro.test(Lobster_ps_clean.rar.rich.df$Shannon)
# W = 0.96781, p-value = 0.000376 (non-normal): p-value > 0.05 reveals that the distribution of the data are not significantly different from normal distribution

shapiro.test(Lobster_ps_clean.rar.rich.df$Observed)
# W = 0.99012, p-value = 0.2523 (normal): p-value > 0.05 reveals that the distribution of the data are not significantly different from normal distribution

shapiro.test(Lobster_ps_clean.rar.rich.df$Lobster_ps_clean.rar.even)
# W = 0.92484, p-value = 5.496e-08 (non-normal): p-value > 0.05 reveals that the distribution of the data are not significantly different from normal distribution


install.packages("viridis")
library(viridis)

head(Lobster_ps_clean.rar.rich.df)


Lobster_ps_clean.rar.rich.df$Sample_date <- factor(Lobster_ps_clean.rar.rich.df$Sample_date, levels=c("19-Jul-16", "20-Jul-16", "21-Jul-16", "11-Nov-16", "30-Nov-16", "1-Dec-16", "2-Dec-16", "6-Jun-17", "7-Jun-17", "NA")) #CHANGE ME


  # ----- If your alpha diversity is not normally distributed  -----------------
# If you have a more complex study design, you can use generalized linear mixed effects models

library(lme4)
library(lmerTest)
library(emmeans)

# does not have a normal distribution, so compare means using glmer and poisson distribution (number of events within a time interval): https://towardsdatascience.com/the-poisson-distribution-and-poisson-process-explained-4e2cb17d459

# run each separately to find out if they are significant on their own. Be sure to drop the control tank samples.
summary(lm(Observed ~ as.numeric(Tank_temp_C), data=subset(Lobster_ps_clean.rar.rich.df, Sample_type=="shell")))
# paste output here

summary(lm(Observed ~ Tank_temp_C + (Tank_temp_C/Timepoint), data=subset(Lobster_ps_clean.rar.rich.df, Sample_type=="shell")))
# paste output here

summary(lm(Observed ~ Shell_Disease_Index, data=subset(Lobster_ps_clean.rar.rich.df, Sample_type=="shell")))
# paste output here

#  
install.packages("tidyverse")
install.packages("ggbeeswarm")
install.packages("gghalves")
library(tidyverse)
library(ggbeeswarm)
library(gghalves)
library(ggplot2)

Lobster_ps_clean.rar.rich.df$Timepoint <- factor(Lobster_ps_clean.rar.rich.df$Timepoint, levels=c("Baseline", "4_months", "10_months"), labels=c("Summer baseline", "Winter, 4 months", "Summer, 10 months")) #CHANGE ME

# FIGURE: Lobster_temp_richness_shelldisease
ggplot(data=subset(Lobster_ps_clean.rar.rich.df, !is.na(Shell_Disease_Index)), aes(x=Shell_Disease_Index, y=Observed))+
  theme_minimal() +
  theme_set(theme_minimal(base_size = 14)) + 
  facet_grid(~Timepoint, scales="free", space = "free") +
  geom_half_boxplot(aes(group = Shell_Disease_Index)) +
  geom_point(aes(color=Tank_temp_C, size=as.numeric(Shell_Stage), shape = Temp_geography), position = position_nudge(x=0.15)) +
  scale_color_viridis(discrete=TRUE, option="A", begin = 0, end =0.75) +
  ylab("Bacterial richness (SVs)") + xlab("Shell Disease Index") + labs(color = "Tank temperature (?C)", size = "Shell molting stage", shape = "Location", linetype="Location")


summary(lm(Observed ~ Shell_Stage, data=subset(Lobster_ps_clean.rar.rich.df, Sample_type=="shell")))
# paste output here


#FIGURE: Lobster_temp_richness_shellstageovertime
ggplot(data=subset(Lobster_ps_clean.rar.rich.df, !is.na(Sample_date)), aes(x=Sample_date, y=Shell_Stage))+
   theme_minimal() +
   theme_set(theme_minimal(base_size = 14)) + 
  #theme(legend.position = "bottom") +
  facet_grid(Temp_geography~., scales= "free", space= "free")+
    # geom_half_boxplot(aes(group = Sample_date)) +
  geom_point(aes(color=Tank_temp_C, size=as.numeric(Observed), shape = Shell_Disease_Index)) +
  geom_jitter() +
  scale_color_viridis(discrete=TRUE, option="A", begin = 0, end =0.75) +
  geom_smooth(aes(group= Shell_Disease_Index, linetype=Shell_Disease_Index), alpha = 1/6) +
  ylab("Shell molting stage") + xlab("Sample date") + labs(color = "Tank temperature (?C)", size = "Bacterial SV richness", shape = "Shell Disease Index", linetype="Shell Disease Index")


summary(lm(Observed ~ as.numeric(Weight_g), data=subset(Lobster_ps_clean.rar.rich.df, Sample_type=="shell")))
# paste output here


summary(lm(Observed ~ as.numeric(Length_mm), data=subset(Lobster_ps_clean.rar.rich.df, Sample_type=="shell")))
# paste output here


summary(lm(Observed ~ Tank, data=subset(Lobster_ps_clean.rar.rich.df, Sample_type=="shell")))
# paste output here

summary(lm(Observed ~ Temp_geography, data=subset(Lobster_ps_clean.rar.rich.df, Sample_type=="shell")))
# paste output here


summary(lm(Observed ~ Timepoint, data=subset(Lobster_ps_clean.rar.rich.df, Sample_type=="shell")))
# paste output here


Lobster_ps_clean.rar.rich.df$Timepoint <- factor(Lobster_ps_clean.rar.rich.df$Timepoint, levels=c("Baseline", "4_months", "10_months"), labels=c("Summer baseline", "Winter, 4 months", "Summer, 10 months")) #CHANGE ME

# FIGURE: Lobster_temp_richness_evenness_geographytime
plot1 <- ggplot(data=subset(Lobster_ps_clean.rar.rich.df, !is.na(Timepoint)), aes(x=Temp_geography, y=Observed))+
  theme_minimal() +
  theme_set(theme_minimal(base_size = 14)) + 
  facet_grid(.~Timepoint, space="free") +
  geom_half_boxplot(aes(group = Temp_geography)) +
  geom_point(aes(color=Tank_temp_C, size=as.numeric(Shell_Disease_Index), shape = Temp_geography), position = position_nudge(x=0.15)) +
  scale_color_viridis(discrete=TRUE, option="A", begin = 0, end =0.75) +
  ylab("Bacterial richness (SVs)") + xlab("") + labs(color = "Tank temperature (?C) ", size = "Shell Disease Index", shape = "Location")

# FIGURE: Lobster_temp_richness_evenness_geographytime
plot2 <- ggplot(data=subset(Lobster_ps_clean.rar.rich.df, !is.na(Timepoint)), aes(x=Temp_geography, y=Lobster_ps_clean.rar.even))+
  theme_minimal() +
  theme_set(theme_minimal(base_size = 14)) + 
  facet_grid(.~Timepoint, space="free") +
  geom_half_boxplot(aes(group = Temp_geography)) +
  geom_point(aes(color=Tank_temp_C, size=as.numeric(Shell_Disease_Index), shape = Temp_geography), position = position_nudge(x=0.15)) +
  scale_color_viridis(discrete=TRUE, option="A", begin = 0, end =0.75) +
  ylab("Bacterial evenness") + xlab("") + labs(color = "Tank temperature (?C) ", size = "Shell Disease Index", shape = "Location")

 library(ggpubr)
# plot together FIGURE: Lobster_temp_richness_evenness_geographytime
 ggarrange(plot1, plot2,
           ncol = 1, nrow=2, labels =c("A", "B"), common.legend = TRUE)


summary(lm(Observed ~ Temp_geography + (Temp_geography/Timepoint), data=subset(Lobster_ps_clean.rar.rich.df, Sample_type=="shell")))
# paste output here
# notes; this really compares everything to the northern ME baseline, but should be OK because all the baseline samples were the same


summary(lm(Observed ~ Temp_geography + (Temp_geography/Shell_Disease_Index), data=subset(Lobster_ps_clean.rar.rich.df, Sample_type=="shell")))
# is temp geography important when nested inside shell disease? i.e., does shell disease being more common in one location explain the effect?
# paste output here


summary(lm(Observed ~ Shell_Disease_Index + (Shell_Disease_Index/Temp_geography), data=subset(Lobster_ps_clean.rar.rich.df, Sample_type=="shell")))
# is shell disease important when nested inside geography?
# paste output here


# add/remove variables if they are significant.  be sure to drop the control tank samples
# cohort effect (1|Tank)
# Repeated measures over time (1/Lobster_ID|Tank)
glm <- (glmer(Observed ~ Temp_geography * Shell_Disease_Index * Shell_Stage + (1/Lobster_ID|Tank) + (1/Tank) + (Temp_geography/Timepoint), family=poisson, data=subset(Lobster_ps_clean.rar.rich.df, Sample_type=="shell")))

summary(glm)
# paste output here


## evenness calculations
summary(lm(Lobster_ps_clean.rar.even ~ as.numeric(Tank_temp_C), data=subset(Lobster_ps_clean.rar.rich.df, Sample_type=="shell")))
# paste output here

summary(lm(Lobster_ps_clean.rar.even ~ Tank_temp_C + (Tank_temp_C/Timepoint), data=subset(Lobster_ps_clean.rar.rich.df, Sample_type=="shell")))
# paste output here

summary(lm(Lobster_ps_clean.rar.even ~ Shell_Disease_Index, data=subset(Lobster_ps_clean.rar.rich.df, Sample_type=="shell")))
# paste output here

summary(lm(Lobster_ps_clean.rar.even ~ Temp_geography + (Temp_geography/Timepoint), data=subset(Lobster_ps_clean.rar.rich.df, Sample_type=="shell")))
# paste output here
# 

summary(lm(Lobster_ps_clean.rar.even ~ Temp_geography + (Temp_geography/Shell_Disease_Index), data=subset(Lobster_ps_clean.rar.rich.df, Sample_type=="shell")))
# paste output here


## ----- Heatmaps --------------
# base plot
plot_heatmap(Lobster_ps_clean.rar)

# follow the tutorial to make prettier versions in phyloseq: https://joey711.github.io/phyloseq/plot_heatmap-examples.html


library(phyloseq)
library(vegan)
require(plyr)
require(dplyr)
require(magrittr)
require(scales)
require(grid)
require(reshape2)
require(knitr)
library(ggplot2)
library(RColorBrewer)

#take out top 100 taxa based on abundance
topNOTUs = names(sort(taxa_sums(Lobster_ps_clean.rar), TRUE)[1:100])
top100 = prune_taxa(topNOTUs, Lobster_ps_clean.rar)

SV_table <- otu_table(top100)

# output phyloseq tax table as a dataframe to make it manipulatable
tax.df <- data.frame(tax_table(top100), stringsAsFactors=FALSE)

## if the Genus is empty, replace with the Family
tax.df$Genus[is.na(tax.df$Genus)] <- tax.df$Family[is.na(tax.df$Genus)]

colnames(SV_table) <- tax.df$Genus

# Combine response and SVs into data frame
SV_table.df <- data.frame(SV_table)

# add some factors that you can use to make your graph pretty, as many as you want
SV_table.df$Sample <- row.names(sample_data(top100)) #always grab the sample names
SV_table.df$Temp_geography <- sample_data(top100)$Temp_geography 
SV_table.df$Temp_C <- sample_data(top100)$Tank_temp_C
SV_table.df$Temp_regime <- sample_data(top100)$Temp_regime
SV_table.df$Timepoint <- sample_data(top100)$Timepoint
SV_table.df$Sample_date <- sample_data(top100)$Sample_date
SV_table.df$Shell_health <- sample_data(top100)$Shell_health

# sort the factors in the timepoint so they will happen in chronological order
SV_table.df$Timepoint <- factor(SV_table.df$Timepoint, levels=c("Baseline", "4_months", "10_months", "NA"), labels=c("Baseline", "4 months", "10 months", "NA"))


# melt and transform the data usign ALL the factors you added
m <- melt(SV_table.df, id.vars=c("Sample", "Temp_geography", "Temp_C", "Temp_regime", "Timepoint", "Sample_date", "Shell_health")) #CHANGE these 'factors' to be the names of the variables you added in the previous step.  add as many as you added above
m <- ddply(m, .(variable), transform, rescale = log(1 + value)) #NOTE: may need to change it to as.numeric(value) if it tells you an error about non-binary operators

saveRDS(m, "melted_temp_data_top100.RDS")

# set color palette  
col.bro <- (rainbow(6))

# add white to that color list
col.bro <- append(col.bro, "#ffffff", after = 6)

# adjusted plot for factorial data, change 'factor a' to be whatever factors in your data you want to accent
ggplot(data = na.omit(m), aes(as.factor(Timepoint), variable)) +
  theme_minimal() +
  facet_grid(.~Temp_geography, space="free", scale="free") + #set up graph facets to separate out levels of FactorA
  geom_tile(aes(fill = rescale), color="gray") + #add the heatmap coloring
  scale_fill_gradientn(colors = rev(col.bro), na.value = 'white') + #use the preset color palette
  labs(fill="Log relative abundance") + #rename legend heading
  theme(legend.position = 'bottom',
        axis.text.x = element_text(angle = 0, size = 8),
        axis.ticks.x = element_line(size = 2),
        axis.text.y = element_text(size = 6)) +
  ylab('Bacterial SVs') +
  xlab('Timepoint') 


# adjusted plot for factorial data, change 'factor a' to be whatever factors in your data you want to accent
ggplot(data = na.omit(m), aes(as.factor(Timepoint), variable)) +
  theme_minimal() +
  facet_grid(Shell_health~Temp_geography, space="free", scale="free") + #set up graph facets to separate out levels of FactorA
  geom_tile(aes(fill = rescale), color="gray") + #add the heatmap coloring
  scale_fill_gradientn(colors = rev(col.bro), na.value = 'white') + #use the preset color palette
  labs(fill="Log relative abundance") + #rename legend heading
  theme(legend.position = 'bottom',
        axis.text.x = element_text(angle = 0, size = 8),
        axis.ticks.x = element_line(size = 2),
        axis.text.y = element_text(size = 6)) +
  ylab('Bacterial SVs') +
  xlab('Timepoint') 


 
## Feature Prediction (Differential Abundance) with Random Forest --------------------------------
#  https://rpubs.com/michberr/randomforestmicrobe
# https://www.stat.berkeley.edu/~breiman/RandomForests/cc_home.htm
# https://www.rdocumentation.org/packages/randomForest/versions/4.6-12/topics/randomForest

library(phyloseq)
library(vegan)
library(plyr)
library(dplyr)
require(magrittr)
require(scales)
require(grid)
require(reshape2)
require(knitr)
library(randomForest)
library(rfPermute)
library(ggplot2)
library(RColorBrewer)

# Make a dataframe of training data with OTUs as column and samples as rows, which is the phyloseq OTU table


# subset to shell only
Lobster_temp_shell.rar <- subset_samples(Lobster_ps_clean.rar, Sample_type == "shell")
Lobster_temp_shell.rar <- prune_taxa(taxa_sums(Lobster_temp_shell.rar) > 0, Lobster_temp_shell.rar)

#Subset to only the shell samples from the first time point to assess disease
Lobster_ps_clean_shell_base.rar <- subset_samples(Lobster_ps_clean.rar, Sample_type == "shell"| Timepoint=="Baseline")
Lobster_ps_clean_shell_base.rar <- prune_taxa(taxa_sums(Lobster_ps_clean_shell_base.rar) > 0, Lobster_ps_clean_shell_base.rar)

Lobster_temp_shell_disease.rar <- subset_samples(Lobster_ps_clean.rar, Sample_type == "shell"& Shell_Disease_Index!=0)
Lobster_temp_shell_disease.rar <- prune_taxa(taxa_sums(Lobster_temp_shell_disease.rar) > 0, Lobster_temp_shell_disease.rar) # 101 samples and 4292 taxa to start with


predictors <- otu_table(Lobster_temp_shell.rar) # use only shell and not tank samples
# predictors <- otu_table(Lobster_ps_clean_shell_base.rar)
# predictors <- otu_table(Lobster_temp_shell_disease.rar)


dim(predictors)


# output phyloseq tax table as a dataframe to make it manipulable
tax.df <- data.frame(tax_table(Lobster_temp_shell.rar), stringsAsFactors=FALSE)
# tax.df <- data.frame(tax_table(Lobster_ps_clean_shell_base.rar), stringsAsFactors=FALSE)
# tax.df <- data.frame(tax_table(Lobster_temp_shell_disease.rar), stringsAsFactors=FALSE)

## if the Genus is empty, replace with the Family
tax.df$Genus[is.na(tax.df$Genus)] <- tax.df$Family[is.na(tax.df$Genus)]

# bind Genus and Species together
tax.df$Genus.species <- paste(tax.df$Genus, tax.df$Species)


# set column of combined genus and species names as the column names for the predictors, replacing the full SV
colnames(predictors) <- tax.df$Genus.species

# clean up some of the other taxa info
colnames(predictors) = gsub("_unclassified", "", colnames(predictors))
colnames(predictors) = gsub("_Intercertae_Sedis", "", colnames(predictors))
colnames(predictors) = gsub(" NA", "", colnames(predictors))

### start here when choosing factors, can reuse the above lines as needed. one example for factorial data, and one for numeric data is provided. select as needed.

# Make one column for our outcome/response variable. 
 response <- as.factor(sample_data(Lobster_temp_shell.rar)$Shell_Disease_Index) 
# response <- as.numeric(sample_data(Lobster_temp_shell.rar)$Tank_temp_C) 
# response <- as.factor(sample_data(Lobster_temp_shell.rar)$Shell_Stage)

# or, baseline only 
# response <- as.factor(sample_data(Lobster_temp_shell_base.rar)$Shell_Disease_Index) 

# or, get the mortality data as yes/no 
# sample_data(Lobster_temp_shell_disease.rar)$Mortality <- sample_data(Lobster_temp_shell_disease.rar)$Died
# sample_data(Lobster_temp_shell_disease.rar)$Mortality[is.na(sample_data(Lobster_temp_shell_disease.rar)$Mortality)] <- "No"  
# sample_data(Lobster_temp_shell_disease.rar)$Mortality <- as.character(sample_data(Lobster_temp_shell_disease.rar)$Mortality); sample_data(Lobster_temp_shell_disease.rar)$Mortality <- ifelse(sample_data(Lobster_temp_shell_disease.rar)$Mortality == "No", "No", "Yes")
# response <- as.factor(sample_data(Lobster_temp_shell_disease.rar)$Mortality) ## look at mortality

# Combine response and SVs into data frame
rf.data <- data.frame(response, predictors)


# set seed for random number generation reproducibility
set.seed(2)

# classify for factorial data
response.pf <- rfPermute(response ~. , data = rf.data, 
                         strata=sample_data(Lobster_temp_shell.rar)$Timepoint, # CHANGE ME
                       na.action = na.omit, ntree= 500, nrep = 100) #na.omit ignores NAs in data (not tolerated). ntrees is how many forests to build, nreps generates p-value


print(response.pf)
# paste the print out here, especially the OOB error. 1-(Out-of-the-box error) = accuracy of your model


# grab which features were labeled "important"
imp <- importance(response.pf, scale = TRUE)

# Make a data frame with predictor names and their importance
imp.df <- data.frame(predictors = rownames(imp), imp) 


# For factorial data, grab only those features with p-value < 0.05
imp.sig <- subset(imp.df, MeanDecreaseAccuracy.pval <= 0.05)
print(dim(imp.sig))

# or For numeric data, grab only those features with p-value < 0.05
imp.sig <- subset(imp.df, IncNodePurity.pval <= 0.05) 
print(dim(imp.sig))


# or For factorial data, sort by importance amount
imp.sort <- imp.sig[order(imp.sig$MeanDecreaseAccuracy),]

# For numeric data, sort by importance amount
imp.sort <- imp.sig[order(imp.sig$IncNodePurity),]
  


#create levels to the factor based on SV table
imp.sort$predictors <- factor(imp.sort$predictors, levels = imp.sort$predictors)

# saveRDS(imp.sort, "Lobster_temp_shelldisease_imp_RF.RDS")
# saveRDS(imp.sort, "Lobster_temp_geography_withintimepoint_imp_RF.RDS")
# saveRDS(imp.sort, "Lobster_temp_geography_base_disease_imp_RF.RDS")
# saveRDS(imp.sort, "Lobster_temp_temp_imp_RF.RDS") #redone
# saveRDS(imp.sort, "Lobster_temp_mortality_withintimepoint_imp_RF.RDS")
# saveRDS(imp.sort, "Lobster_temp_mortalitydiseased_withintimepoint_imp_RF.RDS")
# saveRDS(imp.sort, "Lobster_temp_shellmoltstage_imp_RF.RDS")

imp.sort<-readRDS("Lobster_temp_shelldisease_withintimepoint_imp_RF.RDS")

# Select the top so many predictors (more than 50 is a crowded graph)
imp.40 <- imp.sort[1:40, ]

# figure out what they are and name them, otherwise will just be named the full SV
otunames <- imp.40$predictors


# need to grab the abundance data out of the otu_table and make a new data.frame, and add the taxa names back in
# grab the column names from the otu table that match those in the forest set
pred.abun.colnum <- which(colnames(rf.data) %in% otunames)

# when you find a match, grad the abundance data
pred.abun <- rf.data[,sort(c(pred.abun.colnum))]

# make this into a datafram for manipulation
pred.abun.df <- data.frame(pred.abun, stringsAsFactors=FALSE)

# use the row.names (sample names) from the phyloseq object to name the samples in your forest
row.names(pred.abun.df) <- row.names(sample_data(Lobster_temp_shell.rar))

row.names(pred.abun.df) <- row.names(sample_data(Lobster_temp_shell_disease.rar))


# add some factors that you can use to make your graph pretty, as many as you want
pred.abun.df$Sample <- row.names(sample_data(Lobster_temp_shell.rar)) #always grab the sample names
pred.abun.df$Temp_geography <- sample_data(Lobster_temp_shell.rar)$Temp_geography #CHANGE ME
pred.abun.df$Timepoint <- sample_data(Lobster_temp_shell.rar)$Timepoint #CHANGE ME
pred.abun.df$Shell_Disease_Index <- sample_data(Lobster_temp_shell.rar)$Shell_Disease_Index
pred.abun.df$Tank_temp_C <- sample_data(Lobster_temp_shell.rar)$Tank_temp_C
pred.abun.df$Shell_Stage <- sample_data(Lobster_temp_shell.rar)$Shell_Stage
pred.abun.df$Mortality <- sample_data(Lobster_temp_shell.rar)$Mortality


# or for the shell disease subset, add some factors that you can use to make your graph pretty, as many as you want
pred.abun.df$Sample <- row.names(sample_data(Lobster_temp_shell_disease.rar)) #always grab the sample names
pred.abun.df$Temp_geography <- sample_data(Lobster_temp_shell_disease.rar)$Temp_geography #CHANGE ME
pred.abun.df$Timepoint <- sample_data(Lobster_temp_shell_disease.rar)$Timepoint #CHANGE ME
pred.abun.df$Shell_Disease_Index <- sample_data(Lobster_temp_shell_disease.rar)$Shell_Disease_Index
pred.abun.df$Tank_temp_C <- sample_data(Lobster_temp_shell_disease.rar)$Tank_temp_C
pred.abun.df$Shell_Stage <- sample_data(Lobster_temp_shell_disease.rar)$Shell_Stage
pred.abun.df$Mortality <- sample_data(Lobster_temp_shell_disease.rar)$Mortality

pred.abun.df$Timepoint <- factor(pred.abun.df$Timepoint, levels=c("Baseline", "4_months", "10_months"), labels=c("Summer baseline", "Winter, 4 months", "Summer, 10 months"))                                                                                                                                     

# melt and transform the data using ALL the factors you added
m <- melt(pred.abun.df, id.vars=c("Sample", "Temp_geography", "Timepoint", "Shell_Disease_Index", "Tank_temp_C", "Shell_Stage")) #Mortality
m <- ddply(m, .(variable), transform, rescale = log(1 + value))

library(viridis)

# FIGURE: random forest shell disease, 
ggplot(data=subset(m, !is.na(Shell_Disease_Index)), aes(Sample, variable)) + 
  theme_minimal() + 
  facet_grid(~Shell_Disease_Index, drop = TRUE, space="free", scales="free") + #set up graph facets to separate out levels of FactorA
  geom_tile(aes(fill = rescale)) + #add the heatmap coloring 
  scale_fill_viridis() + #use the preset color palette
  labs(fill="Log abundance") + #rename legend heading
  theme(legend.position = 'bottom',
        axis.text.x = element_blank(),
        axis.ticks.x = element_line(size = 2),
        axis.text.y = element_text(size = 8), 
        strip.text.x = element_text(size = 7)) +
  ylab('Predictor Taxa') +
  xlab('') 


# Figure Lobster_temp_RF_temp
ggplot(m, aes(Sample, variable)) + 
  theme_minimal() + 
  facet_grid(~Tank_temp_C, drop = TRUE, space="free", scales="free") + #set up graph facets to separate out levels of FactorA
  geom_tile(aes(fill = rescale)) + #add the heatmap coloring 
  scale_fill_viridis() + #use the preset color palette
  labs(fill="Log abundance") + #rename legend heading
  theme(legend.position = 'bottom',
        axis.text.x = element_blank(),
        axis.ticks.x = element_line(size = 2),
        axis.text.y = element_text(size = 8), 
        strip.text.x = element_text(size = 7)) +
  ylab('Predictor Taxa') +
  xlab('') 


# FIGURE: random forest mortality all 
ggplot(data=subset(m, !is.na(Shell_Disease_Index)), aes(Sample, variable)) + 
  theme_minimal() + 
  facet_grid(~Mortality*Shell_Disease_Index, drop = TRUE, space="free", scales="free") + #set up graph facets to separate out levels of FactorA
  geom_tile(aes(fill = rescale)) + #add the heatmap coloring 
  scale_fill_viridis() + #use the preset color palette
  labs(fill="Log abundance") + #rename legend heading
  theme(legend.position = 'bottom',
        axis.text.x = element_blank(),
        axis.ticks.x = element_line(size = 2),
        axis.text.y = element_text(size = 8), 
        strip.text.x = element_text(size = 7)) +
  ylab('Predictor Taxa') +
  xlab('') 

# FIGURE: random forest mortality only in shell disease
ggplot(data=subset(m, !is.na(Shell_Disease_Index)), aes(Sample, variable)) + 
  theme_minimal() + 
  facet_grid(~Mortality*Shell_Disease_Index, drop = TRUE, space="free", scales="free") + #set up graph facets to separate out levels of FactorA
  geom_tile(aes(fill = rescale)) + #add the heatmap coloring 
  scale_fill_viridis() + #use the preset color palette
  labs(fill="Log abundance") + #rename legend heading
  theme(legend.position = 'bottom',
        axis.text.x = element_blank(),
        axis.ticks.x = element_line(size = 2),
        axis.text.y = element_text(size = 8), 
        strip.text.x = element_text(size = 7)) +
  ylab('Predictor Taxa') +
  xlab('') 


# adjusted plot for shell disease at baseline 
ggplot(data=subset(m, !is.na(Shell_Disease_Index)), aes(Sample, variable)) + 
  theme_minimal() + 
  facet_grid(~Shell_Disease_Index, space="free", scale="free", labeller= labeller(.multi_line = TRUE)) + #set up graph facets to separate out levels of FactorA
  geom_tile(aes(fill = rescale)) + #add the heatmap coloring 
  scale_fill_viridis() + 
  labs(fill="Log abundance") + #rename legend heading
  theme(legend.position = 'bottom',
        axis.text.x = element_blank(),
        axis.ticks.x = element_line(size = 2),
        axis.text.y = element_text(size = 8)) +
  ylab('Predictor Taxa') +
  xlab('Shell Disease Index') 


# adjusted plot for temp on x axis
ggplot(m, aes(Sample, variable)) + 
  theme_minimal() + 
  facet_grid(.~Tank_temp_C, drop = TRUE, space="free", scales="free") + #set up graph facets to separate out levels of FactorA
  geom_tile(aes(fill = rescale)) + #add the heatmap coloring 
  scale_fill_viridis() + #use the preset color palette
  labs(fill="Log abundance") + #rename legend heading
  theme(legend.position = 'bottom',
        axis.text.x = element_blank(),
        axis.ticks.x = element_line(size = 2),
        axis.text.y = element_text(size = 8), 
        strip.text.x = element_text(size = 7)) +
  ylab('Predictor Taxa') +
  xlab('') 


# FIGURE: random  shell stage
ggplot(data=subset(m, !is.na(Shell_Stage)), aes(Sample, variable)) + 
  theme_minimal() + 
  facet_grid(~Shell_Stage, drop = TRUE, space="free", scales="free") + #set up graph facets to separate out levels of FactorA
  geom_tile(aes(fill = rescale)) + #add the heatmap coloring 
  scale_fill_viridis() + #use the preset color palette
  labs(fill="Log relative abundance") + #rename legend heading
  theme(legend.position = 'bottom',
        axis.text.x = element_blank(),
        axis.ticks.x = element_line(size = 2),
        axis.text.y = element_text(size = 8), 
        strip.text.x = element_text(size = 7)) +
  ylab('Predictor Taxa') +
  xlab('') 


## Beta diversity ordinations, PCoA in phyloseq  -----------------
# ordinations in phyloseq: https://joey711.github.io/phyloseq/plot_ordination-examples.html

EX.ordJ <- ordinate(Lobster_ps_clean.rar, #calculate similarities
                   method ="PCoA", #ordination type
                   "jaccard", binary = TRUE) #similarity type. Jaccard is binary, Bray can be binary (unweighted) or not (weighted)

#set order to some factor levels
sample_data(Lobster_ps_clean.rar)$Sample_type <- factor(sample_data(Lobster_ps_clean.rar)$Sample_type, levels=c("tank", "shell")) #CHANGE ME
sample_data(Lobster_ps_clean.rar)$Timepoint <- factor(sample_data(Lobster_ps_clean.rar)$Timepoint, levels=c("Baseline", "4_months", "10_months", "Control"), labels=c("Summer baseline", "Winter, 4 months", "Summer, 10 months", "Control"))                                                                                                                                     

library(viridis)

# FIGURE:  Lobster_temp_PCOA_BC_J_ellipses
plot1 <- plot_ordination(Lobster_ps_clean.rar, EX.ordJ, type="samples", color="Tank_temp_C", shape="Timepoint") +
  theme_minimal()+
  geom_point(aes(size = as.numeric(Shell_Disease_Index), alpha = Sample_type)) +
    scale_color_viridis(option="A", begin = 0, end = 0.75) + 
  stat_ellipse(aes(group=Temp_geography, linetype=Temp_geography)) +
  labs(color = "Tank temperature (?C)", size = "Shell Disease Index", shape = "Time point", linetype="Location", alpha="Sample type")


# Figure: ordination with Culture data

plot_ordination(Lobster_ps_clean.rar, EX.ordJ, type="samples", color="Tank_temp_C") +
  theme_minimal()+
  geom_point(aes(shape=Timepoint, alpha = Shell_Disease_Index, size = Marine_Per1cm2)) +
  scale_color_viridis(option="A", begin = 0, end = 0.75) + 
  stat_ellipse(aes(group=Temp_geography, linetype=Temp_geography)) +
  labs(color = "Tank temperature (?C)", size = "Marine media CFU per cm2", shape = "Time point", linetype="Location", alpha="Shell Disease Index")
# Summary: Marine CFU seemed more related to water temp than treatment, as more showed up in the winter and cool summer samples (11C) instead of following the hot temps, was not really associated with shell disease index in the context of whole community


EX.ord.bray <- ordinate(Lobster_ps_clean.rar, #calculate similarities
                   method ="PCoA", #ordination type
                   "bray", binary = FALSE) #similarity type. Jaccard is binary, Bray can be binary (unweighted) or not (weighted)


adonis(EX.ord_jac ~ Diet * Week, as(sample_data(EX_ps_clean.rar), "data.frame"), permutations=9999, na.action=na.omit)

plot2 <- plot_ordination(Lobster_ps_clean.rar, EX.ord.bray, type="samples", color="Tank_temp_C", shape="Timepoint") +
  theme_minimal() +
  geom_point(aes(size = as.numeric(Shell_Disease_Index), alpha = Sample_type)) +
  scale_color_viridis(option="A", begin = 0, end = 0.75) + 
 stat_ellipse(aes(group=Temp_geography, linetype=Temp_geography)) +
  labs(color = "Tank temperature (?C)", size = "Shell Disease Index", shape = "Time point", linetype="Location", alpha="Sample type")


library(ggpubr)
#FIGURE: Lobster_temp_PCOA_BC_J_ellipses
ggarrange(plot1, plot2,
          ncol = 1, nrow=2, labels =c("A", "B"), common.legend = TRUE, legend = "right")


library(vegan)

# subset to shell samples only
Lobster_temp_shell.rar <- subset_samples(Lobster_ps_clean.rar, Sample_type == "shell")

# ran through vegan directly because phyloseq to vegan formatting issues
lobster_J = phyloseq::distance(Lobster_temp_shell.rar, type="samples", "jaccard", binary=T) 

# add permutations for repreated measures
ctrl <- with(as.data.frame(sample_data(Lobster_temp_shell.rar)), how(blocks = Lobster_ID, nperm = 999))


adonis(lobster_J ~ Timepoint* Temp_geography * Shell_Disease_Index * Shell_Stage, data = data.frame(sample_data(Lobster_temp_shell.rar)), permutations=ctrl, na.action=na.omit)
# paste output here

adonis(lobster_J ~ Temp_geography * Shell_Disease_Index * Shell_Stage, data = data.frame(sample_data(Lobster_temp_shell.rar)), strata = sample_data(Lobster_temp_shell.rar)$Timepoint, permutations=ctrl, na.action=na.omit) # will run comparisons for samples within each tank grouping
# paste output here


# subset to shell and not summer baseline samples only
Lobster_temp_shell_exp.rar <- subset_samples(Lobster_ps_clean.rar, Sample_type == "shell" & Timepoint != "Summer baseline")

# ran through vegan directly because phyloseq to vegan formatting issues
lobster_exp_J = phyloseq::distance(Lobster_temp_shell_exp.rar, type="samples", "jaccard", binary=T) 

# add permutations for repreated measures
ctrl <- with(as.data.frame(sample_data(Lobster_temp_shell_exp.rar)), how(blocks = Lobster_ID, nperm = 999))

adonis(lobster_exp_J ~ Temp_geography * Shell_Disease_Index * Shell_Stage, data = data.frame(sample_data(Lobster_temp_shell_exp.rar)), strata = sample_data(Lobster_temp_shell_exp.rar)$Timepoint, permutations=ctrl, na.action=na.omit)
# paste output here


# subset to shell samples only
Lobster_temp_shell.rar <- subset_samples(Lobster_ps_clean.rar, Sample_type == "shell")

# ran through vegan directly because phyloseq to vegan formatting issues
lobster_B = phyloseq::distance(Lobster_temp_shell.rar, type="samples", "bray", binary=F) 

# add permutations for repreated measures
ctrl <- with(as.data.frame(sample_data(Lobster_temp_shell.rar)), how(blocks = Lobster_ID, nperm = 999))

adonis(lobster_B ~ Timepoint* Temp_geography * Shell_Disease_Index * Shell_Stage, data = data.frame(sample_data(Lobster_temp_shell.rar)), permutations=ctrl, na.action=na.omit)
# paste output here

adonis(lobster_B ~ Temp_geography * Shell_Disease_Index * Shell_Stage, data = data.frame(sample_data(Lobster_temp_shell.rar)), strata = sample_data(Lobster_temp_shell.rar)$Timepoint, permutations=ctrl, na.action=na.omit) # will run comparisons for samples within each tank grouping
# paste output here


# subset to shell and not summer baseline samples only
Lobster_temp_shell_exp.rar <- subset_samples(Lobster_ps_clean.rar, Sample_type == "shell" & Timepoint != "Summer baseline")

# ran through vegan directly because phyloseq to vegan formatting issues
lobster_exp_B = phyloseq::distance(Lobster_temp_shell_exp.rar, type="samples", "bray", binary=F) 

# add permutations for repreated measures
ctrl <- with(as.data.frame(sample_data(Lobster_temp_shell_exp.rar)), how(blocks = Lobster_ID, nperm = 999))

adonis(lobster_exp_B ~ Temp_geography * Shell_Disease_Index * Shell_Stage, data = data.frame(sample_data(Lobster_temp_shell_exp.rar)), strata = sample_data(Lobster_temp_shell_exp.rar)$Timepoint, permutations=ctrl, na.action=na.omit)

# paste output here


 

## Core community members -----------
#code from this website: https://microbiome.github.io/tutorials/CoremicrobiotaAmplicon.html

#load libraries as needed
library(dplyr)
BiocManager::install("microbiome")
library(microbiome)

# subset to shell samples only
Lobster_temp_shell.rar <- subset_samples(Lobster_ps_clean.rar, Sample_type == "shell")

# core for healthy
Lobster_temp_shell_healthy.rar = subset_samples(Lobster_temp_shell.rar, Shell_Disease_Index == "0") 
Lobster_temp_shell_healthy.rar = prune_taxa(taxa_sums(Lobster_temp_shell_healthy.rar) > 0, Lobster_temp_shell_healthy.rar) # 110 samples and 3808 taxa to start with

  # core by any ESD
  Lobster_temp_shell_ESD.rar = subset_samples(Lobster_temp_shell.rar, Shell_Disease_Index != "0") 
  Lobster_temp_shell_ESD.rar <- prune_taxa(taxa_sums(Lobster_temp_shell_ESD.rar) > 0, Lobster_temp_shell_ESD.rar) # 39 samples and 3099 taxa to start with
  

#calculate compositional version of the data (relative abundances). pick one to start with
core_biomeW.rel1 <- microbiome::transform(Lobster_temp_shell_healthy.rar, "compositional") #CHANGE ME to your phyloseq object
core_biomeW.rel2 <- microbiome::transform(Lobster_temp_shell_ESD.rar, "compositional") #CHANGE ME to your phyloseq object


#This returns the taxa that exceed the given prevalence and minimum abundance detection thresholds. Set your preferred thresholds.
core_taxa_standardW1 <- core_members(core_biomeW.rel1, detection = 1/1000, prevalence = 80/100) #CHANGE ME
core_taxa_standardW2 <- core_members(core_biomeW.rel2, detection = 1/1000, prevalence = 80/100) #CHANGE ME


#A full phyloseq object of the core microbiota at those limits is obtained as follows
phylo.coreW1 <- core(core_biomeW.rel1, detection = 1/1000, prevalence = .8)
phylo.coreW2 <- core(core_biomeW.rel2, detection = 1/1000, prevalence = .8)

###retrieving the associated taxa names from the phyloseq object and add it to what you just made.
core.taxaW1 <- taxa(phylo.coreW1)
class(core.taxaW1)

core.taxaW2 <- taxa(phylo.coreW2)
class(core.taxaW2)

# get the taxonomy data and assign it to what you just made.
tax.matW1 <- tax_table(phylo.coreW1)
tax.dfW1 <- as.data.frame(tax.matW1)

tax.matW2 <- tax_table(phylo.coreW2)
tax.dfW2 <- as.data.frame(tax.matW2)

# add the SVs to last column of what you just made.
tax.dfW1$SV <- rownames(tax.dfW1)
tax.dfW2$SV <- rownames(tax.dfW2)


dim(tax.dfW1)
# 4 8
## CHANGE ME, write down how many 7450 taxa are shared between these samples. The second number is the number of taxonomy columns you have.
dim(tax.dfW2)
#4 8


# select taxonomy of only those OTUs that are core members based on the thresholds that were used.
core.taxa.classW1 <- dplyr::filter(tax.dfW1, rownames(tax.dfW1) %in% core.taxaW1)
knitr::kable(head(core.taxa.classW1))

core.taxa.classW2 <- dplyr::filter(tax.dfW2, rownames(tax.dfW2) %in% core.taxaW2)
knitr::kable(head(core.taxa.classW2))

# save the list so you can access later, can report just the list
write.csv(core.taxa.classW1, "~/core.taxa.lobster_healthy.csv")
write.csv(core.taxa.classW2, "~/core.taxa.lobster_ESD.csv")


# how abundant are all the seqs from the core SVs put together?
core.abundance1 <- sample_sums(core(Lobster_temp_shell_healthy.rar, detection = .01, prevalence = .80))
core.abundance2 <- sample_sums(core(Lobster_temp_shell_ESD.rar, detection = .01, prevalence = .80))

# Or, graph the abundance of those shared taxa, here are some example: https://microbiome.github.io/tutorials/Core.html
#aggregate the genera so we don't get a lot of lines separating all the SVs
plot.gen1 <- aggregate_taxa(core_biomeW.rel1, "Genus")
plot.gen2 <- aggregate_taxa(core_biomeW.rel2, "Genus")

# load libraries as needed
library(ggplot2)
library(scales)
library(viridis)

prevalences <- seq(.05, 1, .05)
detections <- 10^seq(log10(1e-4), log10(.2), length = 5)

plot1 <- plot_core(plot.gen1, 
                plot.type = "heatmap", 
                prevalences = prevalences, 
                detections = detections, min.prevalence = .5) +
  xlab("Detection Threshold (Relative Abundance (%))") + ylab("Bacterial SVs shared by 80% of healthy lobster shells") +
  theme_minimal() + scale_fill_viridis() +
  scale_x_discrete(labels = c("1.0e-04", "6.7e-4", "4.5e-3","0.03", "0.2"))

plot2 <- plot_core(plot.gen2, 
                   plot.type = "heatmap", 
                   prevalences = prevalences, 
                   detections = detections, min.prevalence = .5) +
  xlab("Detection Threshold (Relative Abundance (%))") + ylab("Bacterial SVs shared by 80% of lobster shells with ESD") +
  theme_minimal() + scale_fill_viridis() +
  scale_x_discrete(labels = c("1.0e-04", "6.7e-4", "4.5e-3","0.03", "0.2"))

#Figure: Lobster_temp_core_healthyESD
# plot together
library(ggpubr)
ggarrange(plot1, plot2,
          ncol = 2, nrow=1, labels =c("A", "B"), common.legend = TRUE)


## Putative agents and culturing data ----------
# Aquimarina and Thalassobious species (Chistoserdov et al., 2012, Quinn et al., 2013, Quinn et al., 2017)
# top found by Debbie Bouchard et al.: Aquimarina, Chryseobacterium, Empedobacter, Flavobacterium, Polaribacter,  Pseudoaltermonas, Roseomonas,  Roseovarius, Sulfitobacter and Vibrio, but found in healthy and diseased. Thalossobius and Aquimarina from other sources

Lobster_temp_put = subset_taxa(Lobster_ps_clean.rar, Genus=="Aquimarina" | Genus=="Chryseobacterium" |Genus=="Empedobacter" |Genus=="Flavobacterium" |Genus=="Polaribacter" |Genus=="Pseudoaltermonas" |Genus=="Roseomonas" |Genus=="Roseovarius" |Genus=="Sulfitobacter" | Genus=="Vibrio" | Genus =="Thalossobius")
Lobster_temp_put = prune_taxa(taxa_sums(Lobster_temp_put) > 0, Lobster_temp_put)

Lobster_temp_put_glom <- tax_glom(Lobster_temp_put, "Genus")

#sample_data(Lobster_temp_put_glom)$Timepoint <- factor(sample_data(Lobster_temp_put_glom)$Timepoint, levels=c("Baseline", "4_months", "10_months"), labels=c("Summer baseline", "Winter, 4 months", "Summer, 10 months")) #CHANGE ME

# this version stacks all the genera together. Figure: Lobster_putative_stacked_glom
p <- plot_bar(Lobster_temp_put_glom, fill = "Genus") 
p + facet_grid(~Sample_type*Shell_Disease_Index, space="free", scales="free") + theme_minimal() + scale_fill_viridis(discrete=TRUE) + scale_color_viridis(discrete=TRUE) + geom_bar(aes(color=Genus, fill=Genus), stat="identity", position="stack") +  theme(axis.text.x = element_blank()) 


# this version is really messy to look at
p <- plot_bar(Lobster_temp_put_glom, fill = "Genus") 
p + facet_grid(Genus~Sample_type*Shell_Disease_Index, space="free", scales="free") + theme_minimal() + scale_fill_viridis(discrete=TRUE) + scale_color_viridis(discrete=TRUE) + geom_bar(aes(color=Genus, fill=Genus), stat="identity", position="stack") +  theme(axis.text.x = element_blank()) 

# this version is more selective, Figure: Lobster_aquimarina_sulfitobacter
Lobster_temp_put2 = subset_taxa(Lobster_ps_clean.rar, Genus=="Aquimarina" | Genus=="Sulfitobacter")

p <- plot_bar(Lobster_temp_put2, fill = "Genus") 
p + facet_grid(Genus~Sample_type*Shell_Disease_Index, space="free", scales="free") + theme_minimal() + scale_fill_viridis(discrete=TRUE) + scale_color_viridis(discrete=TRUE) + geom_bar(aes(color=Genus, fill=Genus), stat="identity", position="stack") +  theme(axis.text.x = element_blank()) 


# what about in mortality versus not? 

require(lubridate)
##### as.Date to import date into a usable format for continuous variable
#Lobster_ps_clean2 <- subset_samples(Lobster_ps_clean, !is.na(Sample_date))
sample_data(Lobster_temp_put)$Sample_date = as.Date(sample_data(Lobster_temp_put)$Sample_date, format = "%d-%b-%y")
sample_data(Lobster_temp_put)$Molt = as.Date(sample_data(Lobster_temp_put)$Molt, format = "%m/%d/%Y")
sample_data(Lobster_temp_put)$Died = as.Date(sample_data(Lobster_temp_put)$Died, format = "%m/%d/%Y")
                                                  
 shell_put <- subset_samples(Lobster_temp_put, Sample_type == "shell")                                                 
p <- plot_bar(shell_put, x="Sample_date", fill = "Genus") + facet_grid(Temp_geography*Tank~., space="free") + theme_minimal() + scale_fill_viridis(discrete=TRUE) + scale_color_viridis(discrete=TRUE) + geom_bar(aes(color=Genus, fill=Genus), stat="identity", position="stack") + geom_vline(aes(xintercept=Molt), size=1 ) + geom_vline(aes(xintercept=Died), linetype = "longdash", size=2) + xlab("Sample Date") 


tank_put = subset_samples(Lobster_temp_put, Sample_type == "tank")

p2 <- plot_bar(tank_put, x="Tank", fill = "Genus") + theme_minimal() + scale_fill_viridis(discrete=TRUE) + scale_color_viridis(discrete=TRUE) + geom_bar(aes(color=Genus, fill=Genus), stat="identity", position="stack") 

library(ggpubr)
# plot together
ggarrange(p, p2,
          ncol = 1, nrow=2, heights = c(4, 1), labels =c("A", "B"), common.legend = TRUE)


# do healthy or diseased shells look more like tank controls?
# Summary: visually, not really, there are very few of these in the tank, and much less in shells over time. There are molts in every tanks so hard to tell. maybe if graph by lobster?

p3 <- plot_bar(shell_put, x="Sample_date", fill = "Genus") + facet_grid(Shell_Disease_Index~., space="free", labeller = label_both) + theme_minimal() + scale_fill_viridis(discrete=TRUE) + scale_color_viridis(discrete=TRUE) + geom_bar(aes(color=Genus, fill=Genus), stat="identity", position="stack") + geom_vline(aes(xintercept=Molt), size=1 ) + geom_vline(aes(xintercept=Died), linetype = "longdash", size=2) + xlab("Sample Date") 


# input raw culture data 

culture <- read.csv('lobster_cultured_isolates_data.csv', header = T)
# 14 variables, 724 samples/isolates

require(lubridate)
##### as.Date to import date into a usable format for continuous variable

culture$Swab_sample_date = as.Date(culture$Swab_sample_date, format = "%d-%b-%y")

culture$Timepoint <- factor(culture$Timepoint, levels = c("Baseline_summer", "4_months_winter", "10_months_summer"), labels=c("Baseline, summer", "4 months, winter","10 months, summer")) #CHANGE ME


library(RColorBrewer)
library(ggplot2)
ggplot(culture, aes (x = Shell_disease_index)) + 
  geom_bar(aes(fill = Isolate_genus)) +
  facet_grid(Geographic_location~Timepoint,  space = "free") + theme(legend.position = "bottom") + xlab('Shell Disease Index') + ylab('Isolate count') + labs(fill ="Isolate GenBank match") 


library(reshape)
library(plyr)
library(dplyr)
# melt and transform the data usign ALL the factors you added
culture_sub <- subset(culture, select=c("Swab_sample_date","Timepoint",	"Tank",	"Geographic_location",	"Lobster_ID",	"Mortality", "Shell_disease_index", "Isolate_genus_species"))

culture_sub <- culture_sub %>% add_count(Isolate_genus_species)

# set color palette  
col.bro <- (rainbow(6))


ggplot(culture_sub, aes(Shell_disease_index, Isolate_genus_species)) + 
  facet_grid(~Geographic_location*Timepoint,  space = "free") +
  geom_tile(aes(fill=n)) +
  scale_fill_gradientn(colors = rev(col.bro), na.value = 'white') + #use the preset color palette
   theme(legend.position = "bottom") + xlab('Shell Disease Index') + ylab('Isolate GenBank match') + labs(fill ="Isolate counts")

# note: no visual trend by location or shell disease, although slightly more in the warmer tanks


# supp figures Fig S5 Lobster_isolates_lived_NME culture data lobsters that lived by location and time

ggplot(subset(culture_sub, Mortality == "no" & Isolate_genus_species != "n/a" & Geographic_location == "NorthernME"),  aes(as.factor(Lobster_ID), Isolate_genus_species)) + 
  facet_grid(.~Timepoint*Shell_disease_index,  space = "free", scale= "free") +
  geom_tile(aes(fill=n)) +
  scale_fill_viridis(discrete = FALSE) + #use the preset color palette
  theme(legend.position = "bottom", axis.text.x = element_blank(), strip.text.x = element_text(angle = 90), strip.text.y = element_text(angle = 0)) + xlab('Lobsters which survived in Northern ME tanks') + ylab('Isolate GenBank match') + labs(fill ="Isolate counts") 

# Fig S6 Lobster_isolates_lived_SME
ggplot(subset(culture_sub, Mortality == "no" & Isolate_genus_species != "n/a" & Geographic_location == "SouthernME"),  aes(as.factor(Lobster_ID), Isolate_genus_species)) + 
  facet_grid(.~Timepoint*Shell_disease_index,  space = "free", scale= "free") +
  geom_tile(aes(fill=n)) +
  scale_fill_viridis(discrete = FALSE) + #use the preset color palette
  theme(legend.position = "bottom", axis.text.x = element_blank(), strip.text.x = element_text(angle = 90), strip.text.y = element_text(angle = 0)) + xlab('Lobsters which survived in Southern ME tanks') + ylab('Isolate GenBank match') + labs(fill ="Isolate counts") 

# Fig S7 Lobster_isolates_lived_SNE
ggplot(subset(culture_sub, Mortality == "no" & Isolate_genus_species != "n/a" & Geographic_location == "SouthernNE"),  aes(as.factor(Lobster_ID), Isolate_genus_species)) + 
  facet_grid(.~Timepoint*Shell_disease_index,  space = "free", scale= "free") +
  geom_tile(aes(fill=n)) +
  scale_fill_viridis(discrete = FALSE) + #use the preset color palette
  theme(legend.position = "bottom", axis.text.x = element_blank(), strip.text.x = element_text(angle = 90), strip.text.y = element_text(angle = 0)) + xlab('Lobsters which survived in Southern NE tanks') + ylab('Isolate GenBank match') + labs(fill ="Isolate counts") 


# Figure 10 culture data lobsters which died by location and time
ggplot(subset(culture_sub, Mortality == "yes" & Isolate_genus_species != "n/a"),  aes(as.factor(Lobster_ID), Isolate_genus_species)) + 
  facet_grid(Geographic_location~Timepoint*Shell_disease_index,  space = "free", scale= "free") +
  geom_tile(aes(fill=n)) +
  scale_fill_viridis(discrete = FALSE) + #use the preset color palette
  theme(legend.position = "bottom", axis.text.x = element_blank(), strip.text.x = element_text(angle = 90), strip.text.y = element_text(angle = 0)) + xlab('Lobsters which died') + ylab('Isolate GenBank match') + labs(fill ="Isolate counts") 


# mortality and culturing, not used, was replaced by the previous graph
# lobsters who died: 6876, 6825, 6838, 6839, 6769, 6871, 6873, 6875, 6888, 6820, 6763, 6872, 6826, 6824, 6844

# melt and transform the data usign ALL the factors you added
# culture_sub <- subset(culture, select=c("Swab_sample_date","Timepoint",	"Tank",	"Geographic_location",	"Lobster_ID",	"Shell_disease_index", "Isolate_genus_species"))

# culture_sub_died <- culture_sub %>% # CHANGE ME to reflect the phyloseq object name you have
  filter(Lobster_ID == "6876" | Lobster_ID == "6825" | Lobster_ID ==  "6838" | Lobster_ID == "6839" | Lobster_ID == "6769"| Lobster_ID == "6871" | Lobster_ID == "6873" | Lobster_ID == "6875" | Lobster_ID == "6888" | Lobster_ID == "6820" | Lobster_ID == "6763" | Lobster_ID == "6872" | Lobster_ID ==  "6826" | Lobster_ID == "6824" | Lobster_ID == "6844")


# ggplot(culture_sub_died, aes(Shell_disease_index, Isolate_genus_species)) + 
  facet_grid(~Geographic_location*Timepoint,  space = "free") +
  geom_tile(aes(fill=n)) +
  scale_fill_gradientn(colors = rev(col.bro), na.value = 'white') + #use the preset color palette
  theme(legend.position = "bottom") + xlab('Shell Disease Index') + ylab('Isolate GenBank match') + labs(fill ="Isolate counts")


# culture_sub_lived <- culture_sub %>% # CHANGE ME to reflect the phyloseq object name you have
  filter(Lobster_ID != "6876" & Lobster_ID != "6825" & Lobster_ID !=  "6838" & Lobster_ID != "6839" & Lobster_ID != "6769"& Lobster_ID != "6871" & Lobster_ID != "6873" & Lobster_ID != "6875" & Lobster_ID != "6888" & Lobster_ID != "6820" & Lobster_ID != "6763" & Lobster_ID != "6872" & Lobster_ID !=  "6826" & Lobster_ID != "6824" & Lobster_ID != "6844")


# ggplot(culture_sub_lived, aes(Shell_disease_index, Isolate_genus_species)) + 
  facet_grid(~Geographic_location*Timepoint,  space = "free", scales = "free") +
  geom_tile(aes(fill=n)) +
  scale_fill_gradientn(colors = rev(col.bro), na.value = 'white') + #use the preset color palette
  theme(legend.position = "bottom") + xlab('Shell Disease Index') + ylab('Isolate GenBank match') + labs(fill ="Isolate counts")


# repeat that graph with the 16S data 

culture_ps <- subset_taxa(Lobster_ps_clean.rar, Genus == "Achromobacter" | Genus == "Achromobacter-Vibrio" | Genus == "Acidovorax" | Genus == "Acinetobacter" | Genus == "Aeromonas" | Genus == "Ahrensia" | Genus == "Alcanivorax" | Genus == "Altermonas" | Genus == "Aquaspirillum" | Genus == "Aquimarina" | Genus == "Brevibacterium" | Genus == "Brevundimonas" | Genus == "Brochothrix" | Genus == "Burkholderia" | Genus == "CDC" | Genus == "CDC_group_II" | Genus == "CDC_group_II-E_subgroup_A" | Genus == "CDC_group_II-H" | Genus == "Chromobacterium" | Genus == "Chryseobacterium" | Genus == "Colwellia" | Genus == "Comamonas" | Genus == "Corynebacterium" | Genus == "Curtobacterium" | Genus == "Deinococcus" | Genus == "Empedobacter" | Genus == "Flavobacteriaceae" | Genus == "Flavobacterium" | Genus == "Formosa" | Genus == "Formosa-Tenacibaculum" | Genus == "Gamma_Proteobacteria" | Genus == "Gordonia" | Genus == "Halomonas" | Genus == "Kiloniella" | Genus == "Kytococcus" | Genus == "Mesonia" | Genus == "Microbacterium" | Genus == "Micrococcus" | Genus == "Pantoea" | Genus == "Pasteurella" | Genus == "Phaeobacter" | Genus == "Photobacterium" | Genus == "Polaribacter" | Genus == "Pseudoalteromonas" | Genus == "Pseudomonas" | Genus == "Psychrobacter" | Genus == "Rhodococcus" | Genus == "Rhodococcus/Pseudoalteromonas" | Genus == "Roseomonas" | Genus == "Roseovarius" | Genus == "Ruegeria" | Genus == "Shewanella" | Genus == "Sphingobium" | Genus == "Sphingomonas" | Genus == "Sphingopyxis" | Genus == "Staphylococcus" | Genus == "Sulfitobacter" | Genus == "Tsukamurella" | Genus == "Vibrio" | Genus == "Winogradskyella" | Genus == "Xanthomonas" | Genus == "Zobellia")

# 155 samples and 243 taxa


# get the mortality data as yes/no 
sample_data(culture_ps)$Mortality <- sample_data(culture_ps)$Died
sample_data(culture_ps)$Mortality[is.na(sample_data(culture_ps)$Mortality)] <- "No"  
sample_data(culture_ps)$Mortality <- as.character(sample_data(culture_ps)$Mortality); sample_data(culture_ps)$Mortality <- ifelse(sample_data(culture_ps)$Mortality == "No", "No", "Yes")

culture_ps.df <- as.data.frame(otu_table(culture_ps))


# output phyloseq tax table as a dataframe to make it manipulatable
tax.df <- data.frame(tax_table(culture_ps), stringsAsFactors=FALSE)

## if the Genus is empty, replace with the Family
tax.df$Genus[is.na(tax.df$Genus)] <- tax.df$Family[is.na(tax.df$Genus)]

# bind Genus and Species together
tax.df$Genus.species <- paste(tax.df$Genus, tax.df$Species)

# set column of combined genus and species names as the column names for the predictors, replacing the full SV
colnames(culture_ps.df) <- tax.df$Genus.species

# clean up some of the other taxa info
colnames(culture_ps.df) = gsub("_unclassified", "", colnames(culture_ps.df))
colnames(culture_ps.df) = gsub("_Intercertae_Sedis", "", colnames(culture_ps.df))
colnames(culture_ps.df) = gsub(" NA", "", colnames(culture_ps.df))


# culture_ps.df <- culture_ps.df[ , order(names(culture_ps.df))]


# add some factors that you can use to make your graph pretty, as many as you want
culture_ps.df$Sample <- row.names(sample_data(culture_ps)) #always grab the sample names
culture_ps.df$Temp_geography <- sample_data(culture_ps)$Temp_geography #CHANGE ME
culture_ps.df$Timepoint <- sample_data(culture_ps)$Timepoint #CHANGE ME
culture_ps.df$Shell_Disease_Index <- sample_data(culture_ps)$Shell_Disease_Index
culture_ps.df$Tank_temp_C <- sample_data(culture_ps)$Tank_temp_C
culture_ps.df$Mortality <- sample_data(culture_ps)$Mortality


# melt and transform the data usign ALL the factors you added
m <- melt(culture_ps.df, id.vars=c("Sample", "Temp_geography", "Timepoint", "Shell_Disease_Index", "Tank_temp_C", "Mortality")) #CHANGE these 'factors' to be the names of the variables you added in the previous step.  add as many as you added above

m <- ddply(m, .(variable), transform, rescale = log(1 + value))


ggplot(m, aes(Shell_Disease_Index, variable)) + 
  facet_grid(~Mortality,  space = "free", scales="free") +
  geom_tile(aes(fill = rescale)) +
  scale_fill_gradientn(colors = rev(col.bro), na.value = 'white') + #use the preset color palette
  theme(legend.position = "bottom", axis.text.x = element_blank()) + xlab('Shell Disease Index') + ylab('Bacterial SV') + labs(fill ="Log Sequence Counts") 


## Lobster health and factors -------
summary(lm(Shell_Disease_Index ~ Tank_temp_C, data=subset(Lobster_ps_clean.rar.rich.df, Sample_type=="shell")))
# paste output here

summary(lm(Shell_Disease_Index ~ Shell_Stage, data=subset(Lobster_ps_clean.rar.rich.df, Sample_type=="shell")))
# paste output here

summary(lm(Shell_Disease_Index ~ Timepoint, data=subset(Lobster_ps_clean.rar.rich.df, Sample_type=="shell")))
# paste output here

summary(lm(Shell_Disease_Index ~ Temp_geography, data=subset(Lobster_ps_clean.rar.rich.df, Sample_type=="shell")))
# paste output here

summary(lm(Shell_Disease_Index ~ Temp_geography * Timepoint, data=subset(Lobster_ps_clean.rar.rich.df, Sample_type=="shell")))
# paste output here

# "Molt"                      "Died"                
summary(lm(Shell_Disease_Index ~ as.numeric(BAwNaCL_Per1cm2), data=subset(Lobster_ps_clean.rar.rich.df, Sample_type=="shell")))
# paste output here

summary(lm(Shell_Disease_Index ~ as.numeric(Marine_Per1cm2), data=subset(Lobster_ps_clean.rar.rich.df, Sample_type=="shell")))
# paste output here

# does index change by death yes/no
summary(lm(Shell_Disease_Index ~ as.Date(Died), data=subset(Lobster_ps_clean.rar.rich.df, Sample_type=="shell")))
# paste output here

summary(lm(Observed ~ Shell_Disease_Index * as.Date(Died), data=subset(Lobster_ps_clean.rar.rich.df, Sample_type=="shell")))
# paste output here

# does richness or evenness change by death yes/no
summary(lm(Observed ~ as.Date(Died), data=subset(Lobster_ps_clean.rar.rich.df, Sample_type=="shell")))
# paste output here

Lobster_ps_clean.rar.rich.df$Died <- addNA(Lobster_ps_clean.rar.rich.df$Died)

ggplot(Lobster_ps_clean.rar.rich.df, aes(x=as.Date(Sample_date), y = Observed, color = Tank_temp_C)) + geom_point() + facet_grid(.~Died)

# Summary, after visualizing that corfreclation, this is significant but only because richness is decreasing over time anyway


seqtab.nochim <- otu_table(Lobster_ps_clean)
all.taxa.sp <- tax_table(Lobster_ps_clean)

# load seqtab as needed
seqtab.nochim <- readRDS('~/JV31/JV31seqtab.nochim_lobster.rds') #CHANGE ME

## load taxa table as needed
all.taxa.sp <- read.csv('~/JV31/taxa_species_silva138_JV31.csv', header=T, row.names=1) #CHANGE ME

# load metadata table as needed
meta <- read.csv("~/R/Lobster_metadata.csv", header=T, row.names=1) #CHANGE ME

## create a phyloseq object with all samples (subset out later)
Lobster_ps <- phyloseq(otu_table(seqtab.nochim, taxa_are_rows=FALSE), 
                       sample_data(meta), 
                       tax_table(as.matrix(all.taxa.sp)))
